# Genome sequencing analysis identifies high-risk Epstein-Barr virus subtypes for nasopharyngeal carcinoma

**DOI:** 10.1101/486613

**Authors:** Miao Xu, Youyuan Yao, Hui Chen, Shanshan Zhang, Tong Xiang, Su-Mei Cao, Zhe Zhang, Bing Luo, Zhiwei Liu, Zilin Li, Guiping He, Qi-Sheng Feng, Li-Zhen Chen, Xiang Guo, Weihua Jia, Ming-Yuan Chen, Bingchun Zhao, Xiao Zhang, Shang-Hang Xie, Roujun Peng, Ellen T. Chang, Vincent Pedergnana, Lin Feng, Jin-Xin Bei, Ruihua Xu, Mu Sheng Zeng, Weimin Ye, Hans-Olov Adami, Xihong Lin, Weiwei Zhai, Yi-Xin Zeng, Jianjun Liu

**Affiliations:** State Key Laboratory of Oncology in South China, Collaborative Innovation Center for Cancer Medicine, Sun Yat-sen University Cancer Center, Guangzhou, China; Human Genetics, Genome Institute of Singapore, Agency for Science, Technology and Research (A*STAR), Singapore; Department of Otolaryngology/Head and Neck Surgery, First Affiliated Hospital of Guangxi Medical University, Nanning, China; Department of Medical Microbiology, Qingdao University Medical College, Qingdao, China; Infections and Immunoepidemiology Branch, National Cancer Institute, Rockville, MD, USA; Department of Biostatistics, Harvard T. H. Chan School of Public Health, Boston, MA, USA; Division of Epidemiology, Department of Health Research and Policy, Stanford University School of Medicine, Stanford, CA, USA; Department of Medical Epidemiology and Biostatistics, Karolinska Institutet, Stockholm, Sweden; Key Laboratory of Zoological Systematics and Evolution, Institute of Zoology, Chinese Academy of Sciences, Beijing, China; Center for Excellence in Animal Evolution and Genetics, Chinese Academy of Sciences, Kunming, China; Department of Comprehensive Medical Oncology, Key Laboratory of Head & Neck Cancer Translational Research of Zhejiang Province, Zhejiang Cancer Hospital, Hangzhou, China

## Abstract

Epstein-Barr virus (EBV) infection is ubiquitous worldwide and associated with multiple cancers including nasopharyngeal carcinoma (NPC). The role of EBV viral genomic variation in NPC development and its striking endemicity in southern China has been poorly explored. Through large-scale genome sequencing and association study of EBV isolates from China, we identified two non-synonymous EBV variants within *BALF2* strongly associated with NPC risk (conditional *P* value 1.75×10^-6^ for SNP162476_C and 3.23×10^-13^ for SNP163364_T), whose cumulative effects contributed to 83% of the overall risk in southern China. Phylogenetic analysis of the risk variants revealed a unique origin in southern China followed by clonal expansion. EBV *BALF2* haplotype carrying the risk variants were shown to reduce viral lytic DNA replication, as a result potentially promoting viral latency. Our discovery has not only provided insight to the unique endemic pattern of NPC occurrence in southern China, but also paved the way for the identification of individuals at high risk of NPC and effective intervention program to reduce the disease burden in southern China.

## Introduction

Epstein-Barr virus (EBV) was discovered in 1964^1,2^, and has been the first human virus to be associated with cancers, including nasopharyngeal carcinoma (NPC), a subset of gastric carcinoma, and several kinds of lymphomas^3^. Although EBV infection is ubiquitous in human populations worldwide, its most closely associated malignancy, NPC, has unique geographic distribution. NPC is rare in most part of the world, but very common in southern China where the incidence rate can reach 20 - 40 per 100,000 individuals per year^4^. Multiple human susceptibility loci, including *HLA*, *CDKN2A/2B*, *TNFRSF19*, *MECOM*, and *TERT* loci, have been discovered for NPC, but these loci have limited contribution to overall risk^5–8^. Moreover, the risk variants of these loci distributed widely in Chinese and could not explain the unique endemics of NPC in southern China. Understanding the etiology of NPC, commonly known as the Cantonese cancer, remains enigmatic.

Since the first EBV genome sequence, B95-8, was published in 1984^9^, more than a hundred EBV genomes have been sequenced in spontaneous lymphoblastoid cell lines and patients diagnosed with EBV-associated diseases, which revealed significant genomic variation among EBV isolates from different geographic origins^10–15^. Even though the role of EBV genome variation in the risk of EBV-associated diseases has been explored^15–17^, previous studies suffered from the confounding effect of geographic distribution and insufficient sample size. As a result, robust epidemiological and genetic evidence linking specific EBV strains to the pathogenesis of NPC is yet to be uncovered.

In the current study, we performed large-scale EBV whole-genome sequencing (WGS) study of 215 EBV isolates from patients diagnosed with EBV-associated cancers (including NPC, gastric carcinoma, and lymphomas) and 54 isolates from healthy controls that were recruited from both NPC-endemic and non-endemic regions of China. Through a comprehensive and systematic association analysis of EBV genomic variation and subsequent replication analysis in an independent sample, we identified two high-risk EBV variants for NPC. Functional investigation uncovered their decreased ability to activate EBV lytic DNA replication which potentially promotes EBV latency and NPC tumorigenesis. In addition, phylogenetic analysis of the isolates from the current study and worldwide strains suggested a unique evolutionary origin of the two NPC-high-risk variants in southern China. For the first time, we have uncovered the high-risk EBV subtypes that contributed significantly to the overall risk of NPC, as well as its unique endemics in southern China.

### EBV whole-genome sequencing

Using a capture-based WGS protocol, we obtained EBV genome sequences from 215 tumor tissue, saliva and plasma samples of EBV-associated cancer patients (NPC, gastric carcinoma, and lymphomas) and 54 saliva samples of healthy donors, as well as one from NPC cell line C666.1 (For details, see **Supplementary Table 1** and **Methods**). Of the 269 EBV isolates, 220 were obtained from the NPC-endemic region of southern China (Guangdong and Guangxi Provinces), and 49 were from NPC-non-endemic regions of China. The average sequencing depth of all the isolates was 1,282×, and on average 95% of EBV genome was covered with at least 10× reads (**Supplementary Fig. 1**). Using B95-8 as the reference, we identified a total of 8,469 variants (8015 SNPs, 454 INDELs) across the EBV genome (for variant statistics, see **Supplementary Table 2** and **Supplementary Fig. 1**). The number of variants identified in each sample ranged from 1,006 to 2,104, with EBNAs and LMPs genes being the most polymorphic genes (**Supplementary Fig. 2a, b**), consistent with prior reports^14–16^.

To explore the accuracy in sequencing and variant calling, we compared the re-sequenced C666-1 EBV genome against the previous published record, and a high concordance rate of 97.9% was found^18^ (**Supplementary Table 3**). In addition, when subsets of variants discovered by WGS were re-genotyped by Sanger sequencing and MassArray iPLEX assay; 97.55% and 99.99% of tested variants were confirmed, respectively (**Supplementary Tables 4** and **5**). Both of these evidences suggest a very high accuracy in our sequencing and variant calling.

In order to understand intra-host polymorphism within an individual, two EBV fragments were amplified and sequenced in 25 paired saliva and tumor samples from NPC cases. The variant dissimilarity between the paired saliva and tumor samples (median 1.1%, 1^st^ to 3^rd^ quartile: 0-3.4%) was substantially lower than the between-host dissimilarity (median 13.5%, 1^st^ to 3^rd^ quartile: 3.7-16.9%) (**Supplementary Fig. 3)**. In addition, we sequenced the EBV whole genomes from the same NPC patient in paired tumor and saliva samples, and we observed that 99.27% of the variants were concordant between the EBV tumor and saliva isolates (**Supplementary Table 6**). Taken together, these observations suggested that paired saliva and tumor samples from the same subject contained the same EBV genome or strain. Therefore, we combined the genome sequence information from tumor tissue and saliva samples from NPC cases in subsequent analyses.

### Identification of high-risk EBV variants by two-stage genome-wide association analysis

To investigate the impact of EBV genomic variations on NPC risk, we performed a two-stage genome-wide association study of the EBV genome. The genome-wide discovery analysis was performed by testing 1545 EBV variants in 156 NPC cases and 47 healthy controls from Guangdong and Guangxi Provinces in the NPC-endemic region of Southern China (**Discovery phase**). A principal component analysis (PCA) of the human genome variation of all the cases and controls with the reference population samples from the 1000G project^19^ confirmed the Chinese origin of and genetic match between the cases and controls (**Supplementary Fig. 4a, b**). We also performed a PCA of EBV viral genome variation by using all the strains and C666-1 genome sequence from the current study with 97 publicly available genomes. We observed a continuous distribution of the EBV strains along the first principal component (PC) ranging from Africa and Europe to Asia (**Fig. 1a**). Within Asia, the second PC showed a partial separation of the isolates from NPC-endemic region and the ones from the non-endemic region of Asia (**Fig. 1a, d**).

**Figure 1.**
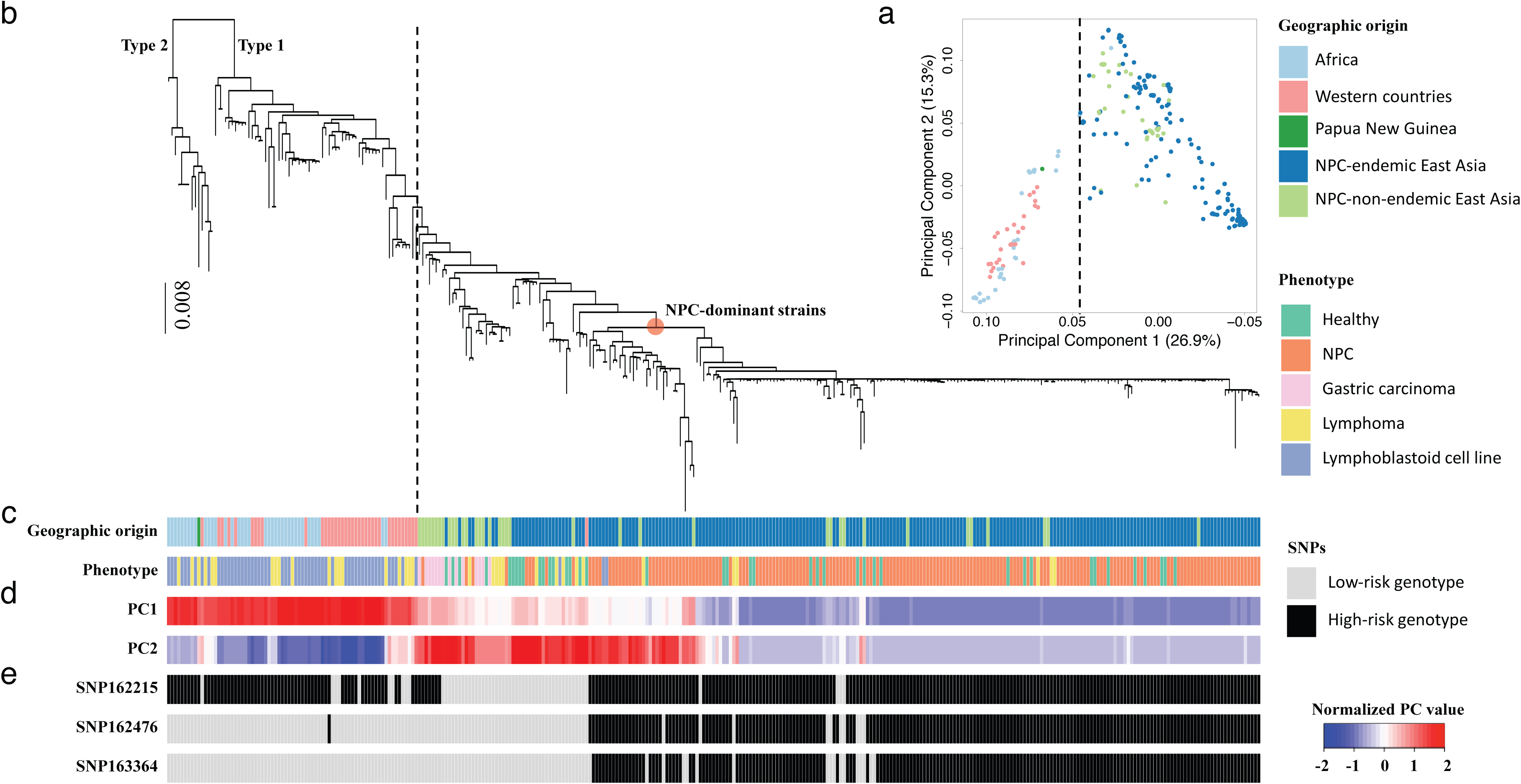
Principal component and phylogenetic analyses of EBV genomes. (**a**) Principal component analysis of 270 EBV isolates sequenced in current study and 97 published isolates. PC1 and PC2 scores are shown. Explaining 26.9% of the total genomic variance, PC1 discriminates between East Asian and Western/African strains. PC2 explains 15.3% of the total variance. (**b**) Phylogeny of 230 EBV single strains sequenced in current study and 97 published strains. Macacine herpesvirus 4 genome sequence (NC_006146) was used as the outgroup to root the tree. (**c**) Geographical origins and phenotypes of samples from which EBV strains were sequenced are shown with colors as indicated. (**d**) The normalized values of the first two principal-component scores (PC1 and PC2) are shown by colors from blue to red. (**e**) The genotypes of SNPs 162215, 162476 and 163364 in each isolate. Dashed lines in (a) and (b) indicate the separation between East Asian and Western/African strains. Red dot on the phylogeny indicates the lineage of NPC-dominant EBV strains, where 22 of 37 strains from healthy controls from NPC-endemic southern China were located.

To control for the potential impact of the population structures of both the human and EBV genomes, the genome-wide discovery association analysis was performed using generalized-linear mixed model with age, sex, the first four human PCs as fixed effect and the genetic relatedness matrix of EBV genomes as random effect^20^. The discovery analysis revealed multiple association signals along the EBV genome, with the strongest association observed within the *BALF2* region (SNP162507, *P* = 9.99×10^-5^) without any indication of inflation (genomic control inflation factor λ_GC_ = 1.01) due to genetic structure (**Fig. 2a** and **Supplementary Table 7**). In addition, we also performed a multi-SNP genome-wide association analysis using Bayesian variable-selection regression by piMASS^21^, which provided consistent and strong evidence for the association within the *BALF2* region (posterior probability = 0.96) (**Fig. 2b**). We evaluated the statistical significance of association using permutation (see **Methods**), and only the associations within the *BALF2* region reached genome-wide significance (suggestive genome-wide significance, *P* < 4.07×10^-4^). Consisting with the extensive linkage disequilibrium (LD) within the EBV genome (**Supplementary Fig. 5**), conditioning on the genetic effects of the SNPs within the *BALF2* region greatly reduced the extensive associations across the entire EBV genome (**Supplementary Fig. 6**).

**Figure 2.**
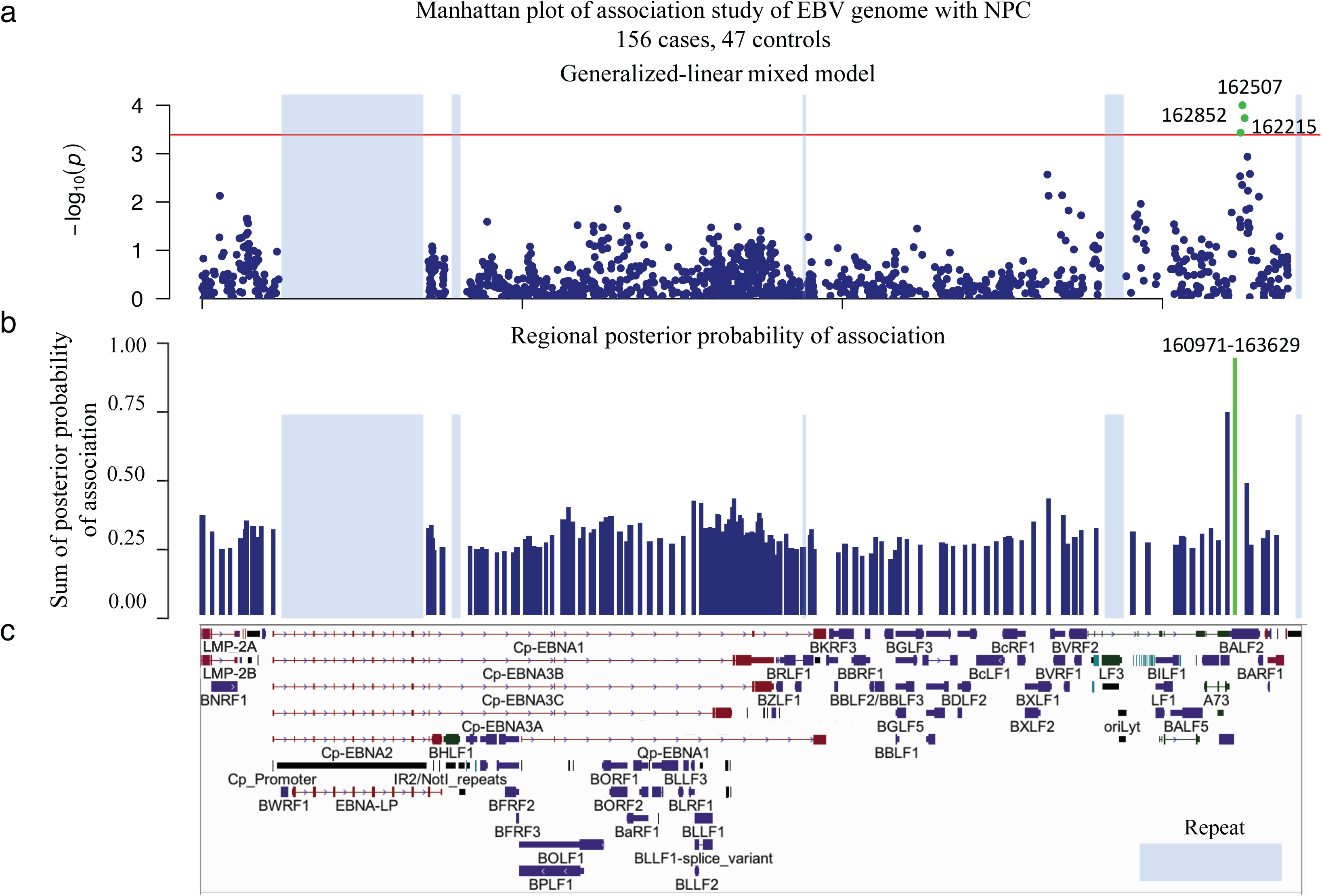
Genome-wide association analysis of EBV variants in 156 NPC cases and 47 controls. (**a**) Manhattan plot of genome-wide *P* values from the association analysis using generalized-linear mixed model. The −log_10_-transformed *P* values (y axis) of 1545 variants in 156 NPC cases and 47 controls are presented according to their positions in the EBV genome. The minimum *P* value (SNP162507) is 9.99×10^-5^. Suggestive genome-wide significance *P* value threshold of 4.07×10^-4^ (red line) was shown. (**b**) The regional plot of posterior probability of association. EBV genome was partitioned into overlapping 20-variant bins with 10-variant overlap between adjacent bins. The sum of the posterior probability for variants was assigned to each region. One region with strong evidence (> 0.90) for association with NPC risk is shown in green. (**c**) Schematic of EBV genes. Repetitive regions in EBV genomes are masked by light blue.

We performed a Bayesian fine-mapping analysis to prioritize potentially causal SNPs in the *BALF2* gene region using PAINTOR and found that only the three non-synonymous coding variants (SNPs 162215, 162476, and 163364) showed significant evidence for association (**Supplementary Fig. 7 and Supplementary Table 8**). We genotyped the three non-synonymous coding SNPs in an independent sample of 483 NPC cases and 605 age- and sex-matched healthy population controls (**Validation phase**) (**Supplementary Table 9**). To eliminate any potential impact of population stratification, all the cases and controls were recruited from the single NPC-endemic region, Zhaoqing County in the Guangdong Province of China. All three SNPs were significantly associated with NPC risk in the independent sample (*P* < 0.017, 0.05/3), consistent with the discovery phase results (**Table 1**). The meta-analysis of the combined discovery and validation phases confirmed the associations with the three SNPs of *BALF2* with genome-wide significance according to both permutation analysis and Bonferroni correction for multiple testing (SNP 162215_C, OR = 7.62, *P* = 2.98×10^-19^; 162476_C, OR = 8.79, *P* = 5.90×10^-26^ and 163364_T OR = 6.52, *P* = 7.18×10^-36^) (**Table 1**). All the three SNPs showed significant LD (**Supplementary Fig. 8**), but conditional analysis demonstrated that the associations with SNPs 162215 and 162476 were correlated, whereas SNP 163364 showed an independent association that also reached genome-wide significance (**Table 1**).

**Table 1.** The association of three non-synonymous SNPs in *BALF2* gene and their odds ratios for NPC risk.

| Position | Genotypes | High-risk genotype | Discovery |  |  | Validation |  |  | Combined |  |  | Odds ratio (95% CI) | *P* value conditional on SNPs |  | Annotation |
| --- | --- | --- | --- | --- | --- | --- | --- | --- | --- | --- | --- | --- | --- | --- | --- |
|  |  |  | 156 cases | 47 controls | *P* value | 483 cases | 605 controls | *P* value | 639 cases | 652 controls | *P* value |  | 163364 | 162476 |  |
| 162215 | C/A | C | 96.15% | 65.96% | 3.69E-04 | 95.03% | 74.71% | 1.84E-16 | 95.31% | 74.08% | 2.98E-19 | 7.62 (5.02-11.57) | 1.30E-04 | 2.39E-01 | *BALF2*, V700L |
| 162476 | T/C | C | 93.59% | 61.70% | 4.44E-03 | 94.00% | 65.12% | 1.21E-24 | 93.90% | 64.88% | 5.90E-26 | 8.79 (5.90-13.09) | 1.75E-06 |  | *BALF2*, I613V |
| 163364 | C/T | T | 88.46% | 48.94% | 5.83E-03 | 83.85% | 45.45% | 1.83E-35 | 84.98% | 45.71% | 7.18E-36 | 6.52 (4.90-8.68) |  | 3.23E-13 | *BALF2*, V317M |

Frequencies of high-risk genotypes in discovery, validation and combined analyses are indicated. The association of three EBV SNPs with NPC risk was tested by meta-analysis of the combined discovery and validation phases. Conditional regression analyses were performed in combined data sets and *P* values of SNP association are listed. Odds ratios conferred by high-risk genotypes and the 95% confidence intervals (CI) were estimated by meta-analysis of combined discovery and validation phases.

We further explored the association between the haplotypes (strains) composed of SNPs 162215, 162476 and 163364 and NPC risk. Taking the haplotype composed of the 3 low-risk variants (A-T-C) as a reference, we did not observe association for the haplotype carrying the high-risk variant for SNP162215 (C-T-C; odd ratio (OR) = 1.10, *P* = 0.83), although the number of haplotypes for testing was limited (**Table 2** and **Supplementary Table 10**). Both the haplotypes carrying the high-risk variants of either all the three SNPs or only SNPs 162215 and 162476 showed strong risk effect (haplotype C-C-T: OR = 12.10, *P* = 1.17×10^-25^; haplotype C-C-C: OR = 3.30, *P* = 2.20×10^-5^) (**Table 2** and **Supplementary Table 10**), but the haplotype C-C-T showed significantly stronger effect than the haplotype C-C-C (*P* = 1.85×10^-12^), clearly indicating the additional risk effect of SNP163364. The haplotype analysis further confirmed that NPC risk is primarily associated with SNPs 162476 and 163364, and SNP162215 needs to be further evaluated. We also performed pair-wise interaction analysis showing no evidence for interaction between SNPs 162476 and 163364 (*P* = 0.67). Lastly, we performed a multiple regression analysis that yielded independent risk effects (OR) of 3.15 for SNP 162476_C and 3.68 for SNP 163364_T (**Supplementary Table 11**), which were consistent with the risk effect of the haplotype carrying the two high-risk variants (haplotype C-C-T: OR = 12.10) (**Table 2**).

**Table 2.** EBV haplotypes composed of SNPs 162215, 162476 and 163364 and their odds ratios for NPC risk in 639 cases and 652 controls.

| EBV subtype<br>(162215-162476-163364) | 639 cases |  | 652 controls |  | Odds ratio<br>(95% CI) * | *P* value |
| --- | --- | --- | --- | --- | --- | --- |
|  | no. | % | no. | % |  |  |
| L-L-L (A-T-C) | 25 | 3.91% | 171 | 26.23% |  |  |
| H-H-H (C-C-T) | 539 | 84.35% | 293 | 44.94% | 12.10 (7.59 - 19.30) | 1.17E-25 |
| H-H-L (C-C-C) | 57 | 8.92% | 118 | 18.10% | 3.30 (1.90 - 5.72) | 2.20E-05 |
| H-L-L (C-T-C) | 13 | 2.03% | 65 | 9.97% | 1.10 (0.49 - 2.47) | 8.27E-01 |
| other subtypes | 5 | 0.78% | 5 | 0.77% | 4.63 (1.04 - 20.54) | 4.40E-02 |

\* Odds ratios of individual EBV subtypes and 95% confidence intervals (CI) were estimated with logistic model by categorizing each subtype as a single variable and adjusted for age, sex and the status of single- or multiple-infection in combined discovery and validation data sets. Subjects with EBV subtype A-T-C, a common low-risk subtype were used as the reference category. H represents high-risk genotypes; L represents low-risk genotypes.

### The population distribution and phylogenetic analysis of EBV risk haplotypes for NPC

In China, the frequency of the two high-risk haplotypes (C-C-T and C-C-C) was very high in the NPC-endemic region (93.27% in NPC cases and 63.04% in controls), but much lower in non-endemic areas (55% in NPC cases, 14.29% in controls) (**Supplementary Table 12**). The high-risk haplotypes were also observed in other EBV-associated cancers with a frequency of about 40% in lymphoma patients from the NPC-endemic region and 8.3% in the lymphoma and gastric carcinoma patients from the non-endemic regions. These frequencies are comparable to those observed in healthy controls and much lower than those observed in NPC patients (**Supplementary Table 12**). However, the number of samples from the non-endemic region and other EBV-association cancers were much smaller than the NPC samples from the endemic region. Interesting, the two risk haplotypes were absent or extremely rare in non-Asian populations (African and western countries) (**Supplementary Table 12**), suggesting the Asian origin of the EBV high-risk variants.

To further explore the origin of the EBV risk variants, we investigated the evolutionary relationship among the EBV strains from the current study and the previously published ones. By examining the frequency and distribution of heterozygous SNPs, we identified 229 EBV single strains from the 269 genome isolates (see **Methods**, **Supplementary Fig. 9 and Supplementary Table 13**). Using these 229 strains and C666-1 EBV genome sequence from the current study as well as 97 publicly available genomes, we performed phylogenetic inference analysis and found that the evolutionary relationship among all sequences was highly unbalanced, with a deep split between Type 1 and Type 2 EBV isolates (**Fig. 1b**). All Type 2 EBV isolates were geographically restricted to Africa, as previously observed^14,15,22^. The Type 1 EBV clade showed a continuous branching starting from Africa, Europe, and Asia, matching the overall distribution along the first PC in the PCA analysis (**Fig. 1b-d**). As shown in previous study^17,23^, 97% of all 269 EBV isolates were found to be China 1 subtype, and 2% were China 2 (defined by LMP1 classification) (**Supplementary Fig. 10**). Within the Asian group, isolates from NPC-non-endemic areas clustered towards the basal position of the lineage, similar to the pattern observed along the second PC in the PCA map (**Fig. 1b-d**). The most striking pattern in the phylogenetic relationship was a rapid radiation of NPC-dominant strains in the endemic population from southern China. EBV genomes from NPC patients appeared to have expanded recently from a common ancestor, and more than half (22 of 37) of healthy controls from this region were also infected with NPC-dominant strains (**Fig. 1b, c**).

We also mapped the three SNPs of *BALF2* (SNP 162215, 162476, and 163364) onto the phylogenetic tree of the EBV genomes. We observed that all the strains carrying the risk variants of SNP162476, and 163364 were within the Asian subclade, whereas the carriers of SNP 162215 had a much broader distribution (**Fig. 1b, e**). Within the Asian subclades, the carriers of SNP162476 and 163364 were enriched in the strains from NPC patients (NPC-dominant strains). These results provided strong evidence for the Asian origin of SNP162476 and 163364 and were consisted with their risk effect on NPC. The results also suggested that SNP 162215 was less likely to be a risk variant for NPC, and its association effect was due to its LD with SNP162476 (LD R-squared = 0.67).

### Functional analysis of EBV *BALF2* nonsynonymous variants

Since all three risk variants encode amino acid alterations within the *BALF2* gene that is responsible for opening the viral DNA for lytic replication. In order to explored the functional role of the three NPC-associated EBV variants, we investigated whether the three viral SNPs influenced viral lytic DNA replication. First, we performed *in vitro* functional analysis in EBV-positive NPC cell line TW03. After the stimulation of lytic cycle activation, we measured viral DNA abundance within cells that were transfected with the reference haplotype of B95-8 (C-T-C), the low-risk haplotype (A-T-C) and the high-risk haplotype (C-C-T) of *BALF2* and the empty vector. We found that the viral DNA level was significantly lower in cells carrying the high-risk haplotype compared to cells carrying the other two haplotypes (*P* < 0.05). No difference was observed between the low-risk and reference haplotypes, and between the high-risk haplotype and the empty vector (**Fig. 3a**). These results indicated that both the reference and low-risk haplotypes of *BALF2* had similar ability, whereas the high-risk haplotype had weaker ability in supporting EBV lytic DNA replication. Furthermore, we performed an *in vivo* analysis of the viral DNA abundance in the saliva samples from the 533 NPC cases and 651 healthy controls of the validation sample. We observed a large variation of viral DNA load across the samples, and found that viral DNA abundance in saliva was significantly lower in NPC patients than in healthy controls (*P* = 6.6×10^-15^) (**Fig. 3b**). However, we did not observe the association of the high-risk haplotype (C-C-T) of *BALF2* with saliva viral DNA abundance (**Supplementary Table 14**).

**Figure 3.**
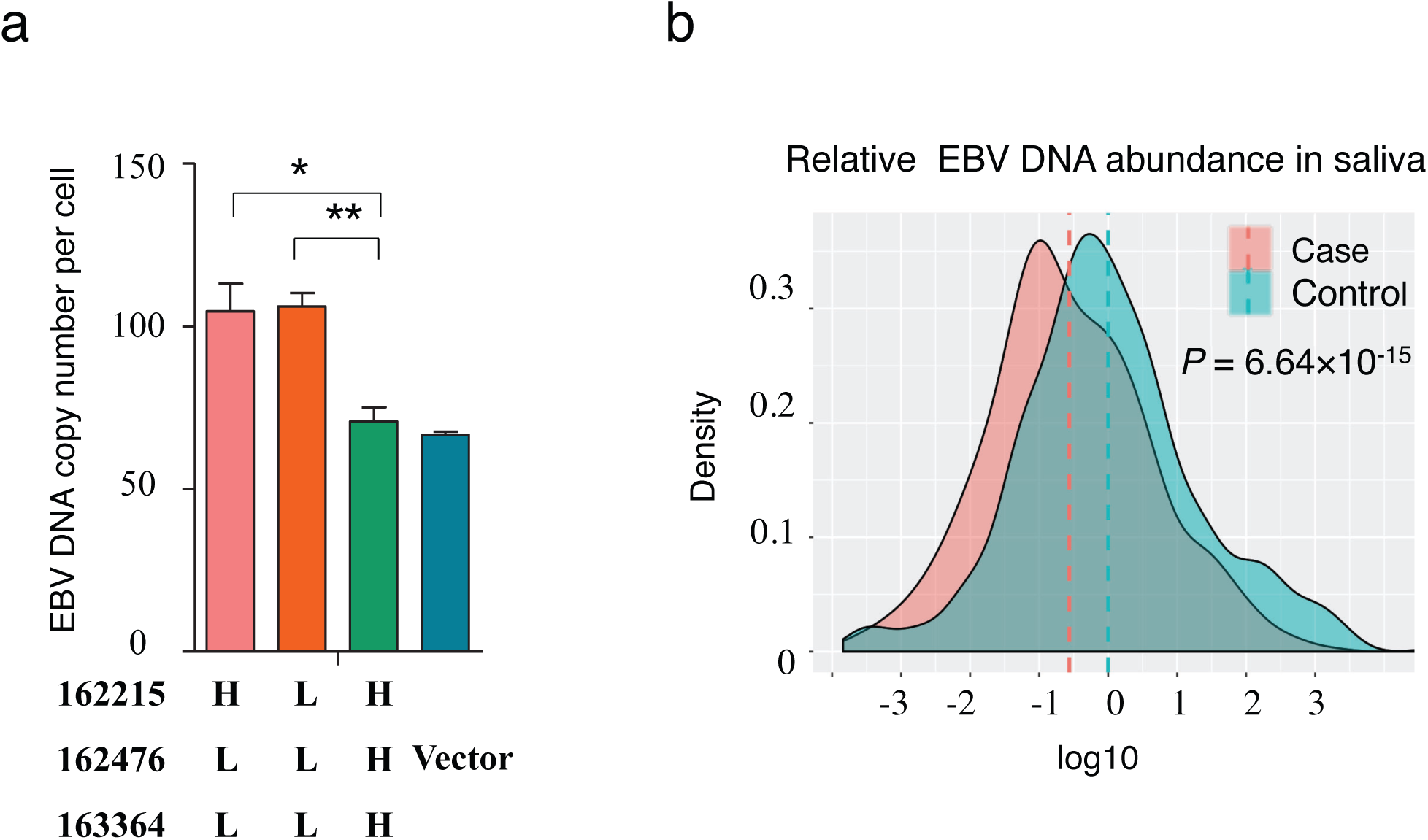
Functional analysis of high-risk genotypes of SNPs 162215, 162476, and 163364 in terms of regulation of EBV lytic DNA replication. (**a**) Effect of the *BALF2* haplotypes composed of three SNPs on EBV lytic DNA replication. EBV DNA inside cells was measured by quantitative PCR in the NPC TW03 cell line 48 hours after induction of viral lytic cycle activation. The cells were transfected with the following haplotypes: reference haplotype H-L-L (C-T-C), low-risk haplotype L-L-L (A-T-C) and high-risk haplotype H-H-H (C-C-T). The cells transfected with empty vector were used as controls. L: low-risk genotype; H: high-risk genotype. Significant differences in EBV DNA amount were calculated by Student’s *t* test: * *P* = 0.024, ** *P* = 0.004. Means and s.d. of three independent experiments are shown. (**b**) Distribution of EBV DNA abundance in saliva samples from 533 NPC cases and 651 controls relative to the average abundance in controls. Of the total 536 saliva samples from cases, three had missing EBV DNA amount values. Distribution frequency density of relative EBV DNA amount is shown. *P* value was determined using linear regression adjusted for age and sex.

## Discussion

Because of the ubiquity of EBV infection, the determinants of the distinctive geographical distribution of NPC have long puzzled the scientific community. Using large-scale sequencing and functional analyses, we discovered for the first time two EBV coding SNPs 162476 and 163364 showing a strong risk effect for NPC. The more than 6-fold increase in NPC risk conferred by these two high-risk EBV variants is far greater than the effects of any other known risk factors for this disease, including host genetic variants (**Table 1**). In particular, with a population frequency of 45% and an OR of 12.10, the EBV haplotype C-T of the two SNPs is the dominant NPC risk factor, contributing 71% (95% confidence interval: 67-74%) of the overall risk of NPC in the endemic population of southern China. The second risk haplotype C-C also contributed about 10%, such that the two high-risk EBV haplotypes combined accounted for 83% (95% confidence interval: 79-87%) of NPC risk in this population (**Supplementary Table 15**). In non-endemic regions of China, the frequency of these high-risk haplotypes is much lower (about 10%), but they still make a significant contribution (~50%) to NPC risk. The frequency of the two high-risk EBV subtypes was not associated with the risk of developing other EBV-related cancers in our study, suggesting that their oncogenic effects might be specific to NPC. However, this observation would benefit from further work since our study was only powered to explore NPC.

When we mapped these two causal variants onto the phylogenetic tree of EBV genomes, we observed a distinct subclade of EBV subtypes carrying the two high-risk variants within Asia. The carriers could only be found inside of Asia, which strongly demonstrates the Asian origin of these two risk variants. Most interestingly, the phylogenetic analysis suggests a rapid clonal expansion of these unique high-risk EBV subtypes. This is consistent with the current distribution of these subtypes in China with very high frequency in the NPC-endemic region (93.27% in NPC cases and 63.04% in controls), but much lower in the non-endemic areas (55% in NPC cases, 9.68% in other samples) (**Supplementary Table 12**). It remains to be investigated what kind of positive selection has driven their emergence. Taken together, the strong risk effect, the confined geographic distribution, and the rapid clonal expansion and consequently extremely enriched frequency of these two high-risk variants in the NPC-endemic region strongly suggest that these two EBV risk factors are the driving factors for the unique endemics of NPC in southern China.

Our genetic discovery has provided novel biological insight in the development of NPC by highlighting the important role of suppressed viral lytic DNA replication in EBV-mediated NPC tumorigenesis. Consistent with the role of *BALF2* as the core component in the viral DNA replication complex, we showed that the high-risk haplotype of *BALF2* suppressed the lytic DNA replication using the *in vitro* cell line analysis. Consistently, EBV DNA abundance in saliva was found to be significantly lower in the NPC cases than in controls, suggesting that EBV in buccal epithelium is less lytic in NPC patients. EBV latency promotes the expression of oncogenes and is therefore indispensable for EBV-mediated carcinogenesis^24,25^, and the expansion of EBV latently-infected nasopharyngeal cells has proven to be an early event in NPC tumorigenesis^26,27^. Our results demonstrated that the impairment of *BALF2* function due to EBV genetic variation potentially promotes viral latency and fosters NPC development by suppressing viral lytic replication. The discovery of these high-risk EBV variants also has major implications for public health efforts to reduce the burden of NPC, particularly in the endemic region of southern China. Testing for these high-risk EBV variants can enable the identification of high-risk individuals for targeted implementation of routine clinical monitoring for early detection of NPC. Primary prevention by developing vaccines against high-NPC-risk EBV strains is expected to lead to great attenuation of the Cantonese Cancer in China.

## Methods

### Study participants and samples

Participants of the current study were enrolled through two recruitments. The first one was a hospital-based study enrolling patients diagnosed with EBV-related cancers, including NPC, Burkitt lymphoma, Hodgkin lymphoma, NK/T cell lymphoma and gastric carcinomas as well as healthy controls from the Sun Yat-sen university Cancer Center in Guangdong Province, the First Affiliated Hospital of Guangxi Medical College in Guangxi Province, and the Affiliated Hospital of the Qingdao University in Shandong Province of China. The geographical origin of the participants covers NPC-endemic southern China (Guangdong and Guangxi Provinces where NPC has highest incidence of 20-40/100,000 individuals per year) and non-endemic regions in China where NPC is rare. After quantitative measurement of EBV DNA, 170 tumor tissue and/or saliva, plasma samples were selected from the first recruitment for EBV whole-genome sequencing (WGS). The second recruitment was a population-based NPC case-control study enrolling NPC cases and population control subjects from Zhaoqing County, Guangdong Province of China (NPC-endemic region). The population controls were matched to the cases by their age and sex. Saliva samples were collected from all the subjects. After quantitative measurement of EBV DNA of the second study, 99 saliva samples (53 NPC cases and 46 controls) were selected for EBV WGS. Written informed consent was obtained from each participant before undertaking any study-related procedures, and all studies was approved by institutional ethics committee of Sun Yat-sen University Cancer Center.

Detailed sample information including the geographic origin of the 269 isolates used for WGS was given and summarized in **Supplementary Table 1**. For discovery phase of EBV whole-genome association study (GWAS) with NPC, we selected 156 cases and 47 controls exclusively from the NPC-endemic region out of the 269 EBV WGS isolates. For the validation phase, 990 NPC cases and 1105 healthy controls from the endemic population-based case-control study were used by genotyping GWAS candidate SNPs (For details, see **Supplementary Note** and **Supplementary Fig. 11**).

### Sample processing

Saliva samples were collected into vials containing lysis buffer (50 mM Tris, pH 8.0, 50 mM EDTA, 50 mM sucrose, 100 mM NaCl, 1% SDS). Tumor specimens were obtained from biopsy samples collected during surgical treatment and confirmed by histopathological examination. Both saliva and tumor specimens were stored at -80 °C. DNA was extracted from the saliva using the Chemagic STAR (Hamilton Robotics, Sweden) and from the tumor biopsy, plasma and NPC cell line C666-1 using the DNeasy blood and tissue kit (Qiagen).

### EBV genome quantification, whole genome sequencing and variant calling

Using real time PCR targeting a DNA fragment at the *BALF5* gene (5’ and 3’ primers, GGTCACAATCTCCACGCTGA and CAACGAGGCTGACCTGATCC), we quantified the amount EBV DNA in each sample. The mean Ct values from three independent replicates was used to select patient samples for viral whole genome sequencing (Ct value < 30, detailed information can be found in the **Supplementary Note**).

EBV genomes were captured using the MyGenostics GenCap Target Enrichment Protocol (GenCap Enrichment, MyGenostics, USA). After capture enrichment, DNA libraries were prepared and sequenced using the Illumina HiSeq 2000 platform according to standard protocols (Illumina Inc., San Diego, CA, USA). After raw sequence processing and quality control, paired-end reads were aligned to the EBV B95-8 reference genome (NC_007605.1) using the Burrows-Wheeler Aligner (BWA, version 0.7.5a)^28^ ^29^. The average sequencing depth was 1,282 (range, 32 to 6,629). High coverage (average, 98.02%; range, 94.44% to 99.91%;) was achieved. (**Supplementary Fig. 1**).

Following GATK’s best practice (version 3.2-2), an initial set of 8,469 variants were first called after base and variant recalibration and filter ^30^. In order to avoid inaccurate calling, we further filtered out variants that has (i) low coverage support (depth < 10×), (ii) in repetitive elements, (iii) within 5 bp of an indel, and 7,962 variants were retained for subsequent EBV phylogenetic, principal component and association analyses. The functional annotation of the EBV variants was performed using the SNPEff package according to the reference genome (NC_007605.1, NCBI annotation, Nov 2013)^31^. A complete description of the sequencing and variant calling is presented in the **Supplementary Note**. Sequencing and variant statistics did not find outlier of EBV isolates sequenced in current study (**Supplementary Fig. 1**).

To evaluate the accuracy of our sequencing and variant calling, subsets of EBV variants were validated using either the Sanger sequencing or MassAarray iPLEX assay (Agena Bioscience). Two independent technologies can provide orthogonal evaluations of the sequencing accuracy. 299 PCR fragments were amplified from 53 randomly selected EBV isolates and re-sequenced using the Sanger sequencing. Comparing the SNPs called by WGS and the Sanger sequencing revealed a concordance rate of 97.55% (**Supplementary Table 4**). Similarly, a high concordance rate of 99.988% between the WGS and MassArray iPLEX assay was found when genotyping 37 variants in 239 selected samples (**Supplementary Table 5**). In addition, when comparing the re-sequenced C666-1 EBV genome against the publicly available sequence^18^, a high concordance rate of 97.93% was found (**Supplementary Table 3**).

In order to understand viral genomes from multiple sample types from the same patient, two EBV fragments (position 80,089 to 80,875 and position 81,092 to 81,829) containing 89 SNPs were resequenced using the Sanger method from paired saliva and tumor samples from the same set of patients. Across 25 NPC patients with paired tumor and saliva samples, pairwise difference (defined as the genotype discordance rate at the 89 SNPs) between the tumor samples of the 25 patients (inter-host difference) as well as between the paired tumor and saliva samples of the same patient (intra-host difference) were calculated and compared (**Supplementary Fig. 3**). The median inter-patient difference was 13.5% (1st to 3rd quartile: 3.7-16.9%), and the median intra-host difference was only 1.1% (1st to 3rd quartile: 0-3.4%). High concordance rate between saliva and tumor tissue suggests that paired saliva and tumor sample from the same patient are highly similar.

### Genotyping analysis of EBV variants by MassArray iPLEX

In order to genotype the EBV variants in the Zhaoqing 990 cases and 1105 controls, genotyping was conducted using customized primers and following the recommended protocol by the Agena Bioscience MassArray iPLEX platform. A fixed position within the human albumin gene was used as a positive control. Since the genotyping success rate strongly correlates with the EBV DNA abundance (**Supplementary Fig. 12**), about half of the validation samples (483 of cases and 605 of controls) could be successfully genotyped for all the three GWAS candidate markers (i.e. SNP 162215, 162476 and 163364). The slightly lower success rate in cases is consistent with the fact that the EBV DNA abundance was lower in the saliva from patients than the healthy controls. For detailed information, see **Supplementary Note**.

### Determining single *versus* multiple EBV infections

Previous studies have found that EBV genome usually underwent clonal expansion in NPC tumors or other malignancies^26,27,32^. In the scenario of clonal expansion, EBV genome is stable and intra-host mutation rate is often low, and heterozygous variants as a result of quasi-species evolution within a host are not frequent^12,18,33^. On the contrary, EBV isolates from specimens with multiple infections will have a higher number of heterozygous variants. We plotted the percentage of heterozygous variants across all the 270 samples from the WGS analysis and observed that heterozygosity (defined as percentage of heterozygous variants) across all the samples showed two different distributions with low and high numbers of heterozygous variants. By fitting two curves to the lower and higher quantiles of the empirical distribution, we defined the reflection point (i.e. the intersection of the two distributions) as the cutoff value (**Supplementary Fig. 9**). Samples with the proportion of heterozygous variants lower than the cutoff value were identified as single-infection samples, whereas samples above this threshold were identified as multi-infection samples. For the validation cohort, samples with the homozygous calls at all the three EBV SNPs were regarded as patients infected by single EBV haplotypes. For samples with multiple EBV haplotype infection, haplotypes of the three SNPs were inferred by Beagle 4.1^34^. For details, see **Supplementary Note**.

### Phylogenetic and principal component analyses of EBV genome sequences

The phylogenetic and principal component analyses were performed using 230 EBV single-infection isolates and 97 publicly accessible EBV genomes. For the phylogenetic analysis, we first created the fasta sequence for each resequenced isolate using the variant data extracted from the variant calling. The 230 whole genomes were subsequently combined with the 97 public genomes and multiple sequence alignment was carried out using the Multiple Alignment using Fast Fourier Transform (MAFFT)^35^. After masking the regions of repetitive sequences and poor coverage in resequencing maximum likelihood inference of the phylogenetic relationship was conducted using the Randomized Axelerated Maximum Likelihood (RAxML) assuming a General Time Reversible (GTR) model^36^. The inferred phylogeny was subsequently rooted using the Evolutionary Placement Algorithm (EPA) algorithm^37^ from RAxML using a Macacine herpesvirus 4 genome sequence (NC_006146) as the outgroup.

In the PCA analysis, genomic variation from the 97 public genomes was generated by global pairwise sequence alignment of published genome sequences against the B95-8 reference genome (NC_007605) using the EMBOSS Stretcher^38^. The variant set is then combined with the polymorphism data extracted from WGS. A combined set of 12,182 SNPs from the 270 newly-sequenced isolates and 97 published ones were then used for the PCA analyses. During the PCA analysis, SNPs were first filtered based on allele frequency (minor genotype frequency > 0.05) and linkage disequilibrium (LD pruning with pairwise correlation r2 value > 0.6 within a 1000-bp sliding window). 495 SNPs were included in the PCA analysis using the R package “SNPRelate”^39^.

### Principal component analysis of the cases and controls

To assess the human population structure of the 156 cases and 47 healthy controls used for the EBV GWAS discovery phase, the human DNAs of these samples were genotyped using the OmniZhongHua-8 Chip (Illumina). After sample filtering based on a series of criteria including (i) the calling rate (above 95%), (ii) SNP filtering by minor allele frequency (above 5%), (iii) Hardy-Weinberg equilibrium (P > 1×10^-6^), (iv) LD-based SNP pruning (r^2^ < 0.1 and not within the five high-LD regions^5^), PCA analysis was performed using the PLINK (Version 1.9) based on the discovery samples alone or by combining them with reference samples from the 1000 Genome project ^19^.

### Association analysis

Genetic association analysis of EBV variants was performed by testing either single or multiple variants. Single variant association analysis was performed using generalized-linear mixed model with EBV genetic relatedness matrix as random effect^20^. Sex and age were included as fixed effect, as well as four human PCs to correct for any potential impact of host population structures on the association results. Both single- and multiple-infection samples were included in the association analysis with the status of single- or multiple-infection being included as a covariate to correct for any potential confounding effect due to multiple infection. The genome-wide discovery analysis was performed by testing 1,545 EBV variants (with missing rate < 10%, minor genotype frequency > 0.05 and heterozygosity < 0.1) in 156 cases and 47 healthy controls. The validation analysis was performed by testing three EBV non-synonymous coding SNPs (162215, 162476 and 163364) of *BALF2* in additional 483 cases and 605 population healthy controls that were matched to the cases in term of age and sex from the case-control study in Zhaoqing County, Guangdong China. The logistic regression model was used for validation phase with the adjustment of age, sex and status of single- or multiple-infection of EBV strains. The meta-analysis of the discovery and validation phases was performed with zscore pooling method. Considering the extensive LD across the EBV genome, to obtained a suggestive genome-wide significance of association, we used permutation of a logistic model with adjustment of age, sex, status of single- or multiple-infection and host and EBV population structures. The genome-wide significance (4.07×10^-4^) was determined by 5% quantile of the empirical distribution of minimum *P*-values from 10, 000 permutations, as the data-driven threshold to control family wise error rate under multiple correlated testing.

The genome-wide multi-variant-based association analysis was performed by testing 1477 bi-allelic EBV variants in Bayesian variable selection regression implemented in piMASS^21^. Age, sex, four human PCs and two EBV PCs were included as covariates. The analysis was performed by partitioning EBV genome into the regions of 20-SNP sliding window with 10 overlapping SNPs. The sum of the posterior probabilities of the SNPs (being associated) within a window was calculated as the “region statistic” indicating the strength of the evidence for genetic association in that region.

To further prioritize potentially casual SNPs in the top hit *BALF2* gene region for validation, we applied further fine-mapping analysis using Bayesian multiple variable selection by PAINTOR3.1^40^. Functional annotation of SNPs was used as a prior to compute the probability of being causal SNPs for each variant in the region. We assumed a single causal variant in *BALF2* genes and calculated 90% credible set which contains the minimum set of variants that jointly have at least 90% probability of including the causal variant.

### Estimation of population attributable fraction of risk

The proportion of NPC risk explained by the effect of the two high-risk haplotypes of SNPs 162476 and 163364 (C-T and C-C) was estimated in the validation sample. The attributable fraction of risk and 95% confidence interval were estimated in a logistic regression model with adjustment for age and sex by R package ‘AF’^41^. As NPC is not a common disease (prevalence < 40/100,000), the risk ratio can be approximated by OR. Thus, the population attributable fraction can be approximated by *AF* ≈ 1 – *E*_*z*_{*OR*^*-X*^(*Z*) | *Y* = 1}.

### Functional analysis of NPC-associated BLAF2 SNPs in cell line

The DNA fragment of the *BALF2* reference haplotype C-T-C was obtained from B95-8 EBV-BAC plasmid p2089^42^ using PCR. The DNA fragments carrying one of the three non-synonymous SNPs 162215, 162476 and 163364 were constructed by site-directed mutagenesis through PCR. The reference and three mutated *BALF2* haplotype PCR fragments was subsequently cloned into the vector (pCDH-CMV-MCS-EF1-Puro), and sequences were verified by Sanger sequencing. The four *BALF2* constructs and the empty vector were transfected into 293T cells using polyethylenimine (PEI) for lentivirus production. Lentivirus infection with EBV-positive TW03 cells and selection of stable cell line were performed as previously described^43^. Overexpression of the *BALF2* construct was confirmed by Western blot.

EBV lytic cycle was induced in TW03 cells using TPA (phorbol-12-myristate-13-acetate) (20 ng/ml) and SB (sodium butyrate) (2.5mM) for 12 hours. After 12 to 48 hours’ culture, total viral DNA within the cells and in the supernatant of culture were extracted using Qiagen DNeasy Blood & Tissue Kits and Qiagen QIAamp DNA Blood Mini Kit, respectively. Three biological replicates were conducted. EBV DNA copy number in the supernatant or cells was measured in triplicate relative to a standard curve by quantitative PCR, and the measurements were normalized by the total number of cells in culture.

## Supporting information

## Acknowledgements

We want to thank all the participants for their generous support for the current study. We would also like to thank Dr. Cui Jie from the Wuhan Institute of Virology, Chinese Academy of Sciences for helpful discussions on viral evolution and phylogenetic analysis, Dr. Zhen Lin from Tulane University for kindly sharing EBV genome annotation files, Mr. Wen-Sheng Liu and Dr. Xiaoyu Zuo from Sun Yat-sen University Cancer Center for providing code supports. This work was supported by the National Natural Science Foundation of China (key project, 81430059), the National Key R&D Program of China (No. 2016YF0902000), the National Cancer Institute at the US National Institutes of Health (R01CA115873-01), 1000-talent award (to W.Z.) of China, and the Agency of Science, Technology and Research (A*STAR), Singapore.

## Author contributions

Y.-X.Z., J.L. and W.Z. were the principal investigators who conceived the study. Y.-X.Z and M.X. obtained financial support. Y.-X.Z., J.L., W.Z. and M.X. designed and oversaw the study. J.L. and X.L. provided supervision over viral genome-wide association study. M.X., Y.Y. and L.Z. performed sample preparation, quality control, sequencing and genetic statistical analysis. H.C. and W.Z. performed the phylogenetic analysis. M.X, S.Z. and Y.Y. performed genotyping by MassArray iPlex assay. M.X., S.Z. and G.H. performed functional experiments. H.-O.A., W.Y. and Y.-X.Z. supervised the design and implementation of the population-based case-control study in Zhaoqing. W.Y., E.T.C., S.-M.C., W.J., S.-H.X., Z.L. participated in the case-control design, study recruitment, and sample storage and preparation. Z.Z. was responsible for the collection of NPC tissue samples from Guangxi. B.L. was responsible for the collection of NPC and EBV-GC tissue samples from North China. X.G., M.-Y.C. and R.-J.P. were responsible for the collection of NPC and EBV-lymphoma tissue samples from Guangdong. The manuscript was drafted by M.X., J.L., W.Z. and Y.-X.Z. and revised by V.P. and E.T.C.. All the authors critically reviewed the article and approved the final manuscript.

