## Supplementary material for "Genome sequencing analysis identifies high-risk Epstein-Barr virus subtypes for nasopharyngeal carcinoma"

**Contents**

**Page 3 – Supplementary Figure 1** Sequencing and variant statistics of each EBV genome isolates indicate no outliers among the 269 EBV isolates.
**Page 4 – Supplementary Figure 2** Regions encoding latent proteins have highest diversity across EBV genomes.
**Page 5 – Supplementary Figure 3** The variant discordance rate between paired tumor and saliva samples Versus between tumors from different patient (inter-host) difference.
**Page 6 – Supplementary Figure 4** Human principal component analysis of the samples used for EBV genome-wide association analysis.
**Page 7 – Supplementary Figure 5** EBV genome-wide linkage disequilibrium heatmap.
**Page 8 – Supplementary Figure 6** NPC and EBV genome association study conditional on SNPs 162215 and 132048.
**Page 9 – Supplementary Figure 7** Posterior probability of association for variants in BALF2 gene region was estimated by PAINTOR.
**Page 10 – Supplementary Figure 8** Linkage disequilibrium structure of BALF2 gene region.
**Page 11 – Supplementary Figure 9** Distribution of genome-wide heterozygous variants in 270 EBV genome isolates.
**Page 12 – Supplementary Figure 10** Classification of 230 newly-sequenced EBV isolates and 97 published EBV isolates based on LMP-1 C-terminal signatures.
**Page 13 – Supplementary Figure 11** Flowchart of participant recruitment from the hospital-based and population-based studies.
**Page 14 – Supplementary Figure 12** Average Ct (cycle of threshold) value of quantitative PCR of EBV DNA in the samples with 0-3 SNPs successfully genotyped in validation phase of association study.
**Page 15 – Supplementary Table 1** List and summary of 270 EBV isolates newly sequenced and 97 publicly accessed genomes included in the analysis.
**Page 35 – Supplementary Table 2** Variant information of EBV genome isolates sequenced in current study.

**Page 42 – Supplementary Table 3** Concordance rate between SNPs from C666-1 EBV genome sequenced in current study and in published study.
**Page 43 – Supplementary Table 4** Concordance rate between variants discovered by targeted EBV whole-genome sequencing (EBV-WGS)and Sanger sequencing.
**Page 44 – Supplementary Table 5** Concordance rate between variants discovered by targeted EBV whole-genome sequencing (EBV-WGS) andMassArray iPlex assay.
**Page 45 – Supplementary Table 6** Variant comparison between EBV isolates from paired saliva and NPC tumor samples from the same NPC patient .
**Page 46 – Supplementary Table 7** Top three associated SNPs in GWAS discovery phase reached genome-wide significance (P < 4.07×10-4).
**Page 47 – Supplementary Table 8** Fine-mapping for casual SNPs associated with NPC risk in BALF2 gene region.
**Page 49 – Supplementary Table 9** Basic characteristics of 483 cases and 605 control individuals used for validation phase by age and sex.
**Page 50 – Supplementary Table 10** EBV subtypes determined by SNPs 162215, 162476 and 163364 and their odds ratios for NPC risk in 536 and 651 population-based cases and controls.
**Page 51 – Supplementary Table 11** Estimation of odds ratios of SNP 162476 and 163364 for NPC risk.
**Page 52 – Supplementary Table 12** Frequency of high-risk EBV haplotypes in different regions.
**Page 53 – Supplementary Table 13** The percentage of heterozygous variants in 270 EBV genome isolates.
**Page 59 – Supplementary Table 14** The association of EBV haplotypes with EBV DNA abundance in saliva of 533 cases and 651 controls.
**Page 60 – Supplementary Table 15** Estimation of the proportion of NPC population risk attributable to high-risk EBV haplotypes in population-based NPC 536 cases and 651 controls.
**Page 61 – Supplementary Note**

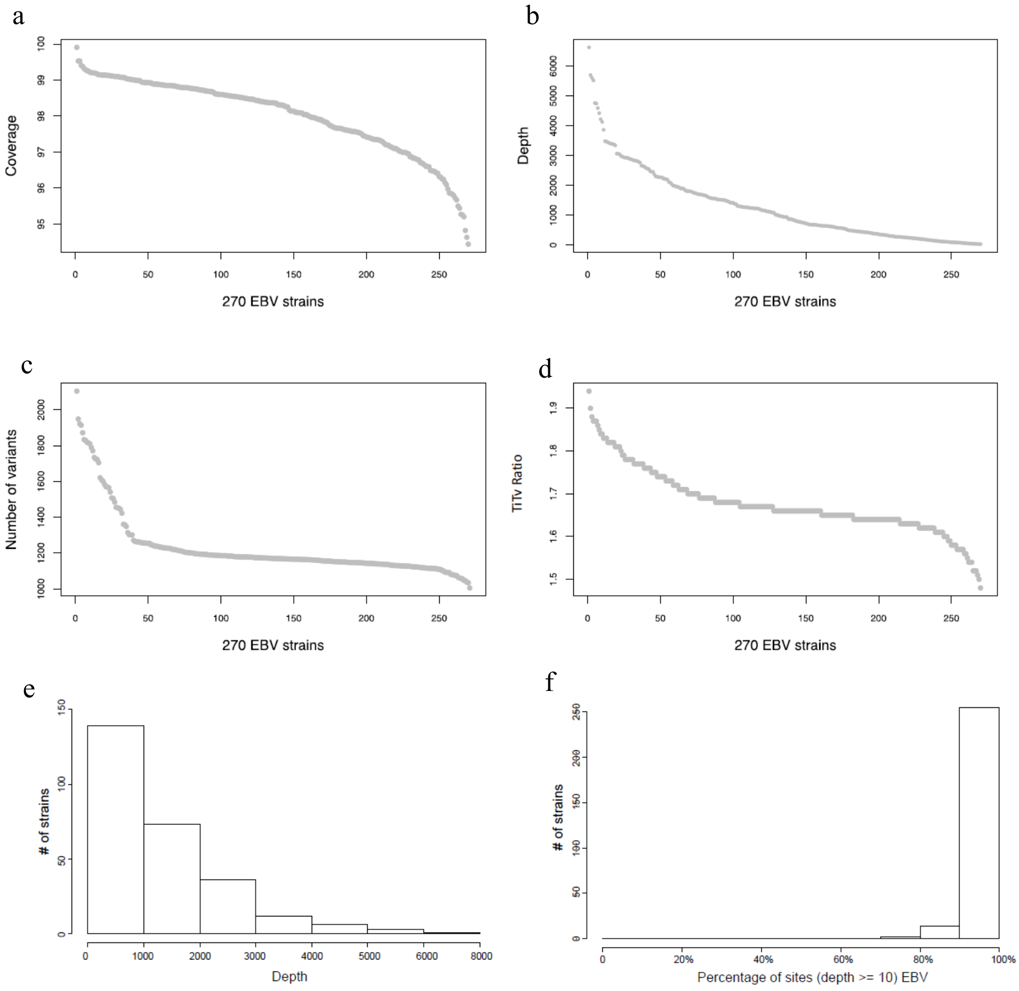


**Supplementary Figure 1 Sequencing and variant statistics of each EBV genome isolates indicate no outliers among the 269 EBV isolates.** **(a)** Sequencing coverage across EBV genome, ranging from 94% to 99%. **(b)** Average sequencing depth. **(c)** Number of variants. **(d)** Ratio of transition to transversion. **(e)** Frequency histogram of average sequencing depth per isolate. **(f)** Frequency histogram of percentage of reference genome that was covered by 10 or more reads.

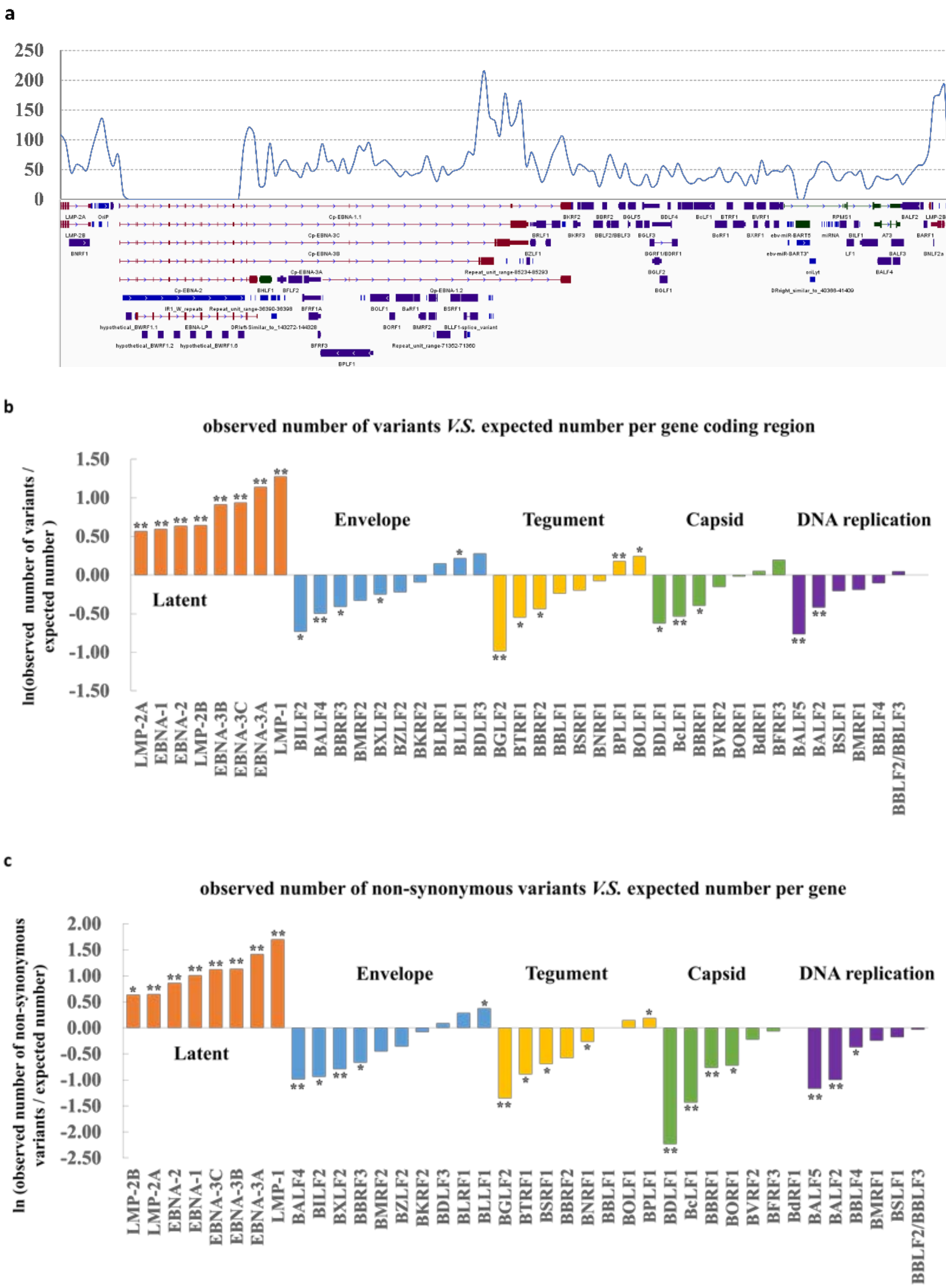


**Supplementary Figure 2 Regions encoding latent proteins have highest diversity across EBV genomes. (a)** Variant frequency across EBV genomes derived from 270 samples. The line graph is plotted across the genome showing the total number of variants in a sliding 1000-nt window. **(b)** Comparison of the observed and expected numbers of variants in gene coding regions. Expected numbers of variants were calculated by multiplying the length of gene coding regions by the number of variants per kilo base. *P* values were calculated by Fisher's exact tests. *, $P < 0.05$; **, $P < 0.001$.**(c)** Comparison of the observed and expected numbers of non-synonymous variants in gene coding regions. Expected numbers of non-synonymous variants were calculated by multiplying the length of gene coding regions by the number of non-synonymous variant per kilobase. *P* values were calculated by Fisher's exact tests. *, $P < 0.05$; **, $P < 0.001$.

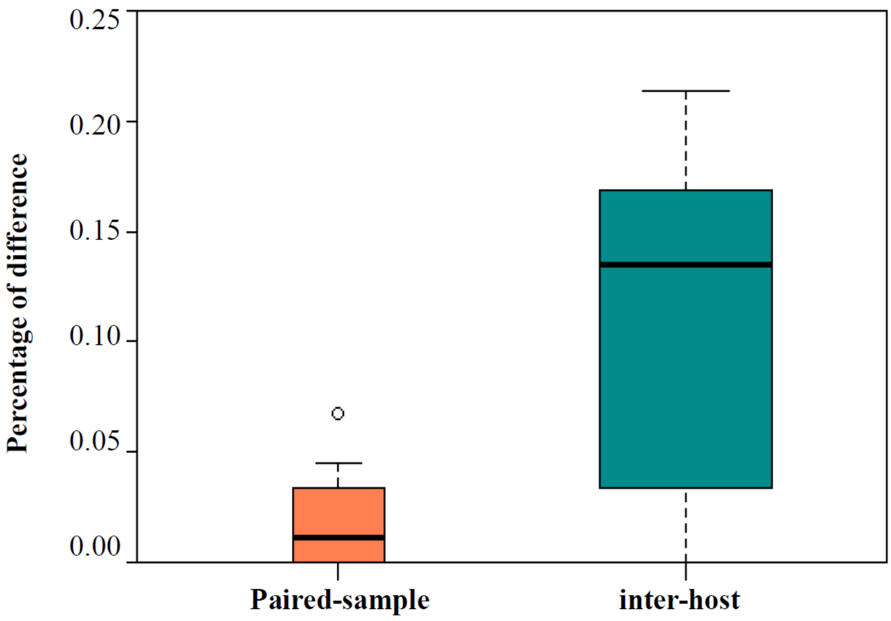


**Supplementary Figure 3 The variant discordance rate between paired tumor and saliva samples *Versus* between tumors from different patient (inter-host) difference.** EBV DNA fragments were sequenced from 25 pairs of NPC tumor and saliva samples. Median, 1st and 3rd quartiles were shown.

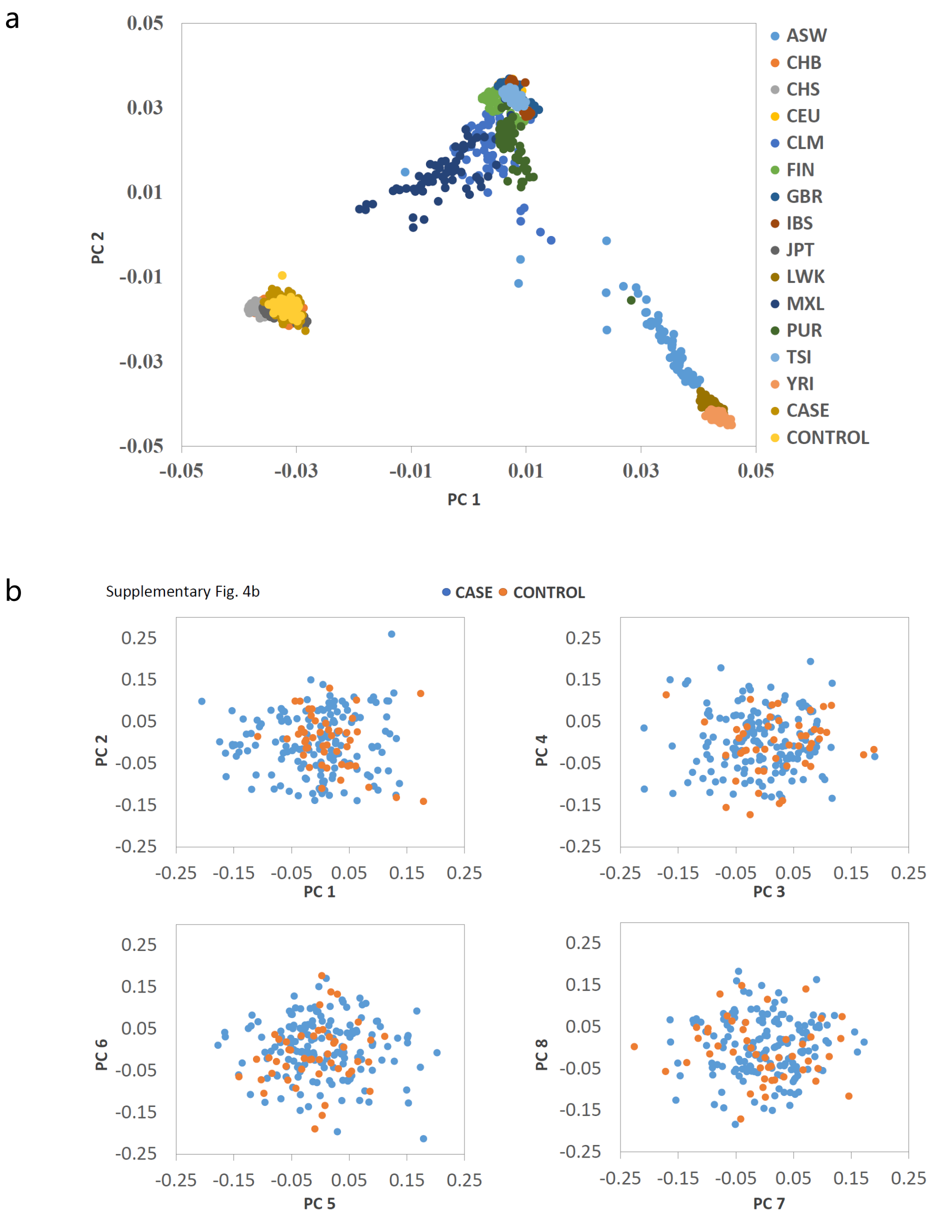


**Supplementary Figure 4 Human principal component analysis of the samples used for EBV genome-wide association analysis. (a)** The PC scores for each sample were plotted against the first two PCs (PC 1 and PC 2), together with 1000 genome project samples. No outlier was observed between our cases and controls using appropriate criteria in the paper of Price 2006, which defines individuals whose ancestry is at least 6 standard deviations from the mean of one of the top ten PC values as outlier . Population codes and NPC cases and controls used for EBV GWAS were listed at the right panel. ASW, Americans of African ancestry in SW USA; CEU, Utah Residents with Northern and Western European Ancestry; CHB, Chinese Han in Beijing, China; CHS, Southern Han Chinese; CLM, Colombians from Medellin, Colombia; FIN, Finnish in Finland; GBR, British in England and Scotland; IBS, Iberian Population in Spain; JPT, Japanese in Tokyo, Japan; LWK, Luhya in Webuye, Kenya; MXL, Mexican Ancestry from Los Angeles USA; PUR, Puerto Ricans from Puerto Rico; TSI, Toscani in Italia; YRI, Yoruba in Ibadan, Nigeria. (**b**) The PC scores for each NPC case and control were plotted against the first eight PCs (PC 1 to PC 8).

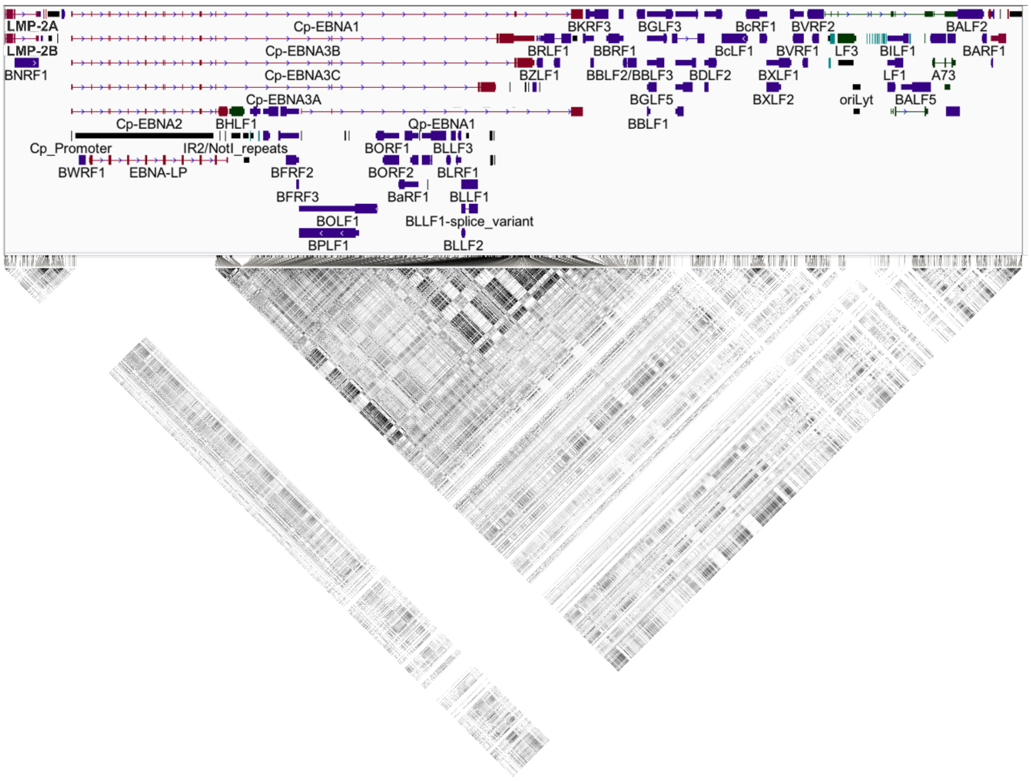


**Supplementary Figure 5 EBV genome-wide linkage disequilibrium heatmap.** Pair-wise R-squared values between 1545 variants with minor genotype frequency > 0.05 in 156 NPC cases and 47 controls were plotted in lower plot. Higher linkage disequilibrium is presented with darker blocks. The upper panel shows EBV genome annotation.

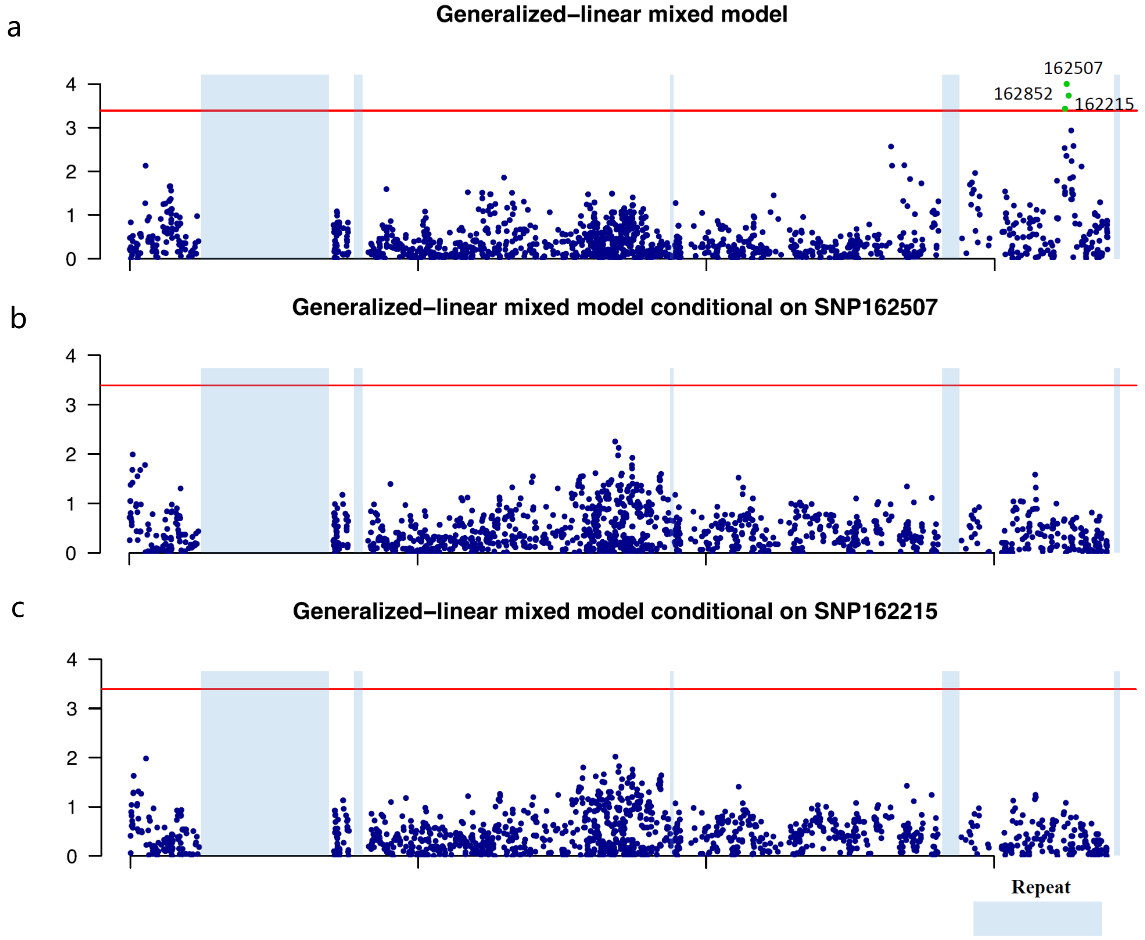


**Supplementary Figure 6 NPC and EBV genome association study conditional on SNPs 162215 and 132048. (a)** Manhattan plot of the genome-wide *P* values of association study. Association was assessed by generalized-linear mixed model with age, sex, status of single or multiple EBV infection and four human PCs as fixed effect and genetic relatedness matrix as random effect. Genome-wide significant *P* value threshold was $4.07 \times 10^{-4}$. Associations reached genome-wide significance were highlighted by green. **(b-c)** Logistic regressions were conditional on SNPs 162507 and 162215 as indicated.

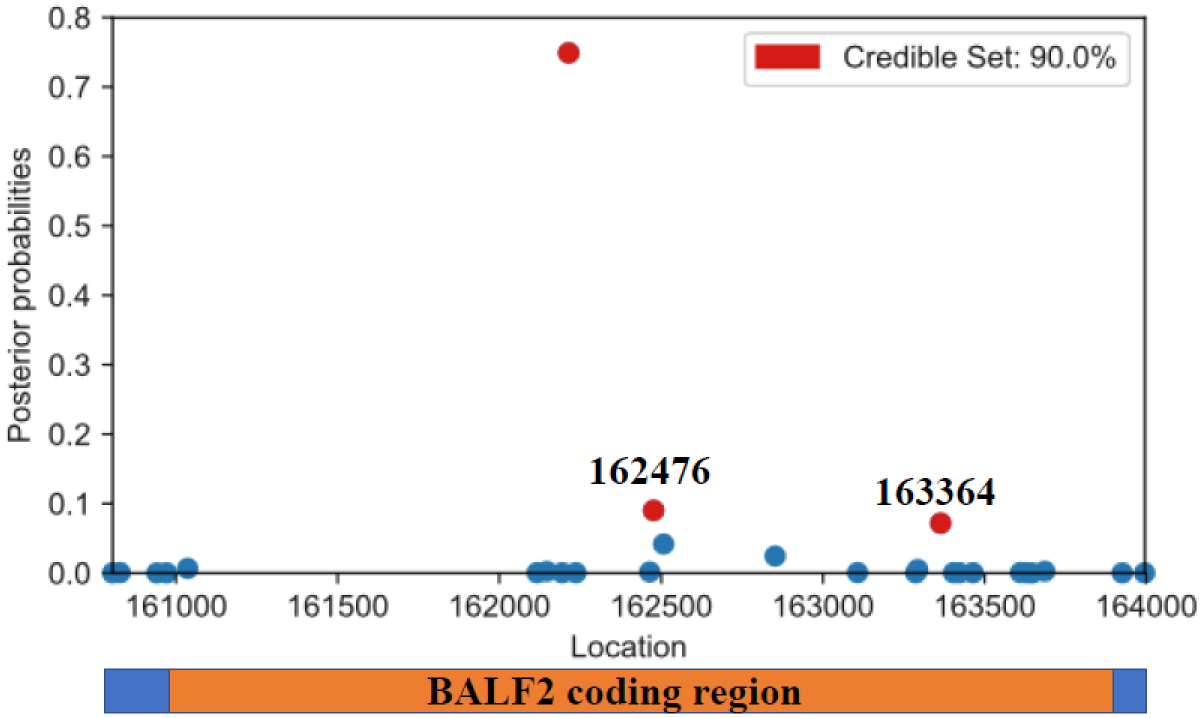


**Supplementary Figure 7 Posterior probability of association for variants in BALF2 gene region was estimated by PAINTOR.**

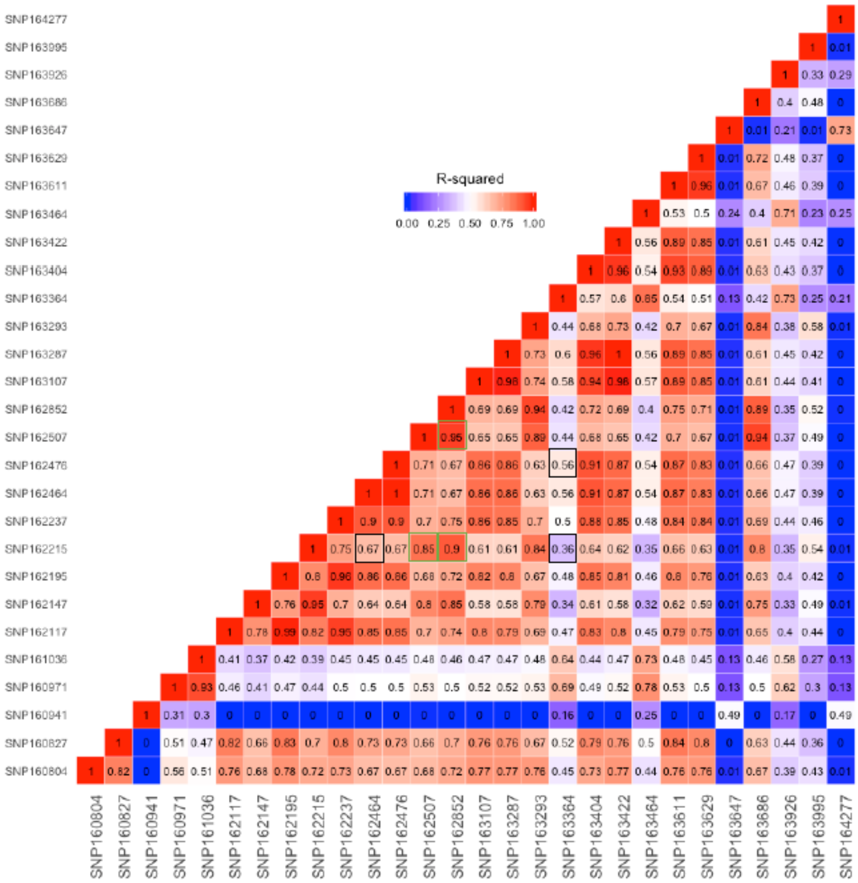


**Supplementary Figure 8** Linkage disequilibrium structure of BALF2 gene region. Pair-wise r-squared values of 28 SNPs in BALF2 gene region are shown. The R-squared values of SNPs 162215, 162476 and 163364 are highlighted with black squares. The R-squared values of SNPs 162215 and 162507, 162850 are highlighted with green squares.

a

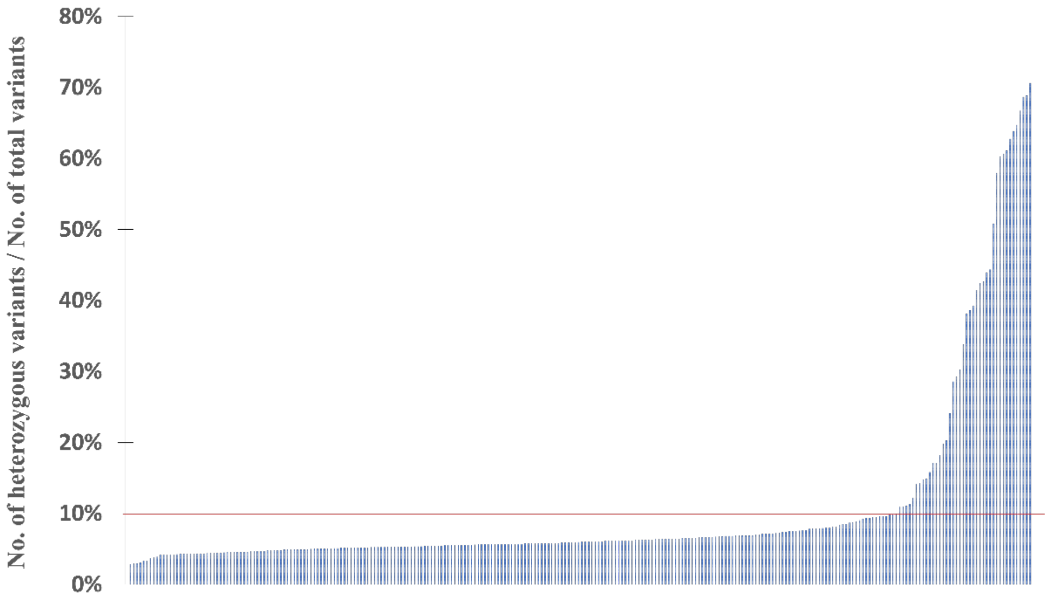


b

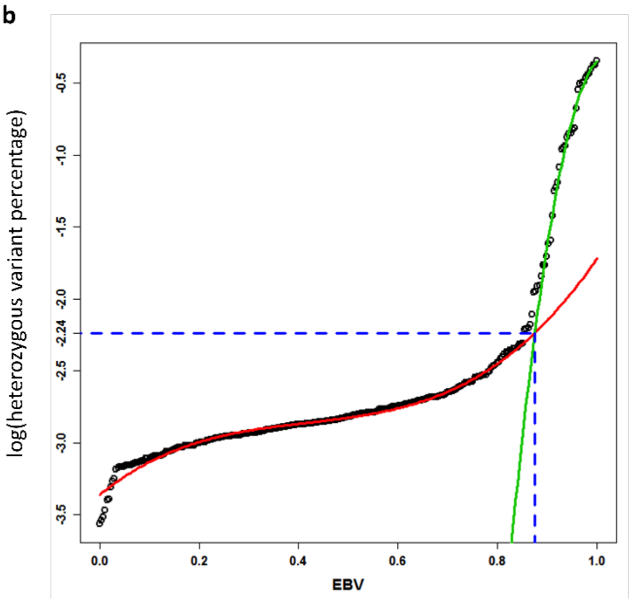


**Supplementary Figure 9 Distribution of genome-wide heterozygous variants in 270 EBV genome isolates.** (**a**) Heterozygous variant percentage in 270 EBV genome isolates. Y axis is the percentage of heterozygous variants out of total variants per sample. Red line represents the threshold of 10.7%. (**b**) Heterozygosity of 10.7% was determined as the cut-off for single infection. Two curves of heterozygosity of single infections were fitted (EBV isolates with heterozygous variant proportion < 8%) and multiple infections (isolates with heterozygous variant proportion > 15%) with cubic model. Since the distribution of heterozygous variant proportion is markedly skewed, we applied log-transformation and used the transformed heterozygous variant proportion as outcome in the analysis. The two fitted curves intersected at heterozygosity of 10.7%, which was used as the cut-off value of single infection determination.

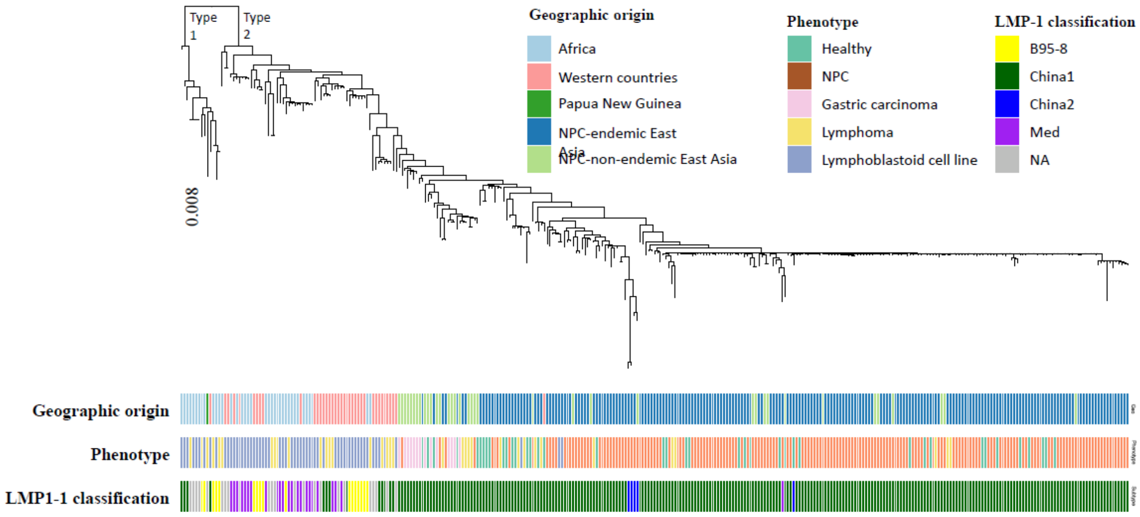


**Supplementary Figure 10 Classification of 230 newly-sequenced EBV isolates and 97 published EBV isolates based on LMP-1 C-terminal signatures.** Phylogeny of 327 EBV strains. Macacine herpesvirus 4 genome sequence (NC_006146) was used as the outgroup to root the tree. LMP-1 classifications, geographical origins and phenotypes from which EBV strains were sequenced are shown with colors as indicated.

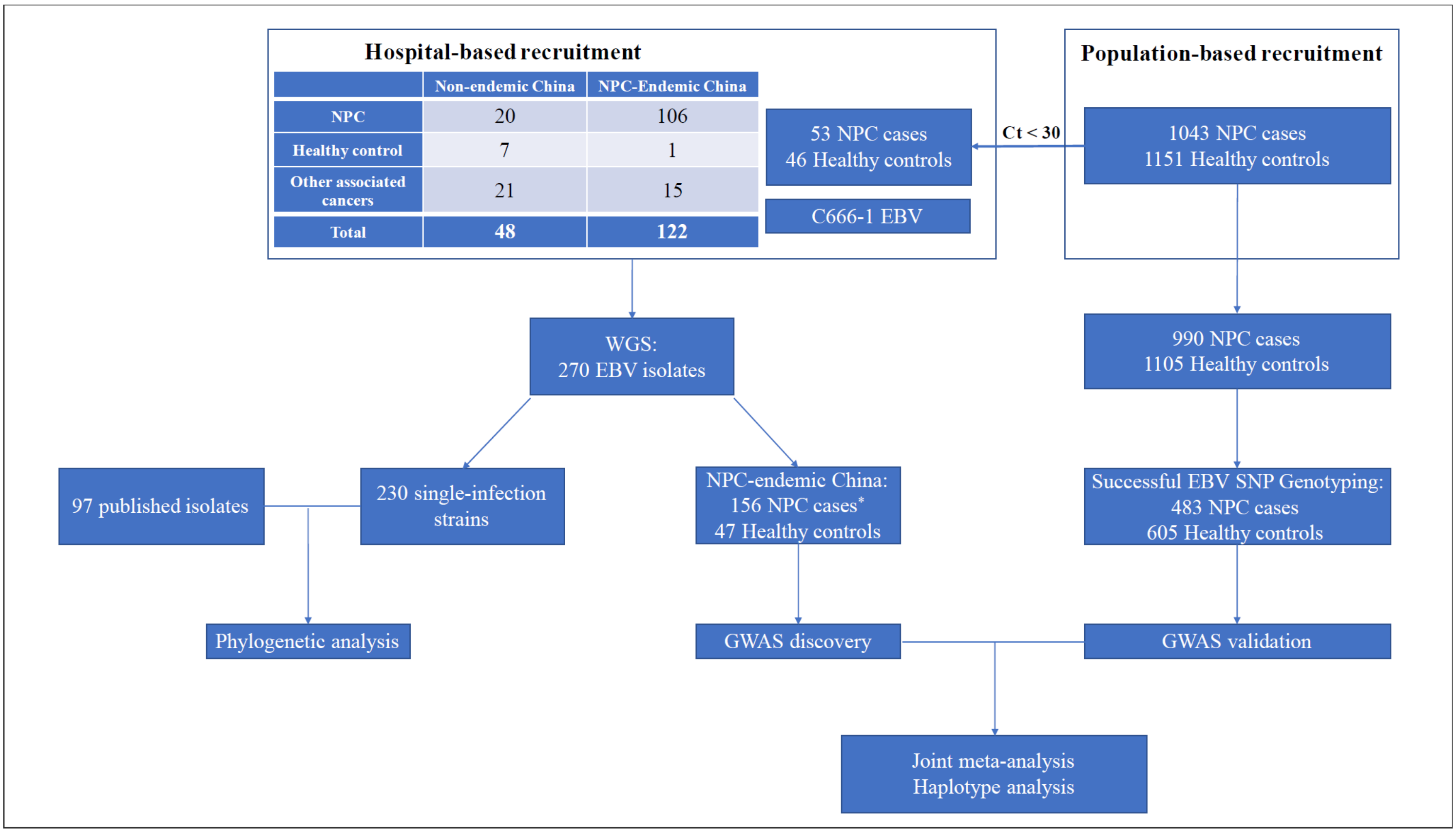


**Supplementary Figure 11** Flowchart of participant recruitment from the hospital-based and population-based studies. *Three EBV isolates from NPC tumor biopsies from NPC-endemic China were excluded from GWAS discovery phase because each of the three isolates had the other paired isolate obtained from the same patient which had been included in GWAS. All the EBV isolates included in GWAS discovery and validation phase were from independent samples.

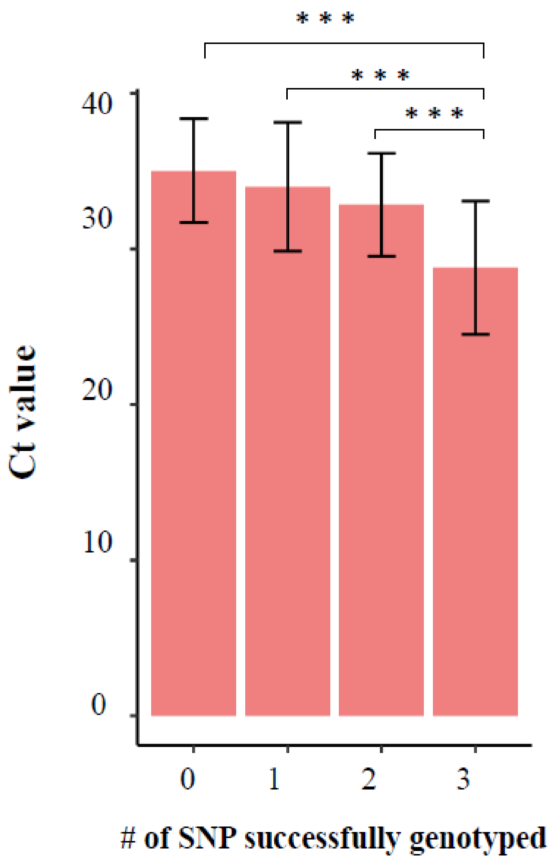


**Supplementary Figure 12** Average Ct (cycle of threshold) value of quantitative PCR in the samples with 0-3 SNPs successfully genotyped. Ct value increases by 1, the relative DNA amount decreases by 0.5 fold. ***, $P$ < 10e-7. $P$ values were determined by ANOVA test.

**Supplementary Table 1 List and summary of 270 EBV isolates sequenced in current study and 97 publicly accessed genomes included in the analysis.** (**a**) Discription of 270 EBV isolates sequenced in current study and 97 publicly accessed genomes. Geographic origins, phenotypes, samples types, and recruitments, sex and age of participants if EBV isolates were sequenced in current study are indicated. We list the NCBI GenBank accession numbers of published isolates and from which references we selected the isolates. We specify whether EBV genomes sequenced in current study were single isolates. The EBV Type1/Type2 and LMP-1 C terminal classification of single EBV isolates are indicated. The analysis for which the particular isolates were used is also indicated. **(b)** Summary of geographic origins and phenotypes of 367 EBV isolates. **(c)** Summary of  age and sex of participants from whom EBV were sequenced in current study. **(d)** Summary of geographic origins, phenotypes and sample types of 270 EBV isolates sequenced in current study.

**Supplementary Table 1 List and summary of 270 EBV isolates sequenced in current study and 97 publicly accessed genomes included in the analysis**

**(a) Discription of 270 EBV isolates sequenced in current study and 97 publicly accessed genomes**

| Sample ID | Sequenced by | Geographic origin | Detailed geographic origin | Phenotype | Sample type | Recruitment | Sex | Age | Strain | EBV Type | LMP1 classification | Analysis | Notes |
|---|---|---|---|---|---|---|---|---|---|---|---|---|---|
| BLT001 | Current study | East Asia | NPC-endemic China | Burkitt's lymphoma | tumor biopsy | SYSUCC | male | 36 | single | Type1 | China1 | phylogeny | |
| BLT002 | Current study | East Asia | NPC-endemic China | Burkitt's lymphoma | tumor biopsy | SYSUCC | male | 34 | single | Type1 | China1 | phylogeny | |
| C666 | Current study | East Asia | NPC-endemic China | NPC | cell line | | male | 50 | single | Type1 | China1 | phylogeny | |
| GCT001 | Current study | East Asia | NPC-non-endemic China | Gastric carcinoma | tumor biopsy | AHQU | male | 43 | single | Type1 | China1 | phylogeny | |
| GCT002 | Current study | East Asia | NPC-non-endemic China | Gastric carcinoma | tumor biopsy | AHQU | female | 51 | single | Type1 | China1 | phylogeny | |
| GCT003 | Current study | East Asia | NPC-non-endemic China | Gastric carcinoma | tumor biopsy | AHQU | male | 26 | single | Type1 | China1 | phylogeny | |
| GCT004 | Current study | East Asia | NPC-non-endemic China | Gastric carcinoma | tumor biopsy | AHQU | male | 44 | single | Type1 | China1 | phylogeny | |
| GCT005 | Current study | East Asia | NPC-non-endemic China | Gastric carcinoma | tumor biopsy | AHQU | male | 63 | single | Type1 | China1 | phylogeny | |
| GCT006 | Current study | East Asia | NPC-non-endemic China | Gastric carcinoma | tumor biopsy | AHQU | male | 57 | single | Type1 | China1 | phylogeny | |
| GCT007 | Current study | East Asia | NPC-non-endemic China | Gastric carcinoma | tumor biopsy | AHQU | male | 44 | single | Type1 | China1 | phylogeny | |
| GCT008 | Current study | East Asia | NPC-non-endemic China | Gastric carcinoma | tumor biopsy | AHQU | male | 42 | | | | | |
| GCT009 | Current study | East Asia | NPC-non-endemic China | Gastric carcinoma | tumor biopsy | AHQU | male | 57 | single | Type1 | China1 | phylogeny | |
| GCT010 | Current study | East Asia | NPC-non-endemic China | Gastric carcinoma | tumor biopsy | AHQU | female | 42 | single | Type1 | China1 | phylogeny | |
| GCT011 | Current study | East Asia | NPC-non-endemic China | Gastric carcinoma | tumor biopsy | AHQU | male | 60 | single | Type1 | China1 | phylogeny | |
| GCT012 | Current study | East Asia | NPC-non-endemic China | Gastric carcinoma | tumor biopsy | AHQU | male | 55 | single | Type1 | China1 | phylogeny | |
| GCT013 | Current study | East Asia | NPC-non-endemic China | Gastric carcinoma | tumor biopsy | AHQU | male | 67 | single | Type1 | China1 | phylogeny | |
| GCT014 | Current study | East Asia | NPC-non-endemic China | Gastric carcinoma | tumor biopsy | AHQU | male | 69 | single | Type1 | China1 | phylogeny | |
| GCT015 | Current study | East Asia | NPC-non-endemic China | Gastric carcinoma | tumor biopsy | AHQU | male | 43 | | | | | |
| GCT016 | Current study | East Asia | NPC-non-endemic China | Gastric carcinoma | tumor biopsy | AHQU | male | 44 | | | | | |
| HLT001 | Current study | East Asia | NPC-endemic China | Hodgkin's lymphoma | tumor biopsy | SYSUCC | male | 35 | single | Type1 | China1 | phylogeny | |
| HLT002 | Current study | East Asia | NPC-endemic China | Hodgkin's lymphoma | tumor biopsy | SYSUCC | male | 69 | single | Type1 | China1 | phylogeny | |
| HLT003 | Current study | East Asia | NPC-endemic China | Hodgkin's lymphoma | tumor biopsy | SYSUCC | male | 56 | | | | | |
| HLT004 | Current study | East Asia | NPC-non-endemic China | Hodgkin's lymphoma | tumor biopsy | SYSUCC | male | 60 | | | | | |
| HLT005 | Current study | East Asia | NPC-endemic China | Hodgkin's lymphoma | tumor biopsy | SYSUCC | male | 54 | single | Type1 | China1 | phylogeny | |

**Supplementary Table 1 List and summary of 270 EBV isolates sequenced in current study and 97 publicly accessed genomes included in the analysis**

**(a) Discription of 270 EBV isolates sequenced in current study and 97 publicly accessed genomes**

| Sample ID | Sequenced by | Geographic origin | Detailed geographic origin | Phenotype | Sample type | Recruitment | Sex | Age | Strain | EBV Type | LMP1 classification | Analysis | Notes |
|---|---|---|---|---|---|---|---|---|---|---|---|---|---|
| HLT006 | Current study | East Asia | NPC-endemic China | Hodgkin's lymphoma | tumor biopsy | SYSUCC | male | 52 | single | Type1 | China1 | phylogeny | |
| HLT007 | Current study | East Asia | NPC-endemic China | Hodgkin's lymphoma | tumor biopsy | SYSUCC | male | 55 | single | Type1 | China1 | phylogeny | |
| HLT009 | Current study | East Asia | NPC-non-endemic China | Hodgkin's lymphoma | tumor biopsy | SYSUCC | male | 60 | | | | | |
| HLT010 | Current study | East Asia | NPC-endemic China | Hodgkin's lymphoma | tumor biopsy | SYSUCC | male | 59 | single | Type1 | China1 | phylogeny | |
| HLT011 | Current study | East Asia | NPC-endemic China | Hodgkin's lymphoma | tumor biopsy | SYSUCC | male | 60 | single | Type1 | China1 | phylogeny | |
| HLT012 | Current study | East Asia | NPC-non-endemic China | Hodgkin's lymphoma | tumor biopsy | SYSUCC | male | 14 | | | | | |
| HS001 | Current study | East Asia | NPC-non-endemic China | Healthy | saliva | SYSUCC | female | 29 | single | Type1 | China1 | phylogeny | |
| HS003 | Current study | East Asia | NPC-endemic China | Healthy | saliva | SYSUCC | female | 60 | single | Type1 | China1 | discovery phase of association study, phylogeny | |
| HS005 | Current study | East Asia | NPC-non-endemic China | Healthy | saliva | SYSUCC | female | 23 | | | | | |
| HS006 | Current study | East Asia | NPC-non-endemic China | Healthy | saliva | SYSUCC | male | 26 | | | | | |
| HS007 | Current study | East Asia | NPC-endemic China | Healthy | saliva | NPC case-control study | male | 56 | single | Type1 | China1 | discovery phase of association study, phylogeny | |
| HS008 | Current study | East Asia | NPC-endemic China | Healthy | saliva | NPC case-control study | male | 40 | single | Type1 | China1 | discovery phase of association study, phylogeny | |
| HS009 | Current study | East Asia | NPC-endemic China | Healthy | saliva | NPC case-control study | male | 46 | single | Type1 | China1 | discovery phase of association study, phylogeny | |
| HS010 | Current study | East Asia | NPC-endemic China | Healthy | saliva | NPC case-control study | male | 68 | | | | discovery phase of association study | |
| HS011 | Current study | East Asia | NPC-endemic China | Healthy | saliva | NPC case-control study | male | 36 | single | Type1 | China1 | discovery phase of association study, phylogeny | |
| HS012 | Current study | East Asia | NPC-endemic China | Healthy | saliva | NPC case-control study | male | 56 | single | Type1 | China1 | discovery phase of association study, phylogeny | |
| HS013 | Current study | East Asia | NPC-endemic China | Healthy | saliva | NPC case-control study | male | 37 | single | Type1 | China1 | discovery phase of association study, phylogeny | |
| HS014 | Current study | East Asia | NPC-endemic China | Healthy | saliva | NPC case-control study | male | 72 | single | Type1 | China1 | discovery phase of association study, phylogeny | |
| HS015 | Current study | East Asia | NPC-endemic China | Healthy | saliva | NPC case-control study | male | 58 | single | Type1 | China1 | discovery phase of association study, phylogeny | |
| HS016 | Current study | East Asia | NPC-endemic China | Healthy | saliva | NPC case-control study | male | 70 | single | Type1 | China1 | discovery phase of association study, phylogeny | |
| HS017 | Current study | East Asia | NPC-endemic China | Healthy | saliva | NPC case-control study | male | 46 | | | | discovery phase of association study | |
| HS018 | Current study | East Asia | NPC-endemic China | Healthy | saliva | NPC case-control study | male | 43 | single | Type1 | China1 | discovery phase of association study, phylogeny | |
| HS019 | Current study | East Asia | NPC-endemic China | Healthy | saliva | NPC case-control study | male | 35 | single | Type1 | China1 | discovery phase of association study, phylogeny | |
| HS020 | Current study | East Asia | NPC-endemic China | Healthy | saliva | NPC case-control study | male | 57 | single | Type1 | China1 | discovery phase of association study, phylogeny | |

**Supplementary Table 1 List and summary of 270 EBV isolates sequenced in current study and 97 publicly accessed genomes included in the analysis**

**(a) Discription of 270 EBV isolates sequenced in current study and 97 publicly accessed genomes**

| Sample ID | Sequenced by | Geographic origin | Detailed geographic origin | Phenotype | Sample type | Recruitment | Sex | Age | Strain | EBV Type | LMP1 classification | Analysis | Notes |
|---|---|---|---|---|---|---|---|---|---|---|---|---|---|
| HS021 | Current study | East Asia | NPC-endemic China | Healthy | saliva | NPC case-control study | male | 56 | single | Type1 | China1 | discovery phase of association study, phylogeny | |
| HS022 | Current study | East Asia | NPC-endemic China | Healthy | saliva | NPC case-control study | male | 60 | | | | discovery phase of association study | |
| HS023 | Current study | East Asia | NPC-endemic China | Healthy | saliva | NPC case-control study | male | 47 | single | Type1 | China1 | discovery phase of association study, phylogeny | |
| HS024 | Current study | East Asia | NPC-endemic China | Healthy | saliva | NPC case-control study | male | 56 | single | Type1 | China1 | discovery phase of association study, phylogeny | |
| HS025 | Current study | East Asia | NPC-endemic China | Healthy | saliva | NPC case-control study | male | 60 | single | Type1 | China1 | discovery phase of association study, phylogeny | |
| HS026 | Current study | East Asia | NPC-endemic China | Healthy | saliva | NPC case-control study | male | 55 | | | | discovery phase of association study | |
| HS027 | Current study | East Asia | NPC-endemic China | Healthy | saliva | NPC case-control study | male | 45 | single | Type1 | China1 | discovery phase of association study, phylogeny | |
| HS028 | Current study | East Asia | NPC-endemic China | Healthy | saliva | NPC case-control study | female | 26 | | | | discovery phase of association study | |
| HS029 | Current study | East Asia | NPC-endemic China | Healthy | saliva | NPC case-control study | male | 46 | single | Type1 | China1 | discovery phase of association study, phylogeny | |
| HS030 | Current study | East Asia | NPC-endemic China | Healthy | saliva | NPC case-control study | male | 43 | | | | discovery phase of association study | |
| HS031 | Current study | East Asia | NPC-endemic China | Healthy | saliva | NPC case-control study | male | 32 | | | | discovery phase of association study | |
| HS032 | Current study | East Asia | NPC-endemic China | Healthy | saliva | NPC case-control study | male | 40 | single | Type1 | China1 | discovery phase of association study, phylogeny | |
| HS033 | Current study | East Asia | NPC-endemic China | Healthy | saliva | NPC case-control study | male | 57 | single | Type1 | China1 | discovery phase of association study, phylogeny | |
| HS034 | Current study | East Asia | NPC-endemic China | Healthy | saliva | NPC case-control study | male | 40 | single | Type1 | China1 | discovery phase of association study, phylogeny | |
| HS035 | Current study | East Asia | NPC-endemic China | Healthy | saliva | NPC case-control study | male | 57 | single | Type1 | China1 | discovery phase of association study, phylogeny | |
| HS036 | Current study | East Asia | NPC-endemic China | Healthy | saliva | NPC case-control study | male | 73 | single | Type1 | China1 | discovery phase of association study, phylogeny | |
| HS037 | Current study | East Asia | NPC-endemic China | Healthy | saliva | NPC case-control study | male | 49 | single | Type1 | China1 | discovery phase of association study, phylogeny | |
| HS038 | Current study | East Asia | NPC-endemic China | Healthy | saliva | NPC case-control study | male | 40 | single | Type1 | China1 | discovery phase of association study, phylogeny | |
| HS039 | Current study | East Asia | NPC-endemic China | Healthy | saliva | NPC case-control study | male | 57 | single | Type1 | China1 | discovery phase of association study, phylogeny | |
| HS040 | Current study | East Asia | NPC-endemic China | Healthy | saliva | NPC case-control study | male | 45 | | | | discovery phase of association study | |
| HS041 | Current study | East Asia | NPC-endemic China | Healthy | saliva | NPC case-control study | male | 43 | single | Type1 | China1 | discovery phase of association study, phylogeny | |
| HS045 | Current study | East Asia | NPC-endemic China | Healthy | saliva | NPC case-control study | male | 49 | single | Type1 | China1 | discovery phase of association study, phylogeny | |
| HS046 | Current study | East Asia | NPC-endemic China | Healthy | saliva | NPC case-control study | male | 50 | | | | discovery phase of association study | |
| HS048 | Current study | East Asia | NPC-endemic China | Healthy | saliva | NPC case-control study | male | 50 | single | Type1 | China1 | discovery phase of association study, phylogeny | |

**Supplementary Table 1 List and summary of 270 EBV isolates sequenced in current study and 97 publicly accessed genomes included in the analysis**

**(a) Discription of 270 EBV isolates sequenced in current study and 97 publicly accessed genomes**

| Sample ID | Sequenced by | Geographic origin | Detailed geographic origin | Phenotype | Sample type | Recruitment | Sex | Age | Strain | EBV Type | LMP1 classification | Analysis | Notes |
|---|---|---|---|---|---|---|---|---|---|---|---|---|---|
| HS050 | Current study | East Asia | NPC-endemic China | Healthy | saliva | NPC case-control study | female | 47 | single | Type1 | China1 | discovery phase of association study, phylogeny | |
| HS051 | Current study | East Asia | NPC-endemic China | Healthy | saliva | NPC case-control study | female | 49 | single | Type1 | China1 | discovery phase of association study, phylogeny | |
| HS052 | Current study | East Asia | NPC-endemic China | Healthy | saliva | NPC case-control study | female | 67 | single | Type1 | China1 | discovery phase of association study, phylogeny | |
| HS053 | Current study | East Asia | NPC-endemic China | Healthy | saliva | NPC case-control study | female | 63 | single | Type1 | China2 | discovery phase of association study, phylogeny | |
| HS054 | Current study | East Asia | NPC-endemic China | Healthy | saliva | NPC case-control study | female | 47 | single | Type1 | China1 | discovery phase of association study, phylogeny | |
| HS055 | Current study | East Asia | NPC-endemic China | Healthy | saliva | NPC case-control study | male | 70 | | | | discovery phase of association study | |
| HS056 | Current study | East Asia | NPC-endemic China | Healthy | saliva | NPC case-control study | female | 45 | | | | discovery phase of association study | |
| HS057 | Current study | East Asia | NPC-endemic China | Healthy | saliva | NPC case-control study | female | 58 | single | Type1 | China1 | discovery phase of association study, phylogeny | |
| NHS001 | Current study | East Asia | NPC-non-endemic China | Healthy | saliva | SYSUCC | male | 26 | | | | | |
| NHS002 | Current study | East Asia | NPC-non-endemic China | Healthy | saliva | SYSUCC | male | 58 | single | Type1 | China1 | phylogeny | |
| NHS003 | Current study | East Asia | NPC-non-endemic China | Healthy | saliva | SYSUCC | female | 50 | | | | | |
| NHS004 | Current study | East Asia | NPC-non-endemic China | Healthy | saliva | SYSUCC | female | 32 | single | Type1 | China1 | phylogeny | |
| NKLT001 | Current study | East Asia | NPC-non-endemic China | NK/T cell lymphoma | tumor biopsy | SYSUCC | female | 49 | | | | | |
| NKLT002 | Current study | East Asia | NPC-endemic China | NK/T cell lymphoma | tumor biopsy | SYSUCC | male | 48 | single | Type1 | China1 | phylogeny | |
| NKLT003-2 | Current study | East Asia | NPC-endemic China | NK/T cell lymphoma | tumor biopsy | SYSUCC | male | 46 | single | Type1 | China1 | phylogeny | |
| NKLT004 | Current study | East Asia | NPC-endemic China | NK/T cell lymphoma | tumor biopsy | SYSUCC | male | 49 | single | Type1 | China1 | phylogeny | |
| NKLT005 | Current study | East Asia | NPC-non-endemic China | NK/T cell lymphoma | tumor biopsy | SYSUCC | male | 21 | | | | | |
| NKLT006 | Current study | East Asia | NPC-endemic China | NK/T cell lymphoma | tumor biopsy | SYSUCC | male | 24 | single | Type1 | China1 | phylogeny | |
| NKLT007 | Current study | East Asia | NPC-endemic China | NK/T cell lymphoma | tumor biopsy | SYSUCC | male | 48 | single | Type1 | China1 | phylogeny | |
| NNPCT001 | Current study | East Asia | NPC-non-endemic China | NPC | tumor biopsy | AHQU | male | 46 | single | Type1 | China1 | phylogeny | |
| NNPCT002 | Current study | East Asia | NPC-non-endemic China | NPC | tumor biopsy | AHQU | male | 49 | single | Type1 | China1 | phylogeny | |
| NNPCT003 | Current study | East Asia | NPC-non-endemic China | NPC | tumor biopsy | AHQU | male | 45 | single | Type1 | China1 | phylogeny | |
| NNPCT004 | Current study | East Asia | NPC-non-endemic China | NPC | tumor biopsy | AHQU | female | 46 | single | Type1 | China1 | phylogeny | |
| NNPCT005 | Current study | East Asia | NPC-non-endemic China | NPC | tumor biopsy | AHQU | male | 17 | single | Type1 | China1 | phylogeny | |

**Supplementary Table 1 List and summary of 270 EBV isolates sequenced in current study and 97 publicly accessed genomes included in the analysis**

**(a) Discription of 270 EBV isolates sequenced in current study and 97 publicly accessed genomes**

| Sample ID | Sequenced by | Geographic origin | Detailed geographic origin | Phenotype | Sample type | Recruitment | Sex | Age | Strain | EBV Type | LMP1 classification | Analysis | Notes |
|---|---|---|---|---|---|---|---|---|---|---|---|---|---|
| NNPCT006 | Current study | East Asia | NPC-non-endemic China | NPC | tumor biopsy | AHQU | female | 40 | | | | | |
| NPCP001 | Current study | East Asia | NPC-non-endemic China | NPC | plasma | SYSUCC | male | 41 | single | Type1 | China1 | phylogeny | NPCP001 and NPCT001 were from the same patient. |
| NPCS001 | Current study | East Asia | NPC-endemic China | NPC | saliva | NPC case-control study | male | 37 | single | Type1 | China1 | discovery phase of association study, phylogeny | |
| NPCS002 | Current study | East Asia | NPC-endemic China | NPC | saliva | NPC case-control study | female | 33 | single | Type1 | China1 | discovery phase of association study, phylogeny | |
| NPCS003-2 | Current study | East Asia | NPC-endemic China | NPC | saliva | NPC case-control study | male | 48 | single | Type1 | China1 | discovery phase of association study, phylogeny | |
| NPCS004 | Current study | East Asia | NPC-endemic China | NPC | saliva | NPC case-control study | male | 49 | | | | discovery phase of association study | |
| NPCS005 | Current study | East Asia | NPC-endemic China | NPC | saliva | NPC case-control study | female | 27 | single | Type1 | China1 | discovery phase of association study, phylogeny | |
| NPCS006 | Current study | East Asia | NPC-endemic China | NPC | saliva | NPC case-control study | male | 45 | single | Type1 | China1 | discovery phase of association study, phylogeny | |
| NPCS007 | Current study | East Asia | NPC-endemic China | NPC | saliva | NPC case-control study | male | 55 | single | Type1 | China1 | discovery phase of association study, phylogeny | |
| NPCS008 | Current study | East Asia | NPC-endemic China | NPC | saliva | NPC case-control study | male | 69 | single | Type1 | China1 | discovery phase of association study, phylogeny | |
| NPCS009 | Current study | East Asia | NPC-endemic China | NPC | saliva | NPC case-control study | female | 73 | single | Type1 | China1 | discovery phase of association study, phylogeny | |
| NPCS010 | Current study | East Asia | NPC-endemic China | NPC | saliva | NPC case-control study | male | 35 | single | Type1 | China1 | discovery phase of association study, phylogeny | |
| NPCS011 | Current study | East Asia | NPC-endemic China | NPC | saliva | NPC case-control study | female | 52 | single | Type1 | China1 | discovery phase of association study, phylogeny | |
| NPCS012 | Current study | East Asia | NPC-endemic China | NPC | saliva | NPC case-control study | male | 47 | single | Type1 | China1 | discovery phase of association study, phylogeny | |
| NPCS013 | Current study | East Asia | NPC-endemic China | NPC | saliva | NPC case-control study | male | 56 | single | Type1 | China2 | discovery phase of association study, phylogeny | |
| NPCS014 | Current study | East Asia | NPC-endemic China | NPC | saliva | NPC case-control study | female | 63 | single | Type1 | China1 | discovery phase of association study, phylogeny | |
| NPCS015 | Current study | East Asia | NPC-endemic China | NPC | saliva | NPC case-control study | male | 72 | | | | discovery phase of association study | |
| NPCS016 | Current study | East Asia | NPC-endemic China | NPC | saliva | NPC case-control study | female | 46 | single | Type1 | China1 | discovery phase of association study, phylogeny | |
| NPCS017 | Current study | East Asia | NPC-endemic China | NPC | saliva | NPC case-control study | male | 53 | single | Type1 | China1 | discovery phase of association study, phylogeny | |
| NPCS018 | Current study | East Asia | NPC-endemic China | NPC | saliva | NPC case-control study | male | 53 | single | Type1 | China1 | discovery phase of association study, phylogeny | |
| NPCS019 | Current study | East Asia | NPC-endemic China | NPC | saliva | NPC case-control study | male | 59 | single | Type1 | China1 | discovery phase of association study, phylogeny | |
| NPCS021 | Current study | East Asia | NPC-endemic China | NPC | saliva | NPC case-control study | male | 30 | single | Type1 | China1 | discovery phase of association study, phylogeny | |
| NPCS022 | Current study | East Asia | NPC-endemic China | NPC | saliva | NPC case-control study | female | 43 | single | Type1 | China1 | discovery phase of association study, phylogeny | |
| NPCS023 | Current study | East Asia | NPC-endemic China | NPC | saliva | NPC case-control study | male | 38 | single | Type1 | China1 | discovery phase of association study, phylogeny | |

**Supplementary Table 1 List and summary of 270 EBV isolates sequenced in current study and 97 publicly accessed genomes included in the analysis**

**(a) Discription of 270 EBV isolates sequenced in current study and 97 publicly accessed genomes**

| Sample ID | Sequenced by | Geographic origin | Detailed geographic origin | Phenotype | Sample type | Recruitment | Sex | Age | Strain | EBV Type | LMP1 classification | Analysis | Notes |
|---|---|---|---|---|---|---|---|---|---|---|---|---|---|
| NPCS024 | Current study | East Asia | NPC-endemic China | NPC | saliva | NPC case-control study | male | 48 | single | Type1 | China1 | discovery phase of association study, phylogeny | |
| NPCS025 | Current study | East Asia | NPC-endemic China | NPC | saliva | NPC case-control study | male | 42 | single | Type1 | China1 | discovery phase of association study, phylogeny | |
| NPCS026 | Current study | East Asia | NPC-endemic China | NPC | saliva | NPC case-control study | male | 50 | single | Type1 | China1 | discovery phase of association study, phylogeny | |
| NPCS027 | Current study | East Asia | NPC-endemic China | NPC | saliva | NPC case-control study | male | 66 | single | Type1 | China1 | discovery phase of association study, phylogeny | |
| NPCS028 | Current study | East Asia | NPC-endemic China | NPC | saliva | NPC case-control study | male | 43 | single | Type1 | China1 | discovery phase of association study, phylogeny | |
| NPCS029 | Current study | East Asia | NPC-endemic China | NPC | saliva | NPC case-control study | female | 46 | single | Type1 | China1 | discovery phase of association study, phylogeny | |
| NPCS030 | Current study | East Asia | NPC-endemic China | NPC | saliva | NPC case-control study | male | 50 | single | Type1 | China1 | discovery phase of association study, phylogeny | |
| NPCS031 | Current study | East Asia | NPC-endemic China | NPC | saliva | NPC case-control study | female | 58 | single | Type1 | China1 | discovery phase of association study, phylogeny | |
| NPCS032 | Current study | East Asia | NPC-endemic China | NPC | saliva | NPC case-control study | female | 33 | | | | discovery phase of association study | |
| NPCS033 | Current study | East Asia | NPC-endemic China | NPC | saliva | NPC case-control study | female | 67 | single | Type1 | China1 | discovery phase of association study, phylogeny | |
| NPCS034 | Current study | East Asia | NPC-endemic China | NPC | saliva | NPC case-control study | male | 62 | single | Type1 | China1 | discovery phase of association study, phylogeny | |
| NPCS035 | Current study | East Asia | NPC-endemic China | NPC | saliva | NPC case-control study | male | 35 | single | Type1 | China1 | discovery phase of association study, phylogeny | |
| NPCS036 | Current study | East Asia | NPC-endemic China | NPC | saliva | NPC case-control study | male | 38 | | | | discovery phase of association study | |
| NPCS037 | Current study | East Asia | NPC-endemic China | NPC | saliva | NPC case-control study | male | 50 | | | | discovery phase of association study | |
| NPCS038 | Current study | East Asia | NPC-endemic China | NPC | saliva | NPC case-control study | female | 65 | single | Type1 | China1 | discovery phase of association study, phylogeny | |
| NPCS039 | Current study | East Asia | NPC-endemic China | NPC | saliva | NPC case-control study | male | 62 | single | Type1 | China1 | discovery phase of association study, phylogeny | |
| NPCS040 | Current study | East Asia | NPC-endemic China | NPC | saliva | NPC case-control study | male | 49 | single | Type1 | China1 | discovery phase of association study, phylogeny | |
| NPCS041 | Current study | East Asia | NPC-endemic China | NPC | saliva | NPC case-control study | male | 53 | | | | discovery phase of association study | |
| NPCS042 | Current study | East Asia | NPC-endemic China | NPC | saliva | NPC case-control study | female | 58 | single | Type1 | China1 | discovery phase of association study, phylogeny | |
| NPCS043 | Current study | East Asia | NPC-endemic China | NPC | saliva | NPC case-control study | male | 55 | | | | discovery phase of association study | |
| NPCS044 | Current study | East Asia | NPC-endemic China | NPC | saliva | NPC case-control study | male | 67 | single | Type1 | China1 | discovery phase of association study, phylogeny | |
| NPCS045 | Current study | East Asia | NPC-endemic China | NPC | saliva | NPC case-control study | male | 51 | single | Type1 | China1 | discovery phase of association study, phylogeny | |
| NPCS046 | Current study | East Asia | NPC-endemic China | NPC | saliva | NPC case-control study | female | 51 | single | Type1 | China1 | discovery phase of association study, phylogeny | |
| NPCS047 | Current study | East Asia | NPC-endemic China | NPC | saliva | NPC case-control study | male | 37 | single | Type1 | China1 | discovery phase of association study, phylogeny | |

**Supplementary Table 1 List and summary of 270 EBV isolates sequenced in current study and 97 publicly accessed genomes included in the analysis**

**(a) Discription of 270 EBV isolates sequenced in current study and 97 publicly accessed genomes**

| Sample ID | Sequenced by | Geographic origin | Detailed geographic origin | Phenotype | Sample type | Recruitment | Sex | Age | Strain | EBV Type | LMP1 classification | Analysis | Notes |
|---|---|---|---|---|---|---|---|---|---|---|---|---|---|
| NPCS048 | Current study | East Asia | NPC-endemic China | NPC | saliva | NPC case-control study | male | 58 | single | Type1 | China1 | discovery phase of association study, phylogeny | |
| NPCS049 | Current study | East Asia | NPC-endemic China | NPC | saliva | NPC case-control study | male | 77 | single | Type1 | China1 | discovery phase of association study, phylogeny | |
| NPCS050 | Current study | East Asia | NPC-endemic China | NPC | saliva | NPC case-control study | male | 31 | single | Type1 | China1 | discovery phase of association study, phylogeny | |
| NPCS051 | Current study | East Asia | NPC-endemic China | NPC | saliva | NPC case-control study | male | 44 | single | Type1 | China1 | discovery phase of association study, phylogeny | |
| NPCS052 | Current study | East Asia | NPC-endemic China | NPC | saliva | NPC case-control study | male | 66 | single | Type1 | China1 | discovery phase of association study, phylogeny | |
| NPCS054 | Current study | East Asia | NPC-endemic China | NPC | saliva | NPC case-control study | male | 50 | single | Type1 | China1 | discovery phase of association study, phylogeny | |
| NPCS055 | Current study | East Asia | NPC-endemic China | NPC | saliva | NPC case-control study | male | 60 | | | | discovery phase of association study | |
| NPCT001 | Current study | East Asia | NPC-non-endemic China | NPC | tumor biopsy | SYSUCC | male | 41 | single | Type1 | China1 | phylogeny | |
| NPCT002 | Current study | East Asia | NPC-endemic China | NPC | tumor biopsy | SYSUCC | male | 51 | single | Type1 | China1 | discovery phase of association study, phylogeny | |
| NPCT003 | Current study | East Asia | NPC-endemic China | NPC | tumor biopsy | SYSUCC | male | 44 | single | Type1 | China1 | discovery phase of association study, phylogeny | |
| NPCT004 | Current study | East Asia | NPC-endemic China | NPC | tumor biopsy | SYSUCC | male | 35 | single | Type1 | China1 | discovery phase of association study, phylogeny | |
| NPCT005 | Current study | East Asia | NPC-endemic China | NPC | tumor biopsy | SYSUCC | male | 49 | single | Type1 | China1 | discovery phase of association study, phylogeny | |
| NPCT006 | Current study | East Asia | NPC-endemic China | NPC | tumor biopsy | SYSUCC | male | 54 | single | Type1 | China1 | discovery phase of association study, phylogeny | |
| NPCT007 | Current study | East Asia | NPC-endemic China | NPC | tumor biopsy | SYSUCC | male | 49 | single | Type1 | China1 | discovery phase of association study, phylogeny | |
| NPCT008 | Current study | East Asia | NPC-non-endemic China | NPC | tumor biopsy | SYSUCC | male | 73 | single | Type1 | China1 | phylogeny | |
| NPCT009 | Current study | East Asia | NPC-endemic China | NPC | tumor biopsy | SYSUCC | male | 58 | single | Type1 | China1 | discovery phase of association study, phylogeny | |
| NPCT010 | Current study | East Asia | NPC-endemic China | NPC | tumor biopsy | SYSUCC | male | 51 | single | Type1 | China1 | discovery phase of association study, phylogeny | |
| NPCT011 | Current study | East Asia | NPC-non-endemic China | NPC | tumor biopsy | SYSUCC | male | 71 | single | Type1 | China1 | phylogeny | |
| NPCT012 | Current study | East Asia | NPC-endemic China | NPC | tumor biopsy | SYSUCC | female | 43 | single | Type1 | China1 | discovery phase of association study, phylogeny | |
| NPCT013 | Current study | East Asia | NPC-endemic China | NPC | tumor biopsy | SYSUCC | male | 34 | single | Type1 | China1 | discovery phase of association study, phylogeny | |
| NPCT014 | Current study | East Asia | NPC-endemic China | NPC | tumor biopsy | FAHGMU | male | 27 | single | Type1 | China1 | discovery phase of association study, phylogeny | |
| NPCT015 | Current study | East Asia | NPC-endemic China | NPC | tumor biopsy | FAHGMU | male | 69 | single | Type1 | China1 | discovery phase of association study, phylogeny | |
| NPCT016 | Current study | East Asia | NPC-endemic China | NPC | tumor biopsy | FAHGMU | male | 62 | | | | discovery phase of association study | |
| NPCT017 | Current study | East Asia | NPC-endemic China | NPC | tumor biopsy | FAHGMU | male | 43 | single | Type1 | China1 | discovery phase of association study, phylogeny | |

**Supplementary Table 1 List and summary of 270 EBV isolates sequenced in current study and 97 publicly accessed genomes included in the analysis**

**(a) Discription of 270 EBV isolates sequenced in current study and 97 publicly accessed genomes**

| Sample ID | Sequenced by | Geographic origin | Detailed geographic origin | Phenotype | Sample type | Recruitment | Sex | Age | Strain | EBV Type | LMP1 classification | Analysis | Notes |
|---|---|---|---|---|---|---|---|---|---|---|---|---|---|
| NPCT018 | Current study | East Asia | NPC-endemic China | NPC | tumor biopsy | FAHGMU | male | 49 | single | Type1 | China1 | discovery phase of association study, phylogeny | |
| NPCT019 | Current study | East Asia | NPC-endemic China | NPC | tumor biopsy | FAHGMU | male | 43 | single | Type1 | China1 | discovery phase of association study, phylogeny | |
| NPCT020-2 | Current study | East Asia | NPC-endemic China | NPC | tumor biopsy | FAHGMU | male | 69 | single | Type1 | China1 | discovery phase of association study, phylogeny | |
| NPCT021 | Current study | East Asia | NPC-endemic China | NPC | tumor biopsy | FAHGMU | male | 66 | single | Type1 | China2 | discovery phase of association study, phylogeny | |
| NPCT022 | Current study | East Asia | NPC-endemic China | NPC | tumor biopsy | FAHGMU | male | 72 | single | Type1 | China1 | discovery phase of association study, phylogeny | |
| NPCT023 | Current study | East Asia | NPC-endemic China | NPC | tumor biopsy | FAHGMU | male | 47 | single | Type1 | China1 | discovery phase of association study, phylogeny | |
| NPCT024 | Current study | East Asia | NPC-endemic China | NPC | tumor biopsy | FAHGMU | male | 50 | single | Type1 | China1 | discovery phase of association study, phylogeny | |
| NPCT025 | Current study | East Asia | NPC-endemic China | NPC | tumor biopsy | FAHGMU | male | 66 | single | Type1 | China1 | discovery phase of association study, phylogeny | |
| NPCT026 | Current study | East Asia | NPC-endemic China | NPC | tumor biopsy | FAHGMU | male | 31 | | | | discovery phase of association study | |
| NPCT027 | Current study | East Asia | NPC-endemic China | NPC | tumor biopsy | FAHGMU | male | 56 | single | Type1 | China1 | discovery phase of association study, phylogeny | |
| NPCT028-2 | Current study | East Asia | NPC-endemic China | NPC | tumor biopsy | FAHGMU | male | 53 | single | Type1 | China1 | discovery phase of association study, phylogeny | |
| NPCT029 | Current study | East Asia | NPC-endemic China | NPC | tumor biopsy | FAHGMU | male | 46 | single | Type1 | China1 | discovery phase of association study, phylogeny | |
| NPCT031 | Current study | East Asia | NPC-endemic China | NPC | tumor biopsy | FAHGMU | female | 47 | single | Type1 | China1 | discovery phase of association study, phylogeny | |
| NPCT032 | Current study | East Asia | NPC-endemic China | NPC | tumor biopsy | FAHGMU | female | 44 | single | Type1 | China1 | discovery phase of association study, phylogeny | |
| NPCT033 | Current study | East Asia | NPC-endemic China | NPC | tumor biopsy | FAHGMU | male | 51 | single | Type1 | China1 | discovery phase of association study, phylogeny | |
| NPCT035 | Current study | East Asia | NPC-endemic China | NPC | tumor biopsy | FAHGMU | male | 57 | single | Type1 | China1 | discovery phase of association study, phylogeny | |
| NPCT036 | Current study | East Asia | NPC-endemic China | NPC | tumor biopsy | FAHGMU | female | 48 | single | Type1 | China1 | discovery phase of association study, phylogeny | |
| NPCT037 | Current study | East Asia | NPC-endemic China | NPC | tumor biopsy | FAHGMU | male | 58 | single | Type1 | China1 | discovery phase of association study, phylogeny | |
| NPCT038 | Current study | East Asia | NPC-endemic China | NPC | tumor biopsy | FAHGMU | female | 57 | single | Type1 | China1 | discovery phase of association study, phylogeny | |
| NPCT039 | Current study | East Asia | NPC-endemic China | NPC | tumor biopsy | FAHGMU | male | 37 | single | Type1 | China1 | discovery phase of association study, phylogeny | |
| NPCT040 | Current study | East Asia | NPC-endemic China | NPC | tumor biopsy | FAHGMU | male | 28 | single | Type1 | China1 | discovery phase of association study, phylogeny | |
| NPCT041 | Current study | East Asia | NPC-endemic China | NPC | tumor biopsy | FAHGMU | male | 49 | single | Type1 | China1 | discovery phase of association study, phylogeny | |
| NPCT042 | Current study | East Asia | NPC-endemic China | NPC | tumor biopsy | FAHGMU | male | 65 | single | Type1 | China1 | discovery phase of association study, phylogeny | |
| NPCT043 | Current study | East Asia | NPC-endemic China | NPC | tumor biopsy | FAHGMU | male | 42 | single | Type1 | China1 | discovery phase of association study, phylogeny | |

**Supplementary Table 1 List and summary of 270 EBV isolates sequenced in current study and 97 publicly accessed genomes included in the analysis**

**(a) Discription of 270 EBV isolates sequenced in current study and 97 publicly accessed genomes**

| Sample ID | Sequenced by | Geographic origin | Detailed geographic origin | Phenotype | Sample type | Recruitment | Sex | Age | Strain | EBV Type | LMP1 classification | Analysis | Notes |
|---|---|---|---|---|---|---|---|---|---|---|---|---|---|
| NPCT045 | Current study | East Asia | NPC-endemic China | NPC | tumor biopsy | SYSUCC | female | 38 | single | Type1 | China1 | discovery phase of association study, phylogeny | |
| NPCT046 | Current study | East Asia | NPC-non-endemic China | NPC | tumor biopsy | SYSUCC | female | 52 | single | Type1 | China2 | phylogeny | |
| NPCT047 | Current study | East Asia | NPC-endemic China | NPC | tumor biopsy | SYSUCC | female | 62 | single | Type1 | China1 | discovery phase of association study, phylogeny | |
| NPCT048 | Current study | East Asia | NPC-endemic China | NPC | tumor biopsy | SYSUCC | female | 39 | single | Type1 | China1 | discovery phase of association study, phylogeny | |
| NPCT049 | Current study | East Asia | NPC-endemic China | NPC | tumor biopsy | FAHGMU | male | 57 | single | Type1 | China2 | discovery phase of association study, phylogeny | |
| NPCT050 | Current study | East Asia | NPC-endemic China | NPC | tumor biopsy | FAHGMU | male | 38 | single | Type1 | China1 | discovery phase of association study, phylogeny | |
| NPCT051 | Current study | East Asia | NPC-endemic China | NPC | tumor biopsy | FAHGMU | male | 54 | | | | discovery phase of association study | |
| NPCT052 | Current study | East Asia | NPC-endemic China | NPC | tumor biopsy | FAHGMU | male | 36 | single | Type1 | China1 | discovery phase of association study, phylogeny | |
| NPCT053 | Current study | East Asia | NPC-endemic China | NPC | tumor biopsy | FAHGMU | male | 63 | single | Type1 | China1 | discovery phase of association study, phylogeny | |
| NPCT054 | Current study | East Asia | NPC-endemic China | NPC | tumor biopsy | SYSUCC | male | 57 | single | Type1 | China1 | discovery phase of association study, phylogeny | NPCT054 and NPCT054M were from the same patient. NPCT054 is from primary tumor; NPCT054M was from metastatic tumor. |
| NPCT054M | Current study | East Asia | NPC-endemic China | NPC | tumor biopsy | SYSUCC | male | 57 | single | Type1 | China1 | phylogeny | |
| NPCT055 | Current study | East Asia | NPC-non-endemic China | NPC | tumor biopsy | SYSUCC | female | 34 | single | Type1 | China1 | phylogeny | NPCT055 and NPCT055M were from the same patient. NPCT055 is from primary tumor; NPCT055M was from metastatic tumor. |
| NPCT055M | Current study | East Asia | NPC-non-endemic China | NPC | tumor biopsy | SYSUCC | female | 34 | single | Type1 | China1 | phylogeny | |
| NPCT056 | Current study | East Asia | NPC-endemic China | NPC | tumor biopsy | SYSUCC | male | 31 | single | Type1 | China1 | discovery phase of association study, phylogeny | NPCT056 and NPCT056M were from the same patient. NPCT056 is from primary tumor; NPCT056M was from metastatic tumor. |
| NPCT056M | Current study | East Asia | NPC-endemic China | NPC | tumor biopsy | SYSUCC | male | 31 | single | Type1 | China1 | phylogeny | |
| NPCT057 | Current study | East Asia | NPC-non-endemic China | NPC | tumor biopsy | SYSUCC | female | 60 | single | Type1 | China1 | phylogeny | NPCT057 and NPCT057M were from the same patient. NPCT057 is from primary tumor; NPCT057M was from metastatic tumor. |
| NPCT057M | Current study | East Asia | NPC-non-endemic China | NPC | tumor biopsy | SYSUCC | female | 60 | single | Type1 | China1 | phylogeny | |
| NPCT058 | Current study | East Asia | NPC-non-endemic China | NPC | tumor biopsy | SYSUCC | male | 44 | single | Type1 | China1 | phylogeny | NPCT058 and NPCT058M were from the same patient. NPCT058 is from primary tumor; NPCT058M was from metastatic tumor. |
| NPCT058M | Current study | East Asia | NPC-non-endemic China | NPC | tumor biopsy | SYSUCC | male | 44 | single | Type1 | China1 | phylogeny | |
| NPCT059 | Current study | East Asia | NPC-endemic China | NPC | tumor biopsy | SYSUCC | male | 49 | single | Type1 | China1 | discovery phase of association study, phylogeny | |
| NPCT060 | Current study | East Asia | NPC-endemic China | NPC | tumor biopsy | SYSUCC | male | 61 | single | Type1 | Med | discovery phase of association study, phylogeny | |
| NPCT061 | Current study | East Asia | NPC-endemic China | NPC | tumor biopsy | SYSUCC | male | 28 | single | Type1 | China1 | discovery phase of association study, phylogeny | |
| NPCT062 | Current study | East Asia | NPC-endemic China | NPC | tumor biopsy | SYSUCC | male | 35 | single | Type1 | China1 | discovery phase of association study, phylogeny | |
| NPCT063 | Current study | East Asia | NPC-endemic China | NPC | tumor biopsy | SYSUCC | female | 47 | single | Type1 | China1 | discovery phase of association study, phylogeny | |

**Supplementary Table 1 List and summary of 270 EBV isolates sequenced in current study and 97 publicly accessed genomes included in the analysis**

**(a) Discription of 270 EBV isolates sequenced in current study and 97 publicly accessed genomes**

| Sample ID | Sequenced by | Geographic origin | Detailed geographic origin | Phenotype | Sample type | Recruitment | Sex | Age | Strain | EBV Type | LMP1 classification | Analysis | Notes |
|---|---|---|---|---|---|---|---|---|---|---|---|---|---|
| NPCT064 | Current study | East Asia | NPC-endemic China | NPC | tumor biopsy | SYSUCC | male | 31 | single | Type1 | China1 | discovery phase of association study, phylogeny | |
| NPCT065 | Current study | East Asia | NPC-endemic China | NPC | tumor biopsy | SYSUCC | male | 44 | single | Type1 | China1 | discovery phase of association study, phylogeny | |
| NPCT066 | Current study | East Asia | NPC-endemic China | NPC | tumor biopsy | SYSUCC | male | 38 | single | Type1 | China1 | discovery phase of association study, phylogeny | |
| NPCT067 | Current study | East Asia | NPC-endemic China | NPC | tumor biopsy | SYSUCC | female | 49 | single | Type1 | China1 | discovery phase of association study, phylogeny | |
| NPCT068 | Current study | East Asia | NPC-endemic China | NPC | tumor biopsy | SYSUCC | male | 42 | single | Type1 | China1 | discovery phase of association study, phylogeny | |
| NPCT069 | Current study | East Asia | NPC-endemic China | NPC | tumor biopsy | SYSUCC | male | 46 | single | Type1 | China1 | discovery phase of association study, phylogeny | |
| NPCT070 | Current study | East Asia | NPC-endemic China | NPC | tumor biopsy | SYSUCC | male | 31 | single | Type1 | China1 | discovery phase of association study, phylogeny | |
| NPCT071 | Current study | East Asia | NPC-endemic China | NPC | tumor biopsy | SYSUCC | female | 60 | single | Type1 | China1 | discovery phase of association study, phylogeny | |
| NPCT072 | Current study | East Asia | NPC-endemic China | NPC | tumor biopsy | SYSUCC | male | 51 | single | Type1 | China1 | discovery phase of association study, phylogeny | |
| NPCT073 | Current study | East Asia | NPC-endemic China | NPC | tumor biopsy | SYSUCC | female | 44 | single | Type1 | China1 | discovery phase of association study, phylogeny | |
| NPCT074 | Current study | East Asia | NPC-endemic China | NPC | tumor biopsy | SYSUCC | female | 66 | single | Type1 | China1 | phylogeny | NPCT074 and NPCT074S were from the same patient. |
| NPCT074S | Current study | East Asia | NPC-endemic China | NPC | saliva | SYSUCC | female | 66 | single | Type1 | China1 | discovery phase of association study, phylogeny | |
| NPCT075 | Current study | East Asia | NPC-endemic China | NPC | tumor biopsy | SYSUCC | male | 32 | single | Type1 | China1 | discovery phase of association study, phylogeny | |
| NPCT076 | Current study | East Asia | NPC-endemic China | NPC | tumor biopsy | SYSUCC | female | 43 | single | Type1 | China1 | discovery phase of association study, phylogeny | |
| NPCT077 | Current study | East Asia | NPC-endemic China | NPC | tumor biopsy | SYSUCC | male | 47 | single | Type1 | China1 | discovery phase of association study, phylogeny | |
| NPCT078 | Current study | East Asia | NPC-non-endemic China | NPC | tumor biopsy | SYSUCC | male | 39 | single | Type1 | China1 | phylogeny | |
| NPCT079 | Current study | East Asia | NPC-non-endemic China | NPC | tumor biopsy | SYSUCC | male | 38 | | | | | |
| NPCT080 | Current study | East Asia | NPC-endemic China | NPC | tumor biopsy | SYSUCC | male | 29 | single | Type1 | China1 | discovery phase of association study, phylogeny | |
| NPCT081 | Current study | East Asia | NPC-endemic China | NPC | tumor biopsy | SYSUCC | female | 45 | single | Type1 | China1 | discovery phase of association study, phylogeny | |
| NPCT082 | Current study | East Asia | NPC-endemic China | NPC | tumor biopsy | SYSUCC | male | 34 | single | Type1 | China1 | discovery phase of association study, phylogeny | |
| NPCT083 | Current study | East Asia | NPC-endemic China | NPC | tumor biopsy | SYSUCC | male | 42 | single | Type1 | China1 | discovery phase of association study, phylogeny | |
| NPCT084 | Current study | East Asia | NPC-endemic China | NPC | tumor biopsy | SYSUCC | male | 45 | single | Type1 | China1 | discovery phase of association study, phylogeny | |
| NPCT085 | Current study | East Asia | NPC-endemic China | NPC | tumor biopsy | SYSUCC | female | 59 | single | Type1 | China1 | discovery phase of association study, phylogeny | |
| NPCT086 | Current study | East Asia | NPC-endemic China | NPC | tumor biopsy | SYSUCC | female | 49 | single | Type1 | China1 | discovery phase of association study, phylogeny | |

**Supplementary Table 1 List and summary of 270 EBV isolates sequenced in current study and 97 publicly accessed genomes included in the analysis**

**(a) Discription of 270 EBV isolates sequenced in current study and 97 publicly accessed genomes**

| Sample ID | Sequenced by | Geographic origin | Detailed geographic origin | Phenotype | Sample type | Recruitment | Sex | Age | Strain | EBV Type | LMP1 classification | Analysis | Notes |
|---|---|---|---|---|---|---|---|---|---|---|---|---|---|
| NPCT087 | Current study | East Asia | NPC-endemic China | NPC | tumor biopsy | SYSUCC | male | 63 | single | Type1 | China1 | discovery phase of association study, phylogeny | |
| NPCT088 | Current study | East Asia | NPC-endemic China | NPC | tumor biopsy | SYSUCC | male | 51 | single | Type1 | China1 | discovery phase of association study, phylogeny | |
| NPCT089 | Current study | East Asia | NPC-endemic China | NPC | tumor biopsy | SYSUCC | female | 24 | single | Type1 | China1 | discovery phase of association study, phylogeny | |
| NPCT090 | Current study | East Asia | NPC-endemic China | NPC | tumor biopsy | SYSUCC | male | 34 | single | Type1 | China1 | discovery phase of association study, phylogeny | |
| NPCT091 | Current study | East Asia | NPC-endemic China | NPC | tumor biopsy | SYSUCC | female | 25 | single | Type1 | China1 | discovery phase of association study, phylogeny | |
| NPCT092 | Current study | East Asia | NPC-endemic China | NPC | tumor biopsy | FAHGMU | male | 30 | single | Type1 | China1 | discovery phase of association study, phylogeny | |
| NPCT093 | Current study | East Asia | NPC-endemic China | NPC | tumor biopsy | FAHGMU | female | 33 | single | Type1 | China1 | discovery phase of association study, phylogeny | |
| NPCT094 | Current study | East Asia | NPC-endemic China | NPC | tumor biopsy | FAHGMU | male | 71 | single | Type1 | China1 | discovery phase of association study, phylogeny | |
| NPCT095 | Current study | East Asia | NPC-endemic China | NPC | tumor biopsy | FAHGMU | female | 42 | | | | discovery phase of association study | |
| NPCT096 | Current study | East Asia | NPC-endemic China | NPC | tumor biopsy | FAHGMU | male | 49 | single | Type1 | China1 | discovery phase of association study, phylogeny | |
| NPCT097 | Current study | East Asia | NPC-endemic China | NPC | tumor biopsy | FAHGMU | male | 43 | | | | discovery phase of association study | |
| NPCT098 | Current study | East Asia | NPC-endemic China | NPC | tumor biopsy | FAHGMU | female | 61 | single | Type1 | China1 | discovery phase of association study, phylogeny | |
| NPCT099 | Current study | East Asia | NPC-endemic China | NPC | tumor biopsy | FAHGMU | male | 41 | single | Type1 | China1 | discovery phase of association study, phylogeny | |
| NPCT100 | Current study | East Asia | NPC-endemic China | NPC | tumor biopsy | FAHGMU | female | 34 | single | Type1 | China1 | discovery phase of association study, phylogeny | |
| NPCT101 | Current study | East Asia | NPC-endemic China | NPC | tumor biopsy | FAHGMU | male | 39 | single | Type1 | China1 | discovery phase of association study, phylogeny | |
| NPCT102 | Current study | East Asia | NPC-endemic China | NPC | tumor biopsy | FAHGMU | male | 59 | single | Type1 | China1 | discovery phase of association study, phylogeny | |
| NPCT103 | Current study | East Asia | NPC-endemic China | NPC | tumor biopsy | FAHGMU | female | 66 | single | Type1 | China1 | discovery phase of association study, phylogeny | |
| NPCT104 | Current study | East Asia | NPC-endemic China | NPC | tumor biopsy | FAHGMU | male | 33 | single | Type1 | China1 | discovery phase of association study, phylogeny | |
| NPCT105 | Current study | East Asia | NPC-endemic China | NPC | tumor biopsy | FAHGMU | male | 56 | single | Type1 | China1 | discovery phase of association study, phylogeny | |
| NPCT106 | Current study | East Asia | NPC-endemic China | NPC | tumor biopsy | FAHGMU | male | 62 | single | Type1 | China1 | discovery phase of association study, phylogeny | |
| NPCT107 | Current study | East Asia | NPC-endemic China | NPC | tumor biopsy | FAHGMU | female | 49 | single | Type1 | China1 | discovery phase of association study, phylogeny | |
| NPCT108 | Current study | East Asia | NPC-non-endemic China | NPC | tumor biopsy | FAHGMU | male | 47 | single | Type1 | China1 | phylogeny | |
| NPCT109 | Current study | East Asia | NPC-endemic China | NPC | tumor biopsy | FAHGMU | female | 43 | single | Type1 | China1 | discovery phase of association study, phylogeny | |
| NPCT110 | Current study | East Asia | NPC-endemic China | NPC | tumor biopsy | FAHGMU | female | 42 | single | Type1 | China1 | discovery phase of association study, phylogeny | |

**Supplementary Table 1 List and summary of 270 EBV isolates sequenced in current study and 97 publicly accessed genomes included in the analysis**

**(a) Discription of 270 EBV isolates sequenced in current study and 97 publicly accessed genomes**

| Sample ID | Sequenced by | Geographic origin | Detailed geographic origin | Phenotype | Sample type | Recruitment | Sex | Age | Strain | EBV Type | LMP1 classification | Analysis | Notes |
|---|---|---|---|---|---|---|---|---|---|---|---|---|---|
| NPCT111 | Current study | East Asia | NPC-endemic China | NPC | tumor biopsy | FAHGMU | male | 48 | single | Type1 | China1 | discovery phase of association study, phylogeny | |
| NPCT112 | Current study | East Asia | NPC-endemic China | NPC | tumor biopsy | FAHGMU | male | 42 | single | Type1 | China1 | discovery phase of association study, phylogeny | |
| NPCT113 | Current study | East Asia | NPC-endemic China | NPC | tumor biopsy | FAHGMU | male | 29 | single | Type1 | China1 | discovery phase of association study, phylogeny | |
| NPCT114 | Current study | East Asia | NPC-endemic China | NPC | tumor biopsy | FAHGMU | female | 51 | single | Type1 | China1 | discovery phase of association study, phylogeny | |
| NPCT115 | Current study | East Asia | NPC-endemic China | NPC | tumor biopsy | FAHGMU | male | 47 | single | Type1 | China1 | discovery phase of association study, phylogeny | |
| NPCT116 | Current study | East Asia | NPC-endemic China | NPC | tumor biopsy | FAHGMU | male | 46 | | | | discovery phase of association study | |
| LN827544.1 | Palser et al., J Virol, 2015 | Papua New Guinea | Papua New Guinea | Burkitt's lymphoma | cell line | | | | single | Type2 | na | phylogeny | |
| KF717093.1 | Anne,W., Tobias,S. and Wolfgang,H., etc | Africa | Nigeria | Burkitt's lymphoma | cell line | | | | single | Type1 | na | phylogeny | Raji |
| LN827580.1 | Palser et al., J Virol, 2015 | Africa | Kenya | sLCL | | | | | single | Type2 | B95_8 | phylogeny | |
| LN827556.1 | Palser et al., J Virol, 2015 | Africa | Kenya | Burkitt's lymphoma | cell line | | | | single | Type2 | B95_8 | phylogeny | |
| LN827548.2 | Palser et al., J Virol, 2015 | Africa | Nigeria | Burkitt's lymphoma | cell line | | | | single | Type2 | B95_8 | phylogeny | |
| LN827557.2 | Palser et al., J Virol, 2015 | Africa | North Africa | sLCL-BL | | | | | single | Type2 | B95_8 | phylogeny | |
| LN827800.1 | Palser et al., J Virol, 2015 | Africa | Nigeria | Burkitt's lymphoma | cell line | | | | single | Type2 | B95_8 | phylogeny | |
| LN827563.2 | Palser et al., J Virol, 2015 | Africa | Kenya | sLCL | | | | | single | Type2 | China1 | phylogeny | |
| LN827591.1 | Palser et al., J Virol, 2015 | Africa | Kenya | sLCL | | | | | single | Type2 | China1 | phylogeny | |
| LN827554.1 | Palser et al., J Virol, 2015 | Africa | Unknown | LCL | | | | | single | Type2 | China1 | phylogeny | |
| NC_009334.1 | Dolan A et al., J Virol, 2006 | Africa | Ghana | Burkitt's lymphoma | cell line | | | | single | Type2 | na | phylogeny | AG876 |
| LN831023.1 | Palser et al., J Virol, 2015 | Africa | Kenya | sLCL | | | | | single | Type2 | na | phylogeny | |
| LN827560.1 | Palser et al., J Virol, 2015 | Africa | Kenya | sLCL | | | | | single | Type2 | na | phylogeny | |
| LN827587.1 | Palser et al., J Virol, 2015 | Africa | Kenya | sLCL | | | | | single | Type2 | na | phylogeny | |
| LN827562.1 | Palser et al., J Virol, 2015 | Africa | Kenya | sLCL | | | | | single | Type1 | na | phylogeny | |
| LN824203.1 | Palser et al., J Virol, 2015 | Africa | Kenya | Burkitt's lymphoma | cell line | | | | single | Type1 | na | phylogeny | |
| LN827551.1 | Palser et al., J Virol, 2015 | Africa | Kenya | Burkitt's lymphoma | cell line | | | | single | Type1 | na | phylogeny | |
| LN827526.1 | Palser et al., J Virol, 2015 | Africa | Africa | Burkitt's lymphoma | cell line | | | | single | Type1 | na | phylogeny | |

**Supplementary Table 1 List and summary of 270 EBV isolates sequenced in current study and 97 publicly accessed genomes included in the analysis**

**(a) Discription of 270 EBV isolates sequenced in current study and 97 publicly accessed genomes**

| Sample ID | Sequenced by | Geographic origin | Detailed geographic origin | Phenotype | Sample type | Recruitment | Sex | Age | Strain | EBV Type | LMP1 classification | Analysis | Notes |
|---|---|---|---|---|---|---|---|---|---|---|---|---|---|
| LN827527.1 | Palser et al., J Virol, 2015 | Africa | North Africa | LCL | | | | | single | Type1 | B95_8 | phylogeny | |
| LN827574.1 | Palser et al., J Virol, 2015 | Africa | Kenya | sLCL | | | | | single | Type1 | Med | phylogeny | |
| NA19384 | Santpere et al., Genome Biol Evol, 2014 | Africa | Kenya | LCL | | | | | single | Type1 | Med | phylogeny | |
| LN827568.1 | Palser et al., J Virol, 2015 | Africa | Kenya | sLCL | | | | | single | Type1 | Med | phylogeny | |
| LN827579.1 | Palser et al., J Virol, 2015 | Africa | Kenya | sLCL | | | | | single | Type1 | Med | phylogeny | |
| LN827581.1 | Palser et al., J Virol, 2015 | Africa | Kenya | sLCL | | | | | single | Type1 | Med | phylogeny | |
| LN824205.1 | Palser et al., J Virol, 2015 | Africa | Kenya | sLCL | | | | | single | Type1 | Med | phylogeny | |
| LN827545.1 | Palser et al., J Virol, 2015 | Africa | Kenya | Burkitt's lymphoma | cell line | | | | single | Type1 | Med | phylogeny | |
| NA19114 | Santpere et al., Genome Biol Evol, 2014 | Africa | Yoruba | LCL | | | | | single | Type1 | Med | phylogeny | |
| LN827577.1 | Palser et al., J Virol, 2015 | Africa | Kenya | sLCL | | | | | single | Type1 | Med | phylogeny | |
| LN827582.1 | Palser et al., J Virol, 2015 | Africa | Kenya | sLCL-BL | | | | | single | Type1 | B95_8 | phylogeny | |
| LN827585.1 | Palser et al., J Virol, 2015 | Africa | Kenya | sLCL | | | | | single | Type1 | Med | phylogeny | |
| NA19315 | Santpere et al., Genome Biol Evol, 2014 | Africa | Kenya | LCL | | | | | single | Type1 | na | phylogeny | |
| KC207814.1 | Lin et al., J Virol, 2013 | Africa | Kenya | Burkitt's lymphoma | cell line | | | | single | Type1 | Med | phylogeny | Mutu |
| LN827573.1 | Palser et al., J Virol, 2015 | Africa | Kenya | sLCL | | | | | single | Type1 | na | phylogeny | |
| LN827550.1 | Palser et al., J Virol, 2015 | Africa | Kenya | sLCL | | | | | single | Type1 | Med | phylogeny | |
| LN827566.1 | Palser et al., J Virol, 2015 | Africa | Kenya | sLCL | | | | | single | Type1 | Med | phylogeny | |
| LN827552.1 | Palser et al., J Virol, 2015 | Africa | Kenya | sLCL | | | | | single | Type1 | Med | phylogeny | |
| LN827565.1 | Palser et al., J Virol, 2015 | Africa | Kenya | sLCL | | | | | single | Type1 | Med | phylogeny | |
| LN827571.1 | Palser et al., J Virol, 2015 | Africa | Kenya | sLCL-BL | | | | | single | Type1 | Med | phylogeny | |
| LN824142.1 | Palser et al., J Virol, 2015 | Western | UK | Healthy | saliva | | | | single | Type1 | China1 | phylogeny | |
| LN824226.1 | Palser et al., J Virol, 2015 | Western | UK | Hodgkin's lymphoma | | | | | single | Type1 | China1 | phylogeny | |
| LN827799.1 | Palser et al., J Virol, 2015 | Western | Australia | sLCL-IM | | | | | single | Type1 | na | phylogeny | |
| LN827586.1 | Palser et al., J Virol, 2015 | Western | Australia | sLCL-PTLD | | | | | single | Type1 | na | phylogeny | |

**Supplementary Table 1 List and summary of 270 EBV isolates sequenced in current study and 97 publicly accessed genomes included in the analysis**

**(a) Discription of 270 EBV isolates sequenced in current study and 97 publicly accessed genomes**

| Sample ID | Sequenced by | Geographic origin | Detailed geographic origin | Phenotype | Sample type | Recruitment | Sex | Age | Strain | EBV Type | LMP1 classification | Analysis | Notes |
|---|---|---|---|---|---|---|---|---|---|---|---|---|---|
| LN827589.1 | Palser et al., J Virol, 2015 | Western | Australia | sLCL-PTLD | | | | | single | Type2 | China1 | phylogeny | |
| LN827564.1 | Palser et al., J Virol, 2015 | Western | UK | Hodgkin's lymphoma | | | | | single | Type1 | na | phylogeny | |
| LN827578.1 | Palser et al., J Virol, 2015 | Western | Australia | sLCL-PTLD | | | | | single | Type1 | na | phylogeny | |
| LN827522.1 | Palser et al., J Virol, 2015 | Western | UK | Hodgkin's lymphoma | | | | | single | Type1 | China1 | phylogeny | |
| LN827596.1 | Palser et al., J Virol, 2015 | Western | Australia | sLCL-IM | | | | | single | Type1 | China1 | phylogeny | |
| LN824204.1 | Palser et al., J Virol, 2015 | Western | UK | Hodgkin's lymphoma | | | | | single | Type1 | China1 | phylogeny | |
| LN827590.1 | Palser et al., J Virol, 2015 | Western | Australia | sLCL-IM | | | | | single | Type1 | China1 | phylogeny | |
| KC440851.1 | Lei et al., BMC Genomics, 2013 | Western | USA | sLCL | | | | | single | Type1 | Med | phylogeny | |
| LN827594.1 | Palser et al., J Virol, 2015 | Western | Australia | sLCL-PTLD | | | | | single | Type1 | na | phylogeny | |
| LN824206.1 | Palser et al., J Virol, 2015 | Western | USA | sLCL-PTLD | | | | | single | Type1 | na | phylogeny | |
| LN824207.1 | Palser et al., J Virol, 2015 | Western | USA | sLCL-PTLD | | | | | single | Type1 | na | phylogeny | |
| LN827558.1 | Palser et al., J Virol, 2015 | Western | Kenya | sLCL | | | | | single | Type1 | Med | phylogeny | |
| LN827567.1 | Palser et al., J Virol, 2015 | Western | Australia | sLCL-IM | | | | | single | Type1 | B95_8 | phylogeny | |
| LN827576.1 | Palser et al., J Virol, 2015 | Western | Australia | sLCL-PTLD | | | | | single | Type1 | China1 | phylogeny | |
| KC440852.1 | Lei et al., BMC Genomics, 2013 | Western | USA | sLCL | | | | | single | Type1 | B95_8 | phylogeny | |
| LN827588.1 | Palser et al., J Virol, 2015 | Western | Australia | sLCL-PTLD | | | | | single | Type1 | B95_8 | phylogeny | |
| LN827593.1 | Palser et al., J Virol, 2015 | Western | Australia | sLCL-PTLD | | | | | single | Type1 | B95_8 | phylogeny | |
| LN827575.1 | Palser et al., J Virol, 2015 | Western | Australia | sLCL-PTLD | | | | | single | Type1 | B95_8 | phylogeny | |
| LN827583.1 | Palser et al., J Virol, 2015 | Western | Australia | sLCL-IM | | | | | single | Type1 | B95_8 | phylogeny | |
| LN827555.1 | Palser et al., J Virol, 2015 | Western | USA | LCL | | | | | single | Type1 | B95_8 | phylogeny | |
| LN827739.1 | Palser et al., J Virol, 2015 | Western | USA | LCL | | | | | single | Type1 | B95_8 | phylogeny | B95-8 (del EBER2) |
| LN827597.1 | Palser et al., J Virol, 2015 | Western | Australia | sLCL-PTLD | | | | | single | Type1 | B95_8 | phylogeny | |
| LN827572.1 | Palser et al., J Virol, 2015 | Western | Australia | sLCL-PTLD | | | | | single | Type1 | B95_8 | phylogeny | |
| LN827559.1 | Palser et al., J Virol, 2015 | Western | USA | sLCL-PTLD | | | | | single | Type1 | na | phylogeny | |

**Supplementary Table 1 List and summary of 270 EBV isolates sequenced in current study and 97 publicly accessed genomes included in the analysis**

**(a) Discription of 270 EBV isolates sequenced in current study and 97 publicly accessed genomes**

| Sample ID | Sequenced by | Geographic origin | Detailed geographic origin | Phenotype | Sample type | Recruitment | Sex | Age | Strain | EBV Type | LMP1 classification | Analysis | Notes |
|---|---|---|---|---|---|---|---|---|---|---|---|---|---|
| LN827584.1 | Palser et al., J Virol, 2015 | Western | Australia | sLCL-PTLD | | | | | single | Type1 | Med | phylogeny | |
| LN824225.1 | Palser et al., J Virol, 2015 | Western | UK | Hodgkin's lymphoma | | | | | single | Type1 | China1 | phylogeny | |
| LN827570.1 | Palser et al., J Virol, 2015 | Western | Australia | sLCL-PTLD | | | | | single | Type1 | China1 | phylogeny | |
| LN827546.1 | Palser et al., J Virol, 2015 | Western | UK | Hodgkin's lymphoma | | | | | single | Type1 | China1 | phylogeny | |
| LN827523.1 | Palser et al., J Virol, 2015 | Western | Germany | Hodgkin's lymphoma | | | | | single | Type1 | na | phylogeny | |
| LN827524.1 | Palser et al., J Virol, 2015 | Western | UK | Hodgkin's lymphoma | | | | | single | Type1 | Med | phylogeny | |
| LN827592.1 | Palser et al., J Virol, 2015 | Western | Australia | sLCL-PTLD | | | | | single | Type1 | Med | phylogeny | |
| LN827553.1 | Palser et al., J Virol, 2015 | Western | Australia | sLCL-PTLD | | | | | single | Type1 | na | phylogeny | |
| LN827569.1 | Palser et al., J Virol, 2015 | Western | Australia | sLCL-PTLD | | | | | single | Type1 | na | phylogeny | |
| LN827595.1 | Palser et al., J Virol, 2015 | Western | Australia | sLCL-PTLD | | | | | single | Type1 | Med | phylogeny | |
| KC207813.1 | Lin et al., J Virol, 2013 | East Asia | Japan | Burkitt's lymphoma | cell line | | | | single | Type1 | China1 | phylogeny | |
| LN824208.1 | Palser et al., J Virol, 2015 | East Asia | Japan | Burkitt's lymphoma | cell line | | | | single | Type1 | China1 | phylogeny | Akata |
| LN827561.1 | Palser et al., J Virol, 2015 | East Asia | South Korea | Gastric carcinoma | cell line | | | | single | Type1 | China1 | phylogeny | |
| LN827525.1 | Palser et al., J Virol, 2015 | East Asia | NPC-endemic China | NPC | cell line | | | | single | Type1 | China1 | phylogeny | C666-1 resequencing |
| LN824209.1 | Palser et al., J Virol, 2015 | East Asia | NPC-endemic China | sLCL | | | | | single | Type1 | China1 | phylogeny | |
| LN824224.1 | Palser et al., J Virol, 2015 | East Asia | NPC-endemic China | sLCL | | | | | single | Type1 | China1 | phylogeny | |
| LN827547.1 | Palser et al., J Virol, 2015 | East Asia | NPC-endemic China | sLCL | | | | | single | Type1 | China1 | phylogeny | |
| AY961628.3 | Zeng et al., J Virol, 2005 | East Asia | NPC-endemic China | NPC | saliva | | | | single | Type1 | China1 | phylogeny | |
| LN827549.1 | Palser et al., J Virol, 2015 | East Asia | NPC-endemic China | NPC | tumor biopsy | | | | single | Type1 | China1 | phylogeny | |
| KF992568.1 | Kwok et al., J Virol, 2014 | East Asia | NPC-endemic China | NPC | tumor biopsy | | | | single | Type1 | China1 | phylogeny | |
| KF992569.1 | Kwok et al., J Virol, 2014 | East Asia | NPC-endemic China | NPC | tumor biopsy | | | | single | Type1 | China1 | phylogeny | |
| KF992564.1 | Kwok et al., J Virol, 2014 | East Asia | NPC-endemic China | NPC | tumor biopsy | | | | single | Type1 | China1 | phylogeny | |
| JQ009376.2 | Kwok et al., PLoS One, 2012 | East Asia | NPC-endemic China | NPC | tumor biopsy | | | | single | Type1 | China1 | phylogeny | |
| KF992566.1 | Kwok et al., J Virol, 2014 | East Asia | NPC-endemic China | NPC | tumor biopsy | | | | single | Type1 | China1 | phylogeny | |

**Supplementary Table 1 List and summary of 270 EBV isolates sequenced in current study and 97 publicly accessed genomes included in the analysis**

**(a) Discription of 270 EBV isolates sequenced in current study and 97 publicly accessed genomes**

| Sample ID | Sequenced by | Geographic origin | Detailed geographic origin | Phenotype | Sample type | Recruitment | Sex | Age | Strain | EBV Type | LMP1 classification | Analysis | Notes |
|---|---|---|---|---|---|---|---|---|---|---|---|---|---|
| HQ020558.1 | Liu et al., J Virol, 2011 | East Asia | NPC-endemic China | NPC | tumor biopsy | | | | single | Type1 | China1 | phylogeny | |
| KF373730.1 | Tsai et al., Cell Rep, 2013 | East Asia | NPC-endemic China | NPC | cell line | | | | single | Type1 | China1 | phylogeny | M81 |
| KF992565.1 | Kwok et al., J Virol, 2014 | East Asia | NPC-endemic China | NPC | tumor biopsy | | | | single | Type1 | China1 | phylogeny | |
| KF992571.1 | Kwok et al., J Virol, 2014 | East Asia | NPC-endemic China | NPC | tumor biopsy | | | | single | Type1 | China1 | phylogeny | |
| KF992567.1 | Kwok et al., J Virol, 2014 | East Asia | NPC-endemic China | NPC | tumor biopsy | | | | single | Type1 | China1 | phylogeny | |
| KF992570.1 | Kwok et al., J Virol, 2014 | East Asia | NPC-endemic China | NPC | tumor biopsy | | | | single | Type1 | China1 | phylogeny | |
| KC617875.1 | Tso et al., Infect Agent Cancer, 2013 | East Asia | NPC-endemic China | NPC | cell line | | | | single | Type1 | China1 | phylogeny | C666-1 |

NPC, Nasopharyngeal carcinoma; LCL, lymphoblastoid cell line; sLCL, spontaneous lymphoblastoid cell line; PTLD, posttransplant lymphoproliferative disease; IM, infectious mononucleosis.
SYSUCC, the Sun Yat-sen University Cancer Center; FAHGMC, the First Affiliated Hospital of Guangxi Medical College; AHQU, the Affiliated Hospital of the Qingdao University.
Geographic origin indicates the birth places of participants in our study or geographic records of samples in published papers. NPC-endemic China includes Guangdong and Guangxi Provinces in China; NPC-non-endemic China includes other provinces in China.
LMP-1 classification is based on LMP-1 C-terminal signature reported in Edwards et al, Virology, 1999, and na indicates strains cannot be determined by LMP-1 C-terminal signature.

**Supplementary Table 1 List and summary of 270 EBV isolates sequenced in current study and 97 publicly accessed genomes included in the analysis**

**(b) Summary of geographic origins and phenotypes of 367 EBV isolates**

| | Africa | Western countries | NPC-endemic China | NPC-non-endemic East Asia | Total |
|---|---|---|---|---|---|
| **NPC** | | | **175** | **20** | **195** |
| **Gastric carcinoma** | | | | **17** | **17** |
| **Healthy control** | | **1** | **47** | **7** | **55** |
| **Lymphoma** | | | | | **43** |
| Hodgkin | | 8 | 8 | 3 | |
| Burkkit | 13 | | 2 | 2 | |
| NK/T cell | | | 5 | 2 | |
| **LCL** | **24** | **5** | | **3** | **32** |
| **sLCL-IM** | | 5 | | | **5** |
| **sLCL-PTLD** | | **19** | | | **19** |
| **Total** | **37** | **38** | **237** | **54** | **366** |

One single published strain was sequenced from Burkkit's lymphoma cell line with Papua New Guinea origin.

**Supplementary Table 1 List and summary of 270 EBV isolates sequenced in current study and 97 publicly accessed genomes included in the analysis**

**(c) Summary of age and sex of participants from whom EBV were sequenced in current study.**

| | Male | | | Male total | Female | | | Female total |
|---|---|---|---|---|---|---|---|---|
| | Age | | | | Age | | | |
| | 0-27 | 28-59 | 60- | | 0-27 | 28-59 | 60- | |
| **Hodgkin lymphoma** | | | | **11** | | | | |
| NPC-endemic China | | 6 | 2 | | | | | |
| non-endemic China | 1 | | 2 | | | | | |
| **Burkitt lymphoma** | | | | **2** | | | | |
| NPC-endemic China | | 2 | | | | | | |
| **NPC** | | | | **130** | | | | **50** |
| NPC-endemic China | 1 | 95 | 21 | | 3 | 30 | 10 | |
| non-endemic China | 1 | 10 | 2 | | | 5 | 2 | |
| **Gastric carcinoma** | | | | **14** | | | | **2** |
| non-endemic China | 1 | 9 | 4 | | | 2 | | |
| **Healthy control** | | | | **41** | | | | **13** |
| NPC-endemic China | | 31 | 7 | | 1 | 5 | 3 | |
| non-endemic China | 2 | 1 | | | 1 | 3 | | |
| **NK/T cell lymphoma** | | | | **6** | | | | **1** |
| NPC-endemic China | 1 | 4 | | | | | | |
| non-endemic China | 1 | | | | | 1 | | |
| **Total** | | | | **204** | | | | **66** |

**Supplementary Table 1 List and summary of 270 EBV isolates sequenced in current study and 97 publicly accessed genomes included in the analysis**

**(d) Summary of geographic origins, phenotypes and sample types of 270 EBV isolates sequenced in current study.**

| | **NPC-endemic China** | **NPC-non-endemic China** |
|---|---|---|
| **NPC** | **160** | **20** |
| Tumor tissue | 105 | 19 |
| Saliva | 54 | |
| Plasma | | 1 |
| Cell line | 1 | |
| **Healthy control saliva** | **47** | **7** |
| **Lymphoma biopsy** | **15** | **5** |
| Hodgkin | 8 | 3 |
| Burkkit | 2 | |
| NK/T cell | 5 | 2 |
| **Gastric carcinoma tissue** | | **16** |
| **Total** | 222 | **48** |

**Supplementary Table 2 Variant information of EBV genome isolates sequenced in current study.**

| Sample ID | nVariants | VarFreq | nSNPs | nInsertions | nDeletions | nComplex | nIndel |
|---|---|---|---|---|---|---|---|
| BLT001 | 1151 | 0.67% | 1043 | 45 | 45 | 18 | 108 |
| BLT002 | 1186 | 0.69% | 1059 | 49 | 56 | 22 | 127 |
| C666 | 1209 | 0.70% | 1072 | 42 | 67 | 28 | 137 |
| GCT001 | 1116 | 0.65% | 1011 | 44 | 45 | 16 | 105 |
| GCT002 | 1114 | 0.65% | 1007 | 42 | 42 | 23 | 107 |
| GCT003 | 1081 | 0.63% | 969 | 42 | 49 | 21 | 112 |
| GCT004 | 1006 | 0.59% | 909 | 35 | 46 | 16 | 97 |
| GCT005 | 1114 | 0.65% | 989 | 39 | 64 | 22 | 125 |
| GCT006 | 1123 | 0.65% | 998 | 48 | 57 | 20 | 125 |
| GCT007 | 1129 | 0.66% | 1018 | 40 | 48 | 23 | 111 |
| GCT008 | 1131 | 0.66% | 1007 | 46 | 56 | 22 | 124 |
| GCT009 | 1091 | 0.63% | 986 | 42 | 48 | 15 | 105 |
| GCT010 | 1164 | 0.68% | 1055 | 42 | 46 | 21 | 109 |
| GCT011 | 1116 | 0.65% | 1021 | 39 | 36 | 20 | 95 |
| GCT012 | 1080 | 0.63% | 980 | 37 | 42 | 21 | 100 |
| GCT013 | 1122 | 0.65% | 1002 | 44 | 57 | 19 | 120 |
| GCT014 | 1138 | 0.66% | 1027 | 47 | 45 | 19 | 111 |
| GCT015 | 1148 | 0.67% | 1033 | 44 | 50 | 21 | 115 |
| GCT016 | 1076 | 0.63% | 977 | 33 | 46 | 20 | 99 |
| HLT001 | 1163 | 0.68% | 1058 | 40 | 46 | 19 | 105 |
| HLT002 | 1180 | 0.69% | 1067 | 37 | 59 | 17 | 113 |
| HLT003 | 1829 | 1.06% | 1717 | 43 | 55 | 14 | 112 |
| HLT004 | 1734 | 1.01% | 1632 | 40 | 49 | 13 | 102 |
| HLT005 | 1143 | 0.67% | 1044 | 35 | 45 | 19 | 99 |
| HLT006 | 1155 | 0.67% | 1056 | 37 | 41 | 21 | 99 |
| HLT007 | 1239 | 0.72% | 1154 | 38 | 30 | 17 | 85 |
| HLT009 | 1565 | 0.91% | 1473 | 35 | 45 | 12 | 92 |
| HLT010 | 1114 | 0.65% | 1029 | 33 | 33 | 19 | 85 |
| HLT011 | 1257 | 0.73% | 1174 | 31 | 35 | 17 | 83 |
| HLT012 | 1569 | 0.91% | 1473 | 36 | 40 | 20 | 96 |
| HS001 | 1035 | 0.60% | 944 | 38 | 37 | 16 | 91 |
| HS003 | 1125 | 0.65% | 1020 | 39 | 45 | 21 | 105 |
| HS005 | 1255 | 0.73% | 1148 | 40 | 51 | 16 | 107 |
| HS006 | 1057 | 0.62% | 956 | 38 | 44 | 19 | 101 |
| HS007 | 1187 | 0.69% | 1059 | 49 | 64 | 15 | 128 |
| HS008 | 1151 | 0.67% | 1037 | 47 | 52 | 15 | 114 |
| HS009 | 1047 | 0.61% | 959 | 38 | 37 | 13 | 88 |
| HS010 | 1725 | 1.00% | 1609 | 48 | 55 | 13 | 116 |
| HS011 | 1217 | 0.71% | 1101 | 44 | 54 | 18 | 116 |
| HS012 | 1206 | 0.70% | 1097 | 49 | 49 | 11 | 109 |

**Supplementary Table 2 Variant information of EBV genome isolates sequenced in current study.**

| Sample ID | nVariants | VarFreq | nSNPs | nInsertions | nDeletions | nComplex | nIndel |
|---|---|---|---|---|---|---|---|
| HS013 | 1170 | 0.68% | 1049 | 45 | 53 | 23 | 121 |
| HS014 | 1156 | 0.67% | 1048 | 38 | 48 | 22 | 108 |
| HS015 | 1197 | 0.70% | 1093 | 38 | 49 | 17 | 104 |
| HS016 | 1180 | 0.69% | 1052 | 48 | 55 | 25 | 128 |
| HS017 | 1949 | 1.13% | 1832 | 43 | 58 | 16 | 117 |
| HS018 | 1214 | 0.71% | 1104 | 45 | 49 | 16 | 110 |
| HS019 | 1140 | 0.66% | 1026 | 45 | 52 | 17 | 114 |
| HS020 | 1141 | 0.66% | 1039 | 39 | 47 | 16 | 102 |
| HS021 | 1129 | 0.66% | 1028 | 37 | 50 | 14 | 101 |
| HS022 | 2104 | 1.22% | 1972 | 45 | 61 | 26 | 132 |
| HS023 | 1188 | 0.69% | 1086 | 41 | 40 | 21 | 102 |
| HS024 | 1123 | 0.65% | 1037 | 34 | 31 | 21 | 86 |
| HS025 | 1257 | 0.73% | 1123 | 46 | 69 | 19 | 134 |
| HS026 | 1196 | 0.70% | 1117 | 35 | 27 | 17 | 79 |
| HS027 | 1153 | 0.67% | 1036 | 45 | 49 | 23 | 117 |
| HS028 | 1621 | 0.94% | 1523 | 46 | 36 | 16 | 98 |
| HS029 | 1142 | 0.66% | 1055 | 30 | 37 | 20 | 87 |
| HS030 | 1601 | 0.93% | 1516 | 36 | 40 | 9 | 85 |
| HS031 | 1220 | 0.71% | 1110 | 40 | 48 | 22 | 110 |
| HS032 | 1156 | 0.67% | 1057 | 39 | 38 | 22 | 99 |
| HS033 | 1147 | 0.67% | 1054 | 32 | 40 | 21 | 93 |
| HS034 | 1151 | 0.67% | 1040 | 44 | 50 | 17 | 111 |
| HS035 | 1303 | 0.76% | 1194 | 45 | 45 | 19 | 109 |
| HS036 | 1194 | 0.69% | 1077 | 49 | 49 | 19 | 117 |
| HS037 | 1094 | 0.64% | 983 | 40 | 52 | 19 | 111 |
| HS038 | 1179 | 0.69% | 1052 | 43 | 60 | 24 | 127 |
| HS039 | 1260 | 0.73% | 1136 | 51 | 51 | 22 | 124 |
| HS040 | 1508 | 0.88% | 1371 | 48 | 72 | 17 | 137 |
| HS041 | 1172 | 0.68% | 1050 | 45 | 56 | 21 | 122 |
| HS045 | 1177 | 0.69% | 1048 | 48 | 58 | 23 | 129 |
| HS046 | 1819 | 1.06% | 1672 | 52 | 73 | 22 | 147 |
| HS048 | 1349 | 0.79% | 1209 | 46 | 70 | 24 | 140 |
| HS050 | 1211 | 0.70% | 1077 | 47 | 62 | 25 | 134 |
| HS051 | 1178 | 0.69% | 1050 | 47 | 61 | 20 | 128 |
| HS052 | 1201 | 0.70% | 1065 | 49 | 64 | 23 | 136 |
| HS053 | 1228 | 0.71% | 1113 | 42 | 57 | 16 | 115 |
| HS054 | 1101 | 0.64% | 1013 | 35 | 38 | 15 | 88 |
| HS055 | 1485 | 0.86% | 1355 | 52 | 55 | 23 | 130 |
| HS056 | 1815 | 1.06% | 1692 | 46 | 59 | 18 | 123 |
| HS057 | 1167 | 0.68% | 1059 | 39 | 47 | 22 | 108 |

**Supplementary Table 2 Variant information of EBV genome isolates sequenced in current study.**

| Sample ID | nVariants | VarFreq | nSNPs | nInsertions | nDeletions | nComplex | nIndel |
|---|---|---|---|---|---|---|---|
| NHS001 | 1723 | 1.00% | 1626 | 39 | 43 | 15 | 97 |
| NHS002 | 1184 | 0.69% | 1072 | 45 | 47 | 20 | 112 |
| NHS003 | 1771 | 1.03% | 1655 | 46 | 54 | 16 | 116 |
| NHS004 | 1178 | 0.69% | 1085 | 37 | 35 | 21 | 93 |
| NKLT001 | 1444 | 0.84% | 1335 | 40 | 46 | 23 | 109 |
| NKLT002 | 1038 | 0.60% | 953 | 30 | 36 | 19 | 85 |
| NKLT003-2 | 1137 | 0.66% | 1064 | 31 | 31 | 11 | 73 |
| NKLT004 | 1048 | 0.61% | 955 | 33 | 38 | 22 | 93 |
| NKLT005 | 1423 | 0.83% | 1318 | 41 | 42 | 22 | 105 |
| NKLT006 | 1137 | 0.66% | 1044 | 34 | 42 | 17 | 93 |
| NKLT007 | 1146 | 0.67% | 1039 | 40 | 46 | 21 | 107 |
| NNPCT001 | 1214 | 0.71% | 1108 | 43 | 43 | 20 | 106 |
| NNPCT002 | 1062 | 0.62% | 965 | 36 | 38 | 23 | 97 |
| NNPCT003 | 1234 | 0.72% | 1114 | 40 | 59 | 21 | 120 |
| NNPCT004 | 1257 | 0.73% | 1149 | 43 | 49 | 16 | 108 |
| NNPCT005 | 1093 | 0.64% | 986 | 43 | 47 | 17 | 107 |
| NNPCT006 | 1571 | 0.91% | 1455 | 48 | 54 | 14 | 116 |
| NPCP001 | 1133 | 0.66% | 1021 | 41 | 49 | 22 | 112 |
| NPCS001 | 1115 | 0.65% | 1028 | 35 | 42 | 10 | 87 |
| NPCS002 | 1127 | 0.66% | 1025 | 37 | 48 | 17 | 102 |
| NPCS003-2 | 1148 | 0.67% | 1043 | 37 | 52 | 16 | 105 |
| NPCS004 | 1361 | 0.79% | 1272 | 34 | 38 | 17 | 89 |
| NPCS005 | 1156 | 0.67% | 1035 | 35 | 67 | 19 | 121 |
| NPCS006 | 1170 | 0.68% | 1054 | 40 | 50 | 26 | 116 |
| NPCS007 | 1065 | 0.62% | 999 | 24 | 31 | 11 | 66 |
| NPCS008 | 1075 | 0.63% | 995 | 28 | 38 | 14 | 80 |
| NPCS009 | 1109 | 0.65% | 1007 | 36 | 46 | 20 | 102 |
| NPCS010 | 1233 | 0.72% | 1130 | 38 | 47 | 18 | 103 |
| NPCS011 | 1130 | 0.66% | 1033 | 37 | 45 | 15 | 97 |
| NPCS012 | 1147 | 0.67% | 1040 | 38 | 51 | 18 | 107 |
| NPCS013 | 1451 | 0.84% | 1309 | 49 | 74 | 19 | 142 |
| NPCS014 | 1124 | 0.65% | 1021 | 39 | 48 | 16 | 103 |
| NPCS015 | 1239 | 0.72% | 1100 | 50 | 64 | 25 | 139 |
| NPCS016 | 1093 | 0.64% | 1008 | 33 | 37 | 15 | 85 |
| NPCS017 | 1119 | 0.65% | 1028 | 35 | 38 | 18 | 91 |
| NPCS018 | 1143 | 0.67% | 1033 | 33 | 61 | 16 | 110 |
| NPCS019 | 1127 | 0.66% | 1024 | 33 | 48 | 22 | 103 |
| NPCS021 | 1131 | 0.66% | 1028 | 34 | 46 | 23 | 103 |
| NPCS022 | 1120 | 0.65% | 1029 | 35 | 42 | 14 | 91 |
| NPCS023 | 1223 | 0.71% | 1117 | 43 | 44 | 19 | 106 |

**Supplementary Table 2 Variant information of EBV genome isolates sequenced in current study.**

| Sample ID | nVariants | VarFreq | nSNPs | nInsertions | nDeletions | nComplex | nIndel |
|---|---|---|---|---|---|---|---|
| NPCS024 | 1147 | 0.67% | 1045 | 35 | 51 | 16 | 102 |
| NPCS025 | 1145 | 0.67% | 1035 | 36 | 52 | 22 | 110 |
| NPCS026 | 1142 | 0.66% | 1037 | 39 | 49 | 17 | 105 |
| NPCS027 | 1147 | 0.67% | 1042 | 37 | 52 | 16 | 105 |
| NPCS028 | 1128 | 0.66% | 1026 | 42 | 47 | 13 | 102 |
| NPCS029 | 1193 | 0.69% | 1077 | 42 | 53 | 21 | 116 |
| NPCS030 | 1190 | 0.69% | 1069 | 45 | 54 | 22 | 121 |
| NPCS031 | 1128 | 0.66% | 1040 | 36 | 35 | 17 | 88 |
| NPCS032 | 1265 | 0.74% | 1120 | 50 | 75 | 20 | 145 |
| NPCS033 | 1120 | 0.65% | 1022 | 34 | 45 | 19 | 98 |
| NPCS034 | 1110 | 0.65% | 1017 | 36 | 38 | 19 | 93 |
| NPCS035 | 1179 | 0.69% | 1053 | 46 | 56 | 24 | 126 |
| NPCS036 | 1453 | 0.85% | 1366 | 34 | 42 | 11 | 87 |
| NPCS037 | 1131 | 0.66% | 1060 | 28 | 27 | 16 | 71 |
| NPCS038 | 1144 | 0.67% | 1036 | 37 | 49 | 22 | 108 |
| NPCS039 | 1148 | 0.67% | 1036 | 37 | 53 | 22 | 112 |
| NPCS040 | 1148 | 0.67% | 1022 | 43 | 57 | 26 | 126 |
| NPCS041 | 1585 | 0.92% | 1474 | 43 | 55 | 13 | 111 |
| NPCS042 | 1130 | 0.66% | 1035 | 33 | 41 | 21 | 95 |
| NPCS043 | 1914 | 1.11% | 1787 | 51 | 56 | 20 | 127 |
| NPCS044 | 1255 | 0.73% | 1140 | 46 | 50 | 19 | 115 |
| NPCS045 | 1117 | 0.65% | 1015 | 38 | 48 | 16 | 102 |
| NPCS046 | 1186 | 0.69% | 1063 | 45 | 64 | 14 | 123 |
| NPCS047 | 1110 | 0.65% | 1018 | 34 | 42 | 16 | 92 |
| NPCS048 | 1173 | 0.68% | 1055 | 41 | 56 | 21 | 118 |
| NPCS049 | 1302 | 0.76% | 1189 | 42 | 51 | 20 | 113 |
| NPCS050 | 1169 | 0.68% | 1042 | 37 | 67 | 23 | 127 |
| NPCS051 | 1139 | 0.66% | 1038 | 35 | 47 | 19 | 101 |
| NPCS052 | 1232 | 0.72% | 1105 | 43 | 62 | 22 | 127 |
| NPCS054 | 1150 | 0.67% | 1036 | 45 | 49 | 20 | 114 |
| NPCS055 | 1789 | 1.04% | 1667 | 44 | 53 | 25 | 122 |
| NPCT001 | 1143 | 0.67% | 1022 | 43 | 54 | 24 | 121 |
| NPCT002 | 1159 | 0.67% | 1032 | 44 | 58 | 25 | 127 |
| NPCT003 | 1173 | 0.68% | 1043 | 49 | 58 | 23 | 130 |
| NPCT004 | 1263 | 0.74% | 1154 | 44 | 44 | 21 | 109 |
| NPCT005 | 1262 | 0.73% | 1143 | 48 | 50 | 21 | 119 |
| NPCT006 | 1175 | 0.68% | 1055 | 45 | 50 | 25 | 120 |
| NPCT007 | 1152 | 0.67% | 1033 | 52 | 51 | 16 | 119 |
| NPCT008 | 1178 | 0.69% | 1062 | 42 | 54 | 20 | 116 |
| NPCT009 | 1185 | 0.69% | 1075 | 44 | 45 | 21 | 110 |

**Supplementary Table 2 Variant information of EBV genome isolates sequenced in current study.**

| Sample ID | nVariants | VarFreq | nSNPs | nInsertions | nDeletions | nComplex | nIndel |
|---|---|---|---|---|---|---|---|
| NPCT010 | 1170 | 0.68% | 1048 | 43 | 60 | 19 | 122 |
| NPCT011 | 1241 | 0.72% | 1127 | 49 | 46 | 19 | 114 |
| NPCT012 | 1179 | 0.69% | 1052 | 46 | 57 | 24 | 127 |
| NPCT013 | 1153 | 0.67% | 1027 | 45 | 61 | 20 | 126 |
| NPCT014 | 1172 | 0.68% | 1053 | 42 | 55 | 22 | 119 |
| NPCT015 | 1246 | 0.73% | 1133 | 41 | 51 | 21 | 113 |
| NPCT016 | 1872 | 1.09% | 1744 | 50 | 58 | 20 | 128 |
| NPCT017 | 1186 | 0.69% | 1057 | 44 | 64 | 21 | 129 |
| NPCT018 | 1166 | 0.68% | 1042 | 44 | 56 | 24 | 124 |
| NPCT019 | 1200 | 0.70% | 1077 | 47 | 58 | 18 | 123 |
| NPCT020-2 | 1229 | 0.72% | 1124 | 38 | 53 | 14 | 105 |
| NPCT021 | 1922 | 1.12% | 1788 | 50 | 62 | 22 | 134 |
| NPCT022 | 1188 | 0.69% | 1062 | 44 | 61 | 21 | 126 |
| NPCT023 | 1168 | 0.68% | 1052 | 36 | 55 | 25 | 116 |
| NPCT024 | 1197 | 0.70% | 1053 | 46 | 80 | 18 | 144 |
| NPCT025 | 1302 | 0.76% | 1179 | 46 | 57 | 20 | 123 |
| NPCT026 | 1609 | 0.94% | 1490 | 48 | 55 | 16 | 119 |
| NPCT027 | 1168 | 0.68% | 1042 | 45 | 56 | 25 | 126 |
| NPCT028-2 | 1167 | 0.68% | 1052 | 43 | 55 | 17 | 115 |
| NPCT029 | 1148 | 0.67% | 1038 | 41 | 49 | 20 | 110 |
| NPCT031 | 1204 | 0.70% | 1074 | 42 | 70 | 18 | 130 |
| NPCT032 | 1179 | 0.69% | 1058 | 42 | 61 | 18 | 121 |
| NPCT033 | 1195 | 0.70% | 1075 | 43 | 55 | 22 | 120 |
| NPCT035 | 1161 | 0.68% | 1049 | 41 | 51 | 20 | 112 |
| NPCT036 | 1165 | 0.68% | 1050 | 43 | 55 | 17 | 115 |
| NPCT037 | 1153 | 0.67% | 1041 | 46 | 49 | 17 | 112 |
| NPCT038 | 1137 | 0.66% | 1046 | 36 | 36 | 19 | 91 |
| NPCT039 | 1237 | 0.72% | 1142 | 36 | 43 | 16 | 95 |
| NPCT040 | 1187 | 0.69% | 1065 | 42 | 57 | 23 | 122 |
| NPCT041 | 1169 | 0.68% | 1047 | 49 | 57 | 16 | 122 |
| NPCT042 | 1272 | 0.74% | 1153 | 46 | 51 | 22 | 119 |
| NPCT043 | 1193 | 0.69% | 1062 | 45 | 58 | 28 | 131 |
| NPCT045 | 1175 | 0.68% | 1054 | 42 | 59 | 20 | 121 |
| NPCT046 | 1456 | 0.85% | 1322 | 45 | 68 | 21 | 134 |
| NPCT047 | 1255 | 0.73% | 1131 | 46 | 55 | 23 | 124 |
| NPCT048 | 1195 | 0.70% | 1071 | 41 | 61 | 22 | 124 |
| NPCT049 | 1810 | 1.05% | 1690 | 46 | 55 | 19 | 120 |
| NPCT050 | 1190 | 0.69% | 1054 | 44 | 71 | 21 | 136 |
| NPCT051 | 1505 | 0.88% | 1392 | 45 | 50 | 18 | 113 |
| NPCT052 | 1161 | 0.68% | 1037 | 44 | 56 | 24 | 124 |

**Supplementary Table 2 Variant information of EBV genome isolates sequenced in current study.**

| Sample ID | nVariants | VarFreq | nSNPs | nInsertions | nDeletions | nComplex | nIndel |
|---|---|---|---|---|---|---|---|
| NPCT053 | 1175 | 0.68% | 1041 | 49 | 63 | 22 | 134 |
| NPCT054 | 1184 | 0.69% | 1054 | 45 | 61 | 24 | 130 |
| NPCT054M | 1194 | 0.69% | 1063 | 48 | 59 | 24 | 131 |
| NPCT055 | 1118 | 0.65% | 1026 | 33 | 43 | 16 | 92 |
| NPCT055M | 1151 | 0.67% | 1038 | 38 | 54 | 21 | 113 |
| NPCT056 | 1128 | 0.66% | 1027 | 36 | 44 | 21 | 101 |
| NPCT056M | 1116 | 0.65% | 1022 | 33 | 41 | 20 | 94 |
| NPCT057 | 1059 | 0.62% | 965 | 35 | 41 | 18 | 94 |
| NPCT057M | 1078 | 0.63% | 970 | 41 | 51 | 16 | 108 |
| NPCT058 | 1160 | 0.68% | 1043 | 41 | 54 | 22 | 117 |
| NPCT058M | 1171 | 0.68% | 1049 | 44 | 55 | 23 | 122 |
| NPCT059 | 1174 | 0.68% | 1058 | 44 | 49 | 23 | 116 |
| NPCT060 | 1219 | 0.71% | 1098 | 43 | 55 | 23 | 121 |
| NPCT061 | 1140 | 0.66% | 1017 | 44 | 55 | 24 | 123 |
| NPCT062 | 1191 | 0.69% | 1094 | 39 | 39 | 19 | 97 |
| NPCT063 | 1166 | 0.68% | 1048 | 42 | 51 | 25 | 118 |
| NPCT064 | 1188 | 0.69% | 1060 | 46 | 58 | 24 | 128 |
| NPCT065 | 1227 | 0.71% | 1099 | 49 | 53 | 26 | 128 |
| NPCT066 | 1134 | 0.66% | 1032 | 36 | 45 | 21 | 102 |
| NPCT067 | 1152 | 0.67% | 1042 | 41 | 50 | 19 | 110 |
| NPCT068 | 1202 | 0.70% | 1077 | 41 | 65 | 19 | 125 |
| NPCT069 | 1158 | 0.67% | 1045 | 45 | 48 | 20 | 113 |
| NPCT070 | 1157 | 0.67% | 1036 | 41 | 62 | 18 | 121 |
| NPCT071 | 1180 | 0.69% | 1058 | 43 | 55 | 24 | 122 |
| NPCT072 | 1229 | 0.72% | 1095 | 46 | 65 | 23 | 134 |
| NPCT073 | 1186 | 0.69% | 1058 | 46 | 58 | 24 | 128 |
| NPCT074 | 1184 | 0.69% | 1057 | 40 | 69 | 18 | 127 |
| NPCT074S | 1149 | 0.67% | 1034 | 36 | 59 | 20 | 115 |
| NPCT075 | 1221 | 0.71% | 1109 | 40 | 50 | 22 | 112 |
| NPCT076 | 1155 | 0.67% | 1041 | 41 | 53 | 20 | 114 |
| NPCT077 | 1174 | 0.68% | 1056 | 43 | 53 | 22 | 118 |
| NPCT078 | 1162 | 0.68% | 1043 | 43 | 53 | 23 | 119 |
| NPCT079 | 1541 | 0.90% | 1420 | 44 | 54 | 23 | 121 |
| NPCT080 | 1170 | 0.68% | 1050 | 44 | 58 | 18 | 120 |
| NPCT081 | 1166 | 0.68% | 1051 | 40 | 50 | 25 | 115 |
| NPCT082 | 1179 | 0.69% | 1057 | 41 | 58 | 23 | 122 |
| NPCT083 | 1167 | 0.68% | 1043 | 46 | 56 | 22 | 124 |
| NPCT084 | 1202 | 0.70% | 1106 | 38 | 39 | 19 | 96 |
| NPCT085 | 1124 | 0.65% | 1025 | 41 | 43 | 15 | 99 |
| NPCT086 | 1167 | 0.68% | 1043 | 42 | 59 | 23 | 124 |

**Supplementary Table 2 Variant information of EBV genome isolates sequenced in current study.**

| Sample ID | nVariants | VarFreq | nSNPs | nInsertions | nDeletions | nComplex | nIndel |
|---|---|---|---|---|---|---|---|
| NPCT087 | 1246 | 0.73% | 1141 | 37 | 46 | 22 | 105 |
| NPCT088 | 1156 | 0.67% | 1045 | 40 | 53 | 18 | 111 |
| NPCT089 | 1204 | 0.70% | 1089 | 42 | 54 | 19 | 115 |
| NPCT090 | 1264 | 0.74% | 1154 | 41 | 50 | 19 | 110 |
| NPCT091 | 1173 | 0.68% | 1054 | 39 | 58 | 22 | 119 |
| NPCT092 | 1183 | 0.69% | 1056 | 43 | 59 | 25 | 127 |
| NPCT093 | 1165 | 0.68% | 1052 | 42 | 53 | 18 | 113 |
| NPCT094 | 1139 | 0.66% | 1038 | 34 | 45 | 22 | 101 |
| NPCT095 | 1705 | 0.99% | 1584 | 42 | 58 | 21 | 121 |
| NPCT096 | 1171 | 0.68% | 1048 | 48 | 55 | 20 | 123 |
| NPCT097 | 1834 | 1.07% | 1708 | 47 | 60 | 19 | 126 |
| NPCT098 | 1181 | 0.69% | 1053 | 42 | 62 | 24 | 128 |
| NPCT099 | 1166 | 0.68% | 1044 | 44 | 62 | 16 | 122 |
| NPCT100 | 1165 | 0.68% | 1042 | 46 | 58 | 19 | 123 |
| NPCT101 | 1133 | 0.66% | 1037 | 38 | 39 | 19 | 96 |
| NPCT102 | 1164 | 0.68% | 1043 | 38 | 60 | 23 | 121 |
| NPCT103 | 1174 | 0.68% | 1046 | 46 | 64 | 18 | 128 |
| NPCT104 | 1163 | 0.68% | 1050 | 40 | 47 | 26 | 113 |
| NPCT105 | 1103 | 0.64% | 1008 | 38 | 38 | 19 | 95 |
| NPCT106 | 1228 | 0.71% | 1120 | 40 | 46 | 22 | 108 |
| NPCT107 | 1168 | 0.68% | 1047 | 41 | 54 | 26 | 121 |
| NPCT108 | 1144 | 0.67% | 1043 | 38 | 42 | 21 | 101 |
| NPCT109 | 1315 | 0.77% | 1200 | 41 | 51 | 23 | 115 |
| NPCT110 | 1162 | 0.68% | 1047 | 41 | 54 | 20 | 115 |
| NPCT111 | 1359 | 0.79% | 1232 | 46 | 57 | 24 | 127 |
| NPCT112 | 1140 | 0.66% | 1030 | 41 | 47 | 22 | 110 |
| NPCT113 | 1164 | 0.68% | 1042 | 44 | 60 | 18 | 122 |
| NPCT114 | 1169 | 0.68% | 1051 | 47 | 50 | 21 | 118 |
| NPCT115 | 1190 | 0.69% | 1062 | 42 | 67 | 19 | 128 |
| NPCT116 | 1179 | 0.69% | 1064 | 42 | 50 | 23 | 115 |
| All | 8469 | 4.93% | 8015 | 140 | 229 | 85 | 454 |

**Supplementary Table 3 Concordance rate between SNPs from C666-1 EBV genome sequenced in current study and in published study.**

| **C666-1** | | **Current study** | | **Total** |
|---|---|---|---|---|
| | | nVariants | nReferences | |
| **Published*** | nVariants | 1021[a] | 117[b] | 1138 |
| | nReferences | 51[c] | 6943[d] | 6994 |
| **Total** | | 1072 | 7060 | 8132 |

Concordance rate was 97.93%, calculated by (a+d)/(a+b+c+d).

*GenBank accession number: KC617875.1

**Supplementary Table 4 Concordance rate between variants discovered by targeted EBV whole-genome sequencing (EBV-WGS) and Sanger sequencing.**

| | | **EBV-WGS** | | **Total** |
|---|---|---|---|---|
| | | nVariants | nReferences | |
| **Sanger** | nVariants | 153[a] | 3[b] | 156 |
| | nReferences | 5[c] | 165[d] | 170 |
| **Total** | | 158 | 168 | 326 |

Concordance rate was 97.55%, calculated by (a+d)/(a+b+c+d).

**Supplementary Table 5 Concordance rate between variants discovered by targeted EBV whole-genome sequencing (EBV-WGS) and MassArray iPlex assay.**

| | | **EBV-WGS** | | **Total** |
|---|---|---|---|---|
| | | nVariants | nReferences | |
| **MassArray iPlex assay** | nVariants | 4328[a] | 0[b] | 4328 |
| | nReferences | 1[c] | 4229[d] | 4230 |
| **Total** | | 4329 | 4229 | 8558 |

Concordance rate was 99.99%, calculated by (a+d)/(a+b+c+d).

**Supplementary Table 6 Variant comparison between EBV isolates from paired saliva and NPC tumor samples from the same NPC patient .**

| | | tumor | | Total |
|---|---|---|---|---|
| | | nReference | nVariants | |
| **saliva** | nReference | 7155[a] | 48[b] | 7203 |
| | nVariants | 13[c] | 1136[d] | 1149 |
| **Total** | | 7168 | 1184 | 8352 |

Concordance rate was 99.27%, calculated by (a+d)/(a+b+c+d).

**Supplementary Table 7 Top three associated SNPs in GWAS discovery phase reaching suggestive genome-wide significance ($P < 4.07\times10^{-4}$)**

| POS | Reference/ alternative genotypes | Alt Freq in cases | Alt Freq in controls | *P* _GWAS* | Zscore_GWAS* | LD r-squared with SNP | | Annotation |
|---|---|---|---|---|---|---|---|---|
| | | | | | | 162215 | 162507 | |
| 162215 | C/A | 3.85% | 40.43% | 3.69E-04 | -3.56 | | 0.85 | BALF2, non-synonymous, V700L |
| 162507 | C/T | 2.56% | 42.55% | 9.99E-05 | -3.89 | 0.85 | | BALF2, synonymous |
| 162852 | G/T | 2.56% | 40.43% | 1.84E-04 | -3.74 | 0.90 | 0.95 | BALF2, synonymous |

*Alternative genotypes were tested against reference genotypes in the mixed model in GWAS discovery phase.

**Supplementary Table 8 Fine-mapping for casual SNPs associated with NPC risk in *BALF2* gene region.**

| Position | Reference/ alternative genotypes | Alternative genotype frequency in cases | Alternative genotype frequency in controls | *P* _GWAS* | Zscore_GWAS* | Posterior probability by PAINTOR | Annotation |
|---|---|---|---|---|---|---|---|
| 160804 | C/T | 7.24% | 39.13% | 1.18E-01 | -1.56 | 0.00 | BALF2, synonymous |
| 160827 | G/T | 94.12% | 65.22% | 1.65E-02 | 2.40 | 0.00 | BALF2, synonymous |
| 160941 | G/A | 5.84% | 13.33% | 3.87E-01 | 0.86 | 0.00 | BALF2, synonymous |
| 160971 | T/C | 90.26% | 59.09% | 1.22E-01 | 1.55 | 0.00 | BALF2, synonymous |
| 161036 | T/C | 88.82% | 56.82% | 1.19E-01 | 1.56 | 0.01 | BALF2, non-synonymous, S1093G |
| 162117 | A/G | 93.59% | 65.96% | 2.31E-02 | 2.27 | 0.00 | BALF2, synonymous |
| 162147 | G/A | 3.85% | 38.30% | 2.93E-03 | -2.97 | 0.00 | BALF2, synonymous |
| 162195 | A/C | 92.95% | 65.96% | 3.36E-02 | 2.13 | 0.00 | BALF2, synonymous |
| 162215 | C/A | 3.85% | 40.43% | 3.69E-04 | -3.56 | 0.75 | BALF2, non-synonymous, V700L |
| 162237 | C/G | 93.51% | 65.22% | 2.96E-02 | 2.17 | 0.00 | BALF2, synonymous |
| 162464 | G/A | 93.59% | 61.70% | 4.44E-03 | 2.85 | 0.00 | BALF2, non-synonymous, I613V |
| 162476 | T/C | 93.59% | 61.70% | 4.44E-03 | 2.85 | 0.09 | BALF2, synonymous |
| 162507 | C/T | 2.56% | 42.55% | 9.99E-05 | -3.89 | 0.04 | BALF2, synonymous |
| 162852 | G/T | 2.56% | 40.43% | 1.84E-04 | -3.74 | 0.02 | BALF2, synonymous |
| 163107 | A/C | 93.59% | 65.22% | 1.46E-02 | 2.44 | 0.00 | BALF2, synonymous |
| 163287 | G/A | 93.59% | 63.83% | 4.38E-02 | 2.02 | 0.00 | BALF2, synonymous |
| 163293 | G/A | 3.85% | 40.43% | 1.16E-03 | -3.25 | 0.01 | BALF2, synonymous |
| 163364 | C/T | 88.46% | 48.94% | 5.83E-03 | 2.76 | 0.07 | BALF2, non-synonymous, V317M |
| 163404 | C/A | 94.19% | 63.04% | 2.65E-02 | 2.22 | 0.00 | BALF2, synonymous |
| 163422 | G/T | 93.55% | 63.04% | 4.38E-02 | 2.02 | 0.00 | BALF2, synonymous |
| 163464 | G/A | 87.10% | 52.17% | 3.18E-02 | 2.15 | 0.00 | BALF2, synonymous |
| 163611 | C/T | 94.77% | 65.22% | 1.36E-02 | 2.47 | 0.00 | BALF2, synonymous |
| 163629 | T/C | 94.12% | 65.22% | 3.41E-02 | 2.12 | 0.00 | BALF2, synonymous |

**Supplementary Table 8 Fine-mapping for casual SNPs associated with NPC risk in *BALF2* gene region.**

| Position | Reference/ alternative genotypes | Alternative genotype frequency in cases | Alternative genotype frequency in controls | *P* _GWAS* | Zscore_GWAS* | Posterior probability by PAINTOR | Annotation |
|---|---|---|---|---|---|---|---|
| 163647 | C/T | 8.44% | 15.22% | 9.29E-01 | 0.09 | 0.00 | BALF2, synonymous |
| 163686 | G/A | 3.92% | 42.55% | 2.63E-03 | -3.01 | 0.00 | BALF2, synonymous |
| 163926 | C/T | 83.97% | 51.06% | 1.04E-01 | 1.62 | 0.00 | BALF2, synonymous |
| 163995 | C/T | 5.77% | 29.79% | 1.57E-01 | -1.42 | 0.00 | BALF2, synonymous |
| 164277 | G/T | 7.69% | 17.02% | 5.39E-01 | -0.61 | 0.00 | BALF2, synonymous |

*Alternative genotypes were tested against reference genotypes in the mixed model in GWAS discovery phase.

**Supplemenary Table 9 Basic characteristics of 483 cases and 605 control individuals used for validation phase by age and sex.**

| Variables | Cases | Controls | ***P*** **(chisq)*** |
|---|---|---|---|
| **Sex** | | | 0.481 |
| Male | 364 (75.4%) | 467 (77.2%) | |
| Female | 119 (24.6%) | 138 (22.8%) | |
| **Age** | | | 0.369 |
| Mean | 48.7 | 49.3 | |
| Standard Deviation | 11.3 | 10.6 | |
| < 37 | 17 (3.5%) | 13 (2.1%) | |
| 37-59 | 387 (80.1%) | 487 (80.5%) | |
| >59 | 79 (16.4%) | 105 (17.4%) | |
| **Total** | 483 | 605 | |

* The *p* values were obtained from $\chi 2$ tests

**Supplementary Table 10 EBV haplotypes composed of SNPs 162215, 162476 and 163364 and their odds ratios for NPC risk in 536 and 651 population-based cases and controls .**

| **EBV subtype (162215-162476-163364)** | **536 cases** | | **651 controls** | | **Odds Ratio (95% CI)*** | ***P*** |
|---|---|---|---|---|---|---|
| | **no.** | **%** | **no.** | **%** | | |
| L-L-L (A-T-C) | 22 | 4.10% | 171 | 26.27% | | |
| H-H-H (C-C-T) | 451 | 84.14% | 292 | 44.85% | 12.22 (7.64 - 19.55) | 1.45E-25 |
| H-H-L (C-C-C) | 51 | 9.51% | 118 | 18.13% | 3.31 (1.90 - 5.76) | 2.26E-05 |
| H-L-L (C-T-C) | 9 | 1.68% | 65 | 9.98% | 1.07 (0.47 - 2.45) | 8.69E-01 |
| other subtypes | 3 | 0.56% | 5 | 0.77% | 4.58 (1.02 - 20.62) | 4.72E-02 |

* Odds ratio for individual EBV subtypes were estimated with logistic model by categorizing each subtype as a single variable and adjusted for age, sex and status of single or multiple infection. Subjects with EBV subtype L-L-L, a common low-risk subtype were used as the reference category. H represents high-risk genotypes; L represents low-risk genotypes.

**Supplementary Table 11 Estimation of odds ratios of SNP 162476 and 163364 for NPC risk.**

| | Beta | Standard Error | *P* | Odds ratio | 95% Confidence interval |
|---|---|---|---|---|---|
| SNP162476 | 1.15 | 0.24 | 1.75E-06 | 3.15 | 1.97-5.04 |
| SNP163364 | 1.30 | 0.18 | 3.23E-13 | 3.68 | 2.59-5.22 |

Odds ratios were estimated in 639 cases and 652 controls using a logistic regression model containing SNPs 162476 and 163364. The logistic regression model was adjusted for age and sex.

**Supplementary Table 12 Frequency of high-risk EBV haplotypes in different regions.**

| Geographic origin | Total frequency of high-risk haplotypes C-C-T and C-C-C | | | | |
|---|---|---|---|---|---|
| | NPC cases | | non-NPC samples | | |
| **Africa** | | | | 0.00% | 0/37 |
| **Western countries** | | | | 2.63% | 1/38 |
| **NPC-endmic China** | **93.27%** | **596/639** | | **62.54%** | **419/670** |
| | | | Healthy control | 63.04% | 411/652 |
| | | | Lymphoma | 40.00% | 6/15 |
| | | | Lymphoblastoid cell lir | 66.67% | 2/3 |
| **NPC-non-endemic East Asia** | **55.00%** | **11/20** | | **9.68%** | **3/31** |
| | | | Healthy control | 14.29% | 1/7 |
| | | | Lymphoma | 28.57% | 2/7 |
| | | | Gastric carcinoma | 0.00% | 0/17 |

| Geographic origin | Frequency of high-risk haplotype C-C-T | | | | |
|---|---|---|---|---|---|
| | NPC cases | | non-NPC samples | | |
| **Africa** | | | | **0.00%** | **0/37** |
| **Western countries** | | | | **0.00%** | **0/38** |
| **NPC-endmic China** | **84.35%** | **539/639** | | **44.93%** | **301/670** |
| | | | Healthy control | 44.94% | 293/652 |
| | | | Lymphoma | 40.00% | 6/15 |
| | | | Lymphoblastoid cell lir | 66.67% | 2/3 |
| **NPC-non-endemic East Asia** | **55.00%** | **11/20** | | **6.45%** | **2/31** |
| | | | Healthy control | 14.29% | 1/7 |
| | | | Lymphoma | 14.29% | 1/7 |
| | | | Gastric carcinoma | 0.00% | 0/17 |

**Supplementary Table 13 The percentage of heterozygous variants in 270 EBV genome isolates.**

| Sample ID | nHet/nVar | infection* |
|---|---|---|
| NPCT084 | 2.83% | single |
| NKLT003-2 | 2.90% | single |
| GCT002 | 2.96% | single |
| NPCS023 | 3.11% | single |
| NPCS029 | 3.35% | single |
| NPCT075 | 3.36% | single |
| HS024 | 3.65% | single |
| NPCT049 | 3.81% | single |
| NPCT039 | 3.88% | single |
| NPCS034 | 4.14% | single |
| NPCS033 | 4.20% | single |
| NPCS016 | 4.21% | single |
| NPCS001 | 4.22% | single |
| NPCT035 | 4.22% | single |
| NPCT101 | 4.24% | single |
| NPCT007 | 4.25% | single |
| NPCS003-2 | 4.27% | single |
| NPCT011 | 4.27% | single |
| NPCS045 | 4.30% | single |
| NPCT052 | 4.31% | single |
| NPCT037 | 4.34% | single |
| NPCT056 | 4.34% | single |
| NPCS014 | 4.36% | single |
| HS029 | 4.38% | single |
| NPCT056M | 4.39% | single |
| NNPCT004 | 4.46% | single |
| NPCT055 | 4.47% | single |
| NPCT112 | 4.47% | single |
| HLT010 | 4.49% | single |
| NPCS011 | 4.51% | single |
| HS054 | 4.54% | single |
| HLT005 | 4.55% | single |
| NPCS022 | 4.55% | single |
| NPCT020-2 | 4.56% | single |
| NPCT058 | 4.57% | single |
| NPCS028 | 4.61% | single |
| HS014 | 4.67% | single |
| BLT001 | 4.69% | single |
| NPCT058M | 4.70% | single |
| NPCT085 | 4.72% | single |
| NPCS026 | 4.73% | single |
| NKLT004 | 4.77% | single |
| NPCT028-2 | 4.80% | single |
| NPCT113 | 4.81% | single |
| NPCS051 | 4.83% | single |
| NPCT069 | 4.84% | single |
| NPCS054 | 4.87% | single |
| NPCT029 | 4.88% | single |
| NPCT036 | 4.89% | single |

**Supplementary Table 13 The percentage of heterozygous variants in 270 EBV genome isolates.**

| Sample ID | nHet/nVar | infection* |
|---|---|---|
| NPCT093 | 4.89% | single |
| NPCT105 | 4.90% | single |
| NNPCT002 | 4.90% | single |
| NPCT067 | 4.95% | single |
| NPCT063 | 4.97% | single |
| NPCT054 | 4.98% | single |
| NPCT012 | 5.00% | single |
| NKLT002 | 5.01% | single |
| NPCT038 | 5.01% | single |
| NPCT017 | 5.06% | single |
| NPCT104 | 5.07% | single |
| NPCT078 | 5.08% | single |
| NPCS017 | 5.09% | single |
| NPCT006 | 5.11% | single |
| NPCS012 | 5.14% | single |
| NPCT018 | 5.15% | single |
| NPCT004 | 5.15% | single |
| NPCT088 | 5.19% | single |
| NPCT076 | 5.19% | single |
| NPCS048 | 5.20% | single |
| NPCT074S | 5.22% | single |
| NPCT023 | 5.22% | single |
| NPCS047 | 5.23% | single |
| NPCT073 | 5.23% | single |
| NPCT009 | 5.23% | single |
| NPCS002 | 5.24% | single |
| NPCS019 | 5.24% | single |
| NPCS038 | 5.24% | single |
| HS015 | 5.26% | single |
| NPCT080 | 5.30% | single |
| NPCT055M | 5.30% | single |
| NPCT114 | 5.30% | single |
| NPCS009 | 5.32% | single |
| NPCT100 | 5.32% | single |
| HLT007 | 5.33% | single |
| HLT001 | 5.33% | single |
| NPCT002 | 5.35% | single |
| NPCS010 | 5.35% | single |
| NPCT094 | 5.36% | single |
| NPCS042 | 5.40% | single |
| NPCT081 | 5.40% | single |
| NPCS018 | 5.42% | single |
| NPCS049 | 5.45% | single |
| HLT006 | 5.45% | single |
| NPCT066 | 5.47% | single |
| NPCT099 | 5.49% | single |
| NPCS024 | 5.49% | single |
| NPCS025 | 5.50% | single |
| NPCT070 | 5.53% | single |

**Supplementary Table 13 The percentage of heterozygous variants in 270 EBV genome isolates.**

| Sample ID | nHet/nVar | infection* |
|---|---|---|
| NPCT062 | 5.54% | single |
| NPCP001 | 5.56% | single |
| NPCS021 | 5.57% | single |
| NPCT057 | 5.57% | single |
| NPCS027 | 5.58% | single |
| NPCT108 | 5.59% | single |
| HS053 | 5.62% | single |
| HS027 | 5.64% | single |
| NPCT010 | 5.64% | single |
| NPCT031 | 5.65% | single |
| NPCT083 | 5.66% | single |
| NPCS039 | 5.66% | single |
| HS011 | 5.67% | single |
| NPCS031 | 5.67% | single |
| NPCT032 | 5.68% | single |
| NPCT008 | 5.69% | single |
| NPCT043 | 5.70% | single |
| NPCT059 | 5.71% | single |
| NPCT050 | 5.71% | single |
| NPCT014 | 5.72% | single |
| NPCT027 | 5.74% | single |
| NKLT007 | 5.76% | single |
| NNPCT001 | 5.77% | single |
| NPCS008 | 5.77% | single |
| NPCS035 | 5.77% | single |
| HS020 | 5.78% | single |
| NPCT077 | 5.79% | single |
| HS013 | 5.81% | single |
| NPCS044 | 5.82% | single |
| NPCS050 | 5.82% | single |
| HS052 | 5.83% | single |
| NPCT005 | 5.86% | single |
| NPCT045 | 5.87% | single |
| NPCT003 | 5.88% | single |
| NPCS046 | 5.90% | single |
| HS037 | 5.94% | single |
| NPCT053 | 5.96% | single |
| NPCT064 | 5.98% | single |
| NPCT013 | 5.98% | single |
| NPCT098 | 6.01% | single |
| NPCT054M | 6.03% | single |
| NPCT042 | 6.05% | single |
| NPCT060 | 6.07% | single |
| NPCT086 | 6.08% | single |
| HS038 | 6.11% | single |
| NPCT065 | 6.11% | single |
| NPCT001 | 6.12% | single |
| HS012 | 6.14% | single |
| HS019 | 6.14% | single |

**Supplementary Table 13 The percentage of heterozygous variants in 270 EBV genome isolates.**

| Sample ID | nHet/nVar | infection* |
|---|---|---|
| NPCS005 | 6.14% | single |
| NPCS006 | 6.15% | single |
| NPCT074 | 6.17% | single |
| NPCS040 | 6.18% | single |
| NPCT102 | 6.27% | single |
| NPCS007 | 6.29% | single |
| NPCT047 | 6.29% | single |
| NPCS030 | 6.30% | single |
| NPCT057M | 6.31% | single |
| HS023 | 6.31% | single |
| NPCT087 | 6.34% | single |
| NPCT103 | 6.39% | single |
| NPCT089 | 6.40% | single |
| HS009 | 6.40% | single |
| BLT002 | 6.41% | single |
| NPCT106 | 6.43% | single |
| NPCT071 | 6.44% | single |
| NPCT033 | 6.44% | single |
| HS041 | 6.48% | single |
| HS008 | 6.52% | single |
| NPCT048 | 6.53% | single |
| NPCT115 | 6.55% | single |
| NPCT040 | 6.57% | single |
| NPCT024 | 6.60% | single |
| HS016 | 6.61% | single |
| NPCT090 | 6.65% | single |
| NPCT061 | 6.67% | single |
| NPCT109 | 6.69% | single |
| C666 | 6.70% | single |
| NPCT068 | 6.74% | single |
| NPCT041 | 6.76% | single |
| NPCS052 | 6.82% | single |
| HS048 | 6.82% | single |
| NPCT091 | 6.82% | single |
| NPCT025 | 6.84% | single |
| HS003 | 6.84% | single |
| GCT003 | 6.85% | single |
| NKLT006 | 6.86% | single |
| NPCT072 | 6.92% | single |
| NPCT019 | 6.92% | single |
| NPCT110 | 6.97% | single |
| GCT012 | 7.04% | single |
| NPCT096 | 7.09% | single |
| NPCT092 | 7.10% | single |
| HS039 | 7.14% | single |
| GCT001 | 7.17% | single |
| HS035 | 7.21% | single |
| NPCT015 | 7.30% | single |
| GCT011 | 7.35% | single |

**Supplementary Table 13 The percentage of heterozygous variants in 270 EBV genome isolates.**

| Sample ID | nHet/nVar | infection* |
|---|---|---|
| NPCT082 | 7.38% | single |
| HLT011 | 7.48% | single |
| HLT002 | 7.54% | single |
| GCT004 | 7.55% | single |
| NPCT022 | 7.58% | single |
| NPCT107 | 7.62% | single |
| HS057 | 7.63% | single |
| HS001 | 7.83% | single |
| HS033 | 7.85% | single |
| HS051 | 7.89% | single |
| NPCT046 | 7.90% | single |
| GCT006 | 7.93% | single |
| HS036 | 7.96% | single |
| NPCS013 | 7.99% | single |
| NHS002 | 8.11% | single |
| GCT007 | 8.15% | single |
| GCT014 | 8.35% | single |
| NPCT111 | 8.46% | single |
| NNPCT005 | 8.51% | single |
| HS050 | 8.67% | single |
| GCT009 | 8.71% | single |
| NNPCT003 | 8.91% | single |
| HS034 | 9.04% | single |
| GCT010 | 9.19% | single |
| HS032 | 9.34% | single |
| HS045 | 9.35% | single |
| HS025 | 9.47% | single |
| HS021 | 9.48% | single |
| HS018 | 9.56% | single |
| GCT005 | 9.61% | single |
| GCT013 | 9.63% | single |
| NHS004 | 9.85% | single |
| HS007 | 9.86% | single |
| NPCT021 | 9.94% | single |
| GCT015 | 10.89% | |
| GCT008 | 10.96% | |
| NPCT116 | 11.03% | |
| HS010 | 11.30% | |
| GCT016 | 12.17% | |
| NPCS041 | 14.20% | |
| HS056 | 14.21% | |
| HS026 | 14.72% | |
| NPCT016 | 14.85% | |
| HS006 | 15.80% | |
| NPCS037 | 17.06% | |
| NPCT051 | 17.08% | |
| NPCS015 | 18.16% | |
| NPCT097 | 19.85% | |
| NPCT095 | 20.29% | |

**Supplementary Table 13 The percentage of heterozygous variants in 270 EBV genome isolates.**

| Sample ID | nHet/nVar | infection* |
|---|---|---|
| HS031 | 24.10% | |
| NPCT026 | 28.59% | |
| HS046 | 29.25% | |
| NNPCT006 | 30.30% | |
| NKLT001 | 33.80% | |
| NPCS036 | 38.13% | |
| HS005 | 38.65% | |
| NPCS032 | 39.21% | |
| HS055 | 41.48% | |
| NPCS004 | 42.47% | |
| HS040 | 42.71% | |
| NKLT005 | 43.92% | |
| HLT009 | 44.35% | |
| NPCT079 | 50.75% | |
| HS028 | 57.93% | |
| HS017 | 60.29% | |
| HLT012 | 60.61% | |
| NHS001 | 61.11% | |
| HLT004 | 62.69% | |
| NPCS043 | 63.79% | |
| HS030 | 64.65% | |
| HS022 | 66.73% | |
| NHS003 | 68.66% | |
| HLT003 | 68.84% | |
| NPCS055 | 70.60% | |

*A threshold of heterozygous variant proportion was set at 10.7% for single EBV infection.

nHet/nVar, number of heterozygous variants / number of total variants

**Supplementary Table 14 The association of EBV haplotypes with EBV DNA abundance in saliva of 533 cases and 651 controls.**

| | ***Beta* ( 95% CI)** | ***P**** |
|---|---|---|
| **H-H-H *versus* other haplotypes** | -0.10 (-0.62, 0.43) | 0.7224 |
| **control *versus* case** | -2.13 (-2.66, -1.60) | 6.50E-15 |
| **Female *versus* male** | 1.31 (0.75, 1.87) | 4.18E-06 |
| **Age** | -0.01 (-0.03, 0.01) | 2.91E-01 |

*The beta and *P* values were obtained from multiple linear regression against cycle-of-threshold value measured by qPCR of EBV DNA with EBV haplotypes, single-multiple infection status, case-control status, sex and age. Fold change of EBV DNA abundance was assessed by 2^(-beta). H represents high-risk genotype. H-H-H represents the high-risk haplotype carrying risk genotypes of SNPs 162215, 162476 and 163364.

**Supplementary Table 15 Estimation of the proportion of NPC population risk attributable to high-risk EBV haplotypes in 536 and 651 population-based cases and controls.**

| High-risk haplotype | Population attributable risk fraction | 95% confidence interval |
|---|---|---|
| C-C-T | 70.90% | 67.41%-74.40% |
| C-C-T and C-C-C | 82.97% | 79.33%-86.60% |

The attributable fraction of risk and 95% confidence interval were estimated in a logistic regression model with adjustment for age, sex and status of single- or multiple-infection. For details, see methods.

### Supplementary Note

#### Patient recruitment in the population-based case-control study

The study design of the population-based case-control study has been previously described in detail (14) [1]. To accommodate available resources, the present analysis was confined to NPC cases and controls enrolled from Zhaoqing County between January 2010 and October 2014, using the following eligibility criteria: (i) histological confirmation of NPC, (ii) age less than 80 years, (iii) no treatment for NPC, and (iv) residence in Zhaoqing city. Among 1,306 eligible NPC patients recruited into the study through a rapid case ascertainment system involving a network of physicians, 1,043 (79.9%) had available saliva samples that were sequenced or genotyped. Through random selection from the total population registry in Zhaoqing County, 1,151 population control subjects without any history of malignancy (84.3% of 1,365 eligible controls enrolled with frequency matching to cases by 5-year age and sex) had available saliva samples that were sequenced or genotyped.

#### Evaluation of EBV DNA abundance in saliva and its correlation with genotyping success rate

EBV DNA abundance in saliva samples from the 1043 cases and 1151 controls in Zhaoqing County was measured in triplicate for each sample by fluorescence quantitative PCR (qPCR) using a DNA fragment of *BALF5*. The relative DNA abundance was calculated by the $2^{-\Delta Ct}$ method where Ct is the cycle of threshold and deduced from qPCR standard curve.

As EBV in buccal mucosa undergoes periodic lytic cycle[2-4], in a large proportion of NPC cases and healthy controls EBV DNA abundance was found to be quite low and did not allow for EBV WGS or successful genotyping. From the cases and controls, 53 and 46 saliva samples with Ct value < 30 were used for EBV WGS to ensure the success of WGS. The remaining 990 cases and 1105 controls were used for genotyping of the three GWAS candidate markers for validation. Saliva EBV DNA amount is highly correlated with genotyping success rate (**Supplementary Figs. 12**). When Ct value was < 30, genotyping success rate reached 87%. However, this rate dropped significantly as Ct value increased (**Supplementary Table 16**). Therefore, all three SNP genotypes were only obtained in saliva samples from 483 cases (48.79%) and 605 controls (54.75%). EBV genotyping success rate in saliva from controls was slightly higher than from cases (**Supplementary Table 17**). Consistently, in these saliva samples, we found that EBV DNA level from cases was significantly lower than from controls (**Fig. 3a**).

**Supplementary Table 16 The proportion of samples successfully genotyped for 0-3 SNPs out of the total samples with EBV DNA Ct values as indicated. (ca: case, co: control; No.: number)**

| | Ct <28 | Ct < 30 | Ct < 33 | Ct <35 | Total |
|---|---|---|---|---|---|
| successfully genotyped SNP No. | 118ca 289co | 207ca 500co | 452ca 775co | 655ca 901co | 990ca 1105co |
| 0 | 3.69% | 5.94% | 11.65% | 17.35% | 29.02% |
| 1 | 2.21% | 2.97% | 4.56% | 8.10% | 9.74% |
| 2 | 3.19% | 4.10% | 8.15% | 9.45% | 9.31% |
| 3 | 90.91% | 86.99% | 75.63% | 65.10% | 51.93% |

**Supplementary Table 17 The summary of cases and controls successfully genotyped for 0-3 SNPs and the average Ct values in each category as indicated. (No.: number)**

| | case (990) | | | control (1105) | | |
|---|---|---|---|---|---|---|
| successfully genotyped SNP No. | No. | percent | average Ct | No. | percent | average Ct |
| 0 | 291 | 29.39% | 36.17 | 317 | 28.69% | 33.97 |
| 1 | 118 | 11.92% | 35.33 | 86 | 7.78% | 32.20 |
| 2 | 98 | 9.90% | 34.26 | 97 | 8.78% | 31.41 |
| 3 | 483 | 48.79% | 30.62 | 605 | 54.75% | 28.38 |

#### EBV whole-genome sequencing, variant calling and filtering

**Targeted EBV WGS**. Genomic DNA extracted from tumor, saliva, plasma and cell line was subject to hybrid capture by an EBV-targeting single-stranded DNA probe developed by MyGenostics. Sequencing libraries were constructed by shearing genomic DNA into 150-200bp DNA fragments, DNA purification, end blunting, and adaptor ligation according to instructions from Illumina. The library concentrations were evaluated by Bioanalyzer 2100 (Agilent Technologies, Santa Clara, CA, USA). EBV DNA was captured from genomic DNA following the MyGenostics GenCap

Target Enrichment Protocol (GenCap Enrichment, MyGenostics, USA). Libraries were hybridized with EBV probes at 65 ℃ for 24 h and then washed to remove uncaptured DNA. The eluted DNA fragments were amplified by 18 PCR cycles to generate libraries for sequencing. Libraries were quantified and subjected to paired-end sequencing on Illumina HiSeq 2000 sequencer according to the manufacturer's instructions (Illumina Inc., San Diego, CA, USA).

**Read Mapping**. Quality assessment was conducted on the raw reads using Trim-galore to remove adaptor sequences and reads that were of low quality. High-quality reads were aligned to wild-type EBV genome (NC_007605.1) as reference using Burrows-Wheeler Aligner (BWA, version 0.7.5a)[5]. Alignments were converted from sequence alignment map format to sorted, indexed binary alignment map (BAM) files[6]. The Picard tool was used to remove duplicate reads. The depth and coverage of each sample were calculated. The average sequencing depth for EBV genomes was 1282 (range, 32 to 6629), and on average 95.28% of the genome were covered with at least 10× reads (**Supplementary Fig. 1**).

**Variant calling and filtering**. GATK software tools (version 3.2-2) were used for improvement of alignments and genotype calling following GATK's Best Practice[7]. Briefly, BAM files were realigned with the GATK IndelRealigner. The base quality of the mapped reads was recalibrated by GATK base quality recalibration tool BQSR. As BQSR requires genuine SNP database to do recalibration, we generated our own high-quality EBV SNP database. Raw variants were first called by GATK UnifiedGenotyper and HaplotypeCaller separately against the WT EBV genome (NC_007605.1). Common variants identified by the two callers were selected and filtered to generate database SNP for BQSR. Analysis-ready reads were prepared after three cycles of BQSR when before-after BQSR plots converged and the recalibration reached saturation. Subsequently, variants were called by GATK UnifiedGenotyper using analysis-ready reads. As EBV has a small genome and a small number of variants, hard-filtering was recommended by GATK developer to exclude the low-quality variant due to (i) low variant confidence, (ii) low read-mapping quality, (iii) strand bias (the variation being seen on only the forward or only the reverse strand) in the reads and (iv) reads that were aligned to multiple positions in EBV genome. In particular, SNPs and

INDELs were filtered separately by GATK VariantFiltration with the parameter "MQ0 >= 4 && ((MQ0 / (1.0 * DP)) > 0.1)", "QUAL < 50.0", "QD < 2.0", "MQ < 40.0", "FS > 250.0" for SNPs and "MQ0 >= 4 && ((MQ0 / (1.0 * DP)) > 0.1)", "QUAL < 50.0", "FS > 200.0" for INDELs. We identified an initial set of high-quality 8469 variants from 269 samples and the C666-1 cell line.

In order to avoid inaccurate calling, we further filtered out variants that has (i) low coverage support (depth < 10×), (ii) in repetitive elements (NCBI annotation of reference NC_007605.1), (iii) within 5 bp of an indel, and 7,962 variants were retained for subsequent EBV phylogenetic, principal component and association analyses. Metrics including the number of filtered SNP counts, concordance of variants among samples, and ratio of heterozygous to single variants were evaluated using GATK VariantEval. The annotation of variants was performed and summarized by SNPEff[8] according to the annotation of NC_007605.1 (NCBI annotation, NOV 2013).

#### Determining single *versus* multiple EBV infections.

As EBV genome is stable and intra-host mutation rate is often low[9], heterozygous variants caused by intra-host mutation occur at low frequency. We sequenced EBV genomes in quadruple replicates from the NPC cell line C666-1[10]. EBV genomes in cell lines and EBV-associated tumors usually undergo clonal expansion[11-13], and the heterozygous variants come from low-level genomic evolution during cell proliferation over decades. The proportions of heterozygous variants ranged from 6.7% to 9.5% discovered by quadruple replicates of C666-1 EBV whole-genome sequencing. By contrast, the EBV isolates with multiple infections tend to have higher number of heterozygous variants. Therefore, we first extracted the empirical distribution of the percentage of heterozygous variants across all the samples. By fitting two different curves to the lower (< 8%) and higher quantiles (> 15%) of the empirical distribution, we identify a reflection point separating the two tails of the distribution (**Supplementary Fig. 4**). The reflection point (10.7%) was then used to define a threshold of the number of heterozygous SNPs. The 230 samples with the number of heterozygous SNPs lower than the threshold were identified as single infection samples for subsequent phylogenetic analysis and principal component analysis.

In 483 NPC cases and 605 controls from Zhaoqing case-control study, we genotyped all three EBV GWAS candidate markers. In saliva from 464 cases (96.07%) and 570 controls (94.25%) we detected EBV infection with single haplotypes where all three markers were homozygous, whereas saliva from only 19 cases (3.93%) and 35 controls (5.75%) contained EBV infection with multiple EBV haplotypes with heterozygous markers. The multiple-infection EBV haplotypes were deduced by phasing using Beagle 4.1[14]. In the association study, we included cases and controls carrying both single infection and multiple infection with EBV haplotypes. As we adjusted for multiple infection which only accounted for a small proportion of cases and controls in the association study, our association results would not be confounded by multiple infection.

1 Ye, W. *et al.* Development of a population-based cancer case-control study in southern china. *Oncotarget* **8**, 87073-87085, doi:10.18632/oncotarget.19692 (2017).

2 Kieff, E. D. & Rickinson, A. B. in *Fields' virology* Vol. 68A (eds D.M. Knipe & P.M. Howley) 2603-2654 (Lippincott Williams & Wilkins, Wolters Kluwer, 2007).

3 Borza, C. M. & Hutt-Fletcher, L. M. Alternate replication in B cells and epithelial cells switches tropism of Epstein-Barr virus. *Nat Med* **8**, 594-599, doi:10.1038/nm0602-594 (2002).

4 Frangou, P., Buettner, M. & Niedobitek, G. Epstein-Barr virus (EBV) infection in epithelial cells in vivo: rare detection of EBV replication in tongue mucosa but not in salivary glands. *J Infect Dis* **191**, 238-242, doi:10.1086/426823 (2005).

5 Li, H. & Durbin, R. Fast and accurate short read alignment with Burrows-Wheeler transform. *Bioinformatics* **25**, 1754-1760, doi:10.1093/bioinformatics/btp324 (2009).

6 Li, H. *et al.* The Sequence Alignment/Map format and SAMtools. *Bioinformatics* **25**, 2078-2079, doi:10.1093/bioinformatics/btp352 (2009).

7 DePristo, M. A. *et al.* A framework for variation discovery and genotyping using next-generation DNA sequencing data. *Nature genetics* **43**, 491-498,

doi:10.1038/ng.806 (2011).

8 Cingolani, P. *et al.* A program for annotating and predicting the effects of single nucleotide polymorphisms, SnpEff: SNPs in the genome of Drosophila melanogaster strain w1118; iso-2; iso-3. *Fly (Austin)* **6**, 80-92, doi:10.4161/fly.19695 (2012).

9 Weiss, E. R. *et al.* Early Epstein-Barr Virus Genomic Diversity and Convergence toward the B95.8 Genome in Primary Infection. *J Virol* **92**, doi:10.1128/JVI.01466-17 (2018).

10 Cheung, S. T. *et al.* Nasopharyngeal carcinoma cell line (C666-1) consistently harbouring Epstein-Barr virus. *Int J Cancer* **83**, 121-126 (1999).

11 Raab-Traub, N. & Flynn, K. The structure of the termini of the Epstein-Barr virus as a marker of clonal cellular proliferation. *Cell* **47**, 883-889 (1986).

12 Pathmanathan, R., Prasad, U., Sadler, R., Flynn, K. & Raab-Traub, N. Clonal proliferations of cells infected with Epstein-Barr virus in preinvasive lesions related to nasopharyngeal carcinoma. *N Engl J Med* **333**, 693-698, doi:10.1056/NEJM199509143331103 (1995).

13 Neri, A. *et al.* Epstein-Barr virus infection precedes clonal expansion in Burkitt's and acquired immunodeficiency syndrome-associated lymphoma. *Blood* **77**, 1092-1095 (1991).

14 Browning, S. R. & Browning, B. L. Rapid and accurate haplotype phasing and missing-data inference for whole-genome association studies by use of localized haplotype clustering. *American journal of human genetics* **81**, 1084-1097, doi:10.1086/521987 (2007).
